# Development of coarse-grained model for a minimal stratum corneum lipid mixture

**DOI:** 10.1101/2021.10.04.463115

**Authors:** Parashara Shamaprasad, Timothy C. Moore, Donna Xia, Christopher R. Iacovella, Annette L. Bunge, Clare McCabe

## Abstract

Molecular dynamics simulations of mixtures of the ceramide N-(tetracosanoyl)-sphingosine (NS), cholesterol, and a free fatty acid are performed to gain a molecular-level understanding of the structure of the lipids found in the stratum corneum layer of skin. A new coarse-grained model for cholesterol, developed using the multistate iterative Boltzmann inversion method, is compatible with previously developed coarse-grained forcefields for ceramide NS, free fatty acid, and water, and validated against atomistic simulations of these lipids using the CHARMM force field. Self-assembly simulations of multilayer structures using these coarse-grained force fields are performed, revealing that a large fraction of the ceramides adopt extended conformations, which cannot occur in the bilayer structures typically studied using simulation. Cholesterol fluidizes the membrane by promoting packing defects and it is observed that an increase in cholesterol content reduces the bilayer height, due to an increase in interdigitation of the C24 lipid tails, consistent with experimental observations. Through the use of a simple reverse-mapping procedure, a self-assembled coarse-grained multilayer system is used to construct an equivalent structure with atomistic resolution. Simulations of this atomistic structure are found to closely agree with experimentally derived neutron scattering length density profiles. Significant interlayer hydrogen bonding is observed in the inner layers of the atomistic multilayer structure that are not found in the outer layers in contact with water or in equivalent bilayer structures. These results identify several significant differences in the structure and hydrogen bonding of multilayer structures as compared to the more commonly studied bilayer systems, and, as such, highlight the importance of simulating multilayer structures for more accurate comparisons with experiment. These results also provide validation of the efficacy of the coarse-grained forcefields and the framework for multiscale simulation.

## Introduction

The visible, outermost skin layer, called the stratum corneum (SC), is essential for skin to act as an effective barrier for water loss from the body and invasion of toxic chemicals and infectious organisms. The SC is organized into a brick-and-mortar like arrangement in which corneocytes (the bricks) are surrounded by a dense, lamellar-structured lipid matrix (the mortar). The lipid matrix is a complex mixture of ceramides (CERs), cholesterol (CHOL), and free fatty acids (FFAs) with a distribution of chain lengths. At least 21 unique CERs, which consists a fatty acid chain connected by an amide bond to a sphingoid base, have been identified in human SC and a distribution of FFA chain lengths.^1–4^ The structure and composition of this lipid matrix are crucial for a properly functioning skin barrier.^5^ X-ray scattering studies have been used to probe the lamellar organization of the lipids in the SC, finding that a ∼13 nm long periodicity phase (LPP) and a ∼6 nm short-periodicity phase (SPP) coexist in healthy SC.^6,7^ Skin diseases that impair the barrier function of the skin, such as psoriasis and atopic dermatitis, affect the expression of lipid synthases, resulting in altered lipid compositions in the SC.^8^ Furthermore, x-ray scattering studies have found that SC from psoriasis and atopic dermatitis patients exhibits abnormal lamellar arrangements compared to healthy skin, which further compromises the skin barrier.^8^ While the SC lipid composition is thus known to influence SC structure and barrier function, the complexity of the SC lipid mixture poses challenges for study in terms of elucidating the mechanism behind the composition-structure-barrier function relationship.^1^

In order to better understand the cause-and-effect relationships that exist between lipid composition and structure/barrier function, model systems that have well defined lipid compositions have more recently been considered.^9–13^ In particular, several X-ray and neutron scattering studies have examined mixtures that form an SPP-like phase without a co-existing LPP to elucidate the structure of the SPP in isolation^14,15^ and also to examine structural changes in the SPP with variations in lipid composition.^16–18^ Although not direct models of native SC, these simplified mixtures can provide important mechanistic insights into the structural properties of SC lipids and the role of individual lipids on these properties. In addition to well-defined compositions, the use of synthetic SC lipid mixtures also allows for selective lipid deuteration, which can be used to determine lipid localization using neutron scattering or Fourier transform infrared spectroscopy (FTIR). For example, Groen et al. found that the fatty acid tails of the CERs interdigitate in the SPP using selectively deuterated mixtures of CERs NS, NP, AP, AS, CHOL and FFA.^15^ Schmitt et al. found that varying ratios of CER NP C_18_ to CER AP C_18_ in mixtures with 1:0.7:1 molar ratios of CER, CHOL, and FFA C_24_ induced changes in the degree of interdigitation of lipid tails in the SPP.^16^ The work of Engberg et al. focused on a simpler model of SC lipid mixtures containing only CER NS C_24_, CHOL, and FFA C_24_ in 1:1:1 and 1:0.45:1 and found via x-ray scattering that an SPP forms, albeit with a smaller repeat distance of 5.38 nm.^19^ They also found that CHOL is phase separated in the 1:1:1 mixture but not in the 1:0.45:1 mixture, suggesting that the solubility limit of CHOL in the SPP is near the 1:0.45:1 ratio.^19^ Similarly, Mojumdar, et al. determined that the solubility limit of CHOL before phase separation was near a 1:0.5:1 ratio of CER:CHOL:FFA, in SPP systems containing a mixture of CERs NS, NP, AS, and AP, as well as 7 different FFA chain lengths ranging from C_16_-C_26_.^20^ Using x-ray scattering, they also found that the repeat distance of the SPP-like phase increases as the CHOL fraction is reduced below the solubility limit.^20^

Although experiments can reveal changes in lamellar phase behavior using axial neutron scattering length density profiles from X-ray scattering studies, these studies are costly and do not provide a direct and clear representation of the lateral and axial lipid arrangement. To address these issues, researchers have turned to molecular dynamics (MD) simulations, which provide an atomic/molecular-scale description of the structure and arrangement of lipids within the lamellar. In addition, MD simulations allow for explicit control over the system, and thus can be used to examine different aspects of parameter space, such as the effect of lipid type, chain length, and composition. For example, Das et al.^21^ examined CER NS C_24_:CHOL:FFA C_24_ bilayers and varied the molar ratio of each lipid systematically, finding increased CHOL fraction reduced nematic order. They also observed a smaller bilayer thickness in systems with higher CHOL content, which was attributed to longer CER acyl and FFA tails bending to fill the space around the shorter CHOL molecules.^21^ In another study, Moore et al.^22^ conducted MD simulations of bilayer structures that screened over 4 ratios of CER NS, CHOL, and FFA C_24_ and 5 ratios of CER NS C_24_ to CER NS C_16_ to determine the effect of composition changes on structural properties. An increase in CHOL content of CER NS C_24_:CHOL bilayers was found to result in an increase in density of the tail region, decrease in bilayer thickness, and an increase in interdigitation.^22^ In addition, increased CHOL content resulted in fewer hydrogen bonds between lipids, which may lead to a less stable bilayer interface.^22^ Based on CHOL-CHOL coordination numbers, it was also determined that CHOL does not preferentially neighbor specific lipid tails, whereas FFA molecules prefer to reside by FFA tails and CER fatty acid chains, suggested to be driven by van der Waals attractions between the long alkane chains.^22^ In a similar study, Wang and Klauda explored structural changes in equimolar CER NS:CHOL:FFA C_24_ mixtures using either CER NS C_16_ or CER NS C_24_ at four temperatures.^23^ It was observed that increasing temperature disrupts chain order and packing, as would be expected, shown by decreases in the deuterium order parameter and hydrogen bonding, and an increase in area per lipid.^23^ However, it was also found that hydrogen bonding did not decrease uniformly for all lipids as temperature was increased.^23^ Based on the relative magnitudes of the decrease in the number of hydrogen bonds over the temperature range of 32 to 65 ºC, CER-CER hydrogen bonds appeared to be more resilient to increases in temperatures as compared to those involving FFA or CHOL.^23^ In another study, Del Regno and Notman^24^ simulated model membranes comprised of CER NS C_24_, CHOL, and FFA C_24_ in equimolar and 2:2:1 ratios and observed the formation of CHOL-rich regions in bilayer simulations. In addition, they found that the free energy barrier of water traversing the CHOL rich regions was lower than in CHOL-poor regions, suggesting that water more favorably permeates through CHOL-rich regions in the SC.^24^

While molecular simulations are a powerful tool for gaining a structural understanding of SC lipid membranes, the high computational cost of atomistic simulations limits the size and timescales that can be reasonably studied. In addition, the gel-phase nature of the SC lamellae results in very low lipid mobility, requiring much longer simulation times to reach equilibrium than in more commonly studied fluid-phase membranes. Two major consequences have arisen as a result of these computational limitations. First, due to the long microsecond time scales required to complete self-assembly, atomistic simulations thus far have largely used preassembled membrane configurations, rather than self-assembling membranes from isotropic mixtures; unless careful equilibration procedures are used, the final configuration of the simulation may not be able to adequately decorrelate from the pre-assembled initial configuration, thus potentially biasing the results.^22,25^ The second consequence of computational limitations is that atomistic simulations have, to date, focused almost exclusively on bilayer structures, where lipid conformations are strongly dictated by the lipid-water interface, whereas, lipids in the SC are largely dehydrated and have minimal contact with a water interface.^15^ Furthermore, in bilayer simulations, CER molecules are restricted to the hairpin conformation (Fig 1a), while X-ray scattering studies of model SPP membranes suggest that a mixture of hairpin and extended conformation are present in the multilamellar systems seen in the SC.^15,18^ While pre-assembled multilamellar structures could certainly be simulated using atomistic models, the transitions between hairpin and extended conformations, from one leaflet to another, or from well-mixed systems to lipid clusters would not likely be observed over computationally feasible timescales. That is, lipids would likely remain in which ever conformation or in-plane morphological arrangement for which they were initialized, potentially biasing the results.^22^ Rather than using pre-assembled configurations, self-assembly provides a possible route to allow membrane configurations to arise naturally. However, such studies are likely not computationally attainable with atomistic models at the current time, given the long timescales that would be required for self-assembly to be observed in mixed lipid systems, as well as the need for larger simulation sizes in order to capture multilamellar structures, which significantly increases the computational cost.^26^ In order to overcome limitations in computational efficiency, coarse-grained (CG) simulation approaches, where groups of atoms are treated as single interaction sites (CG beads), have been considered.^27–30^

**Figure 1:**
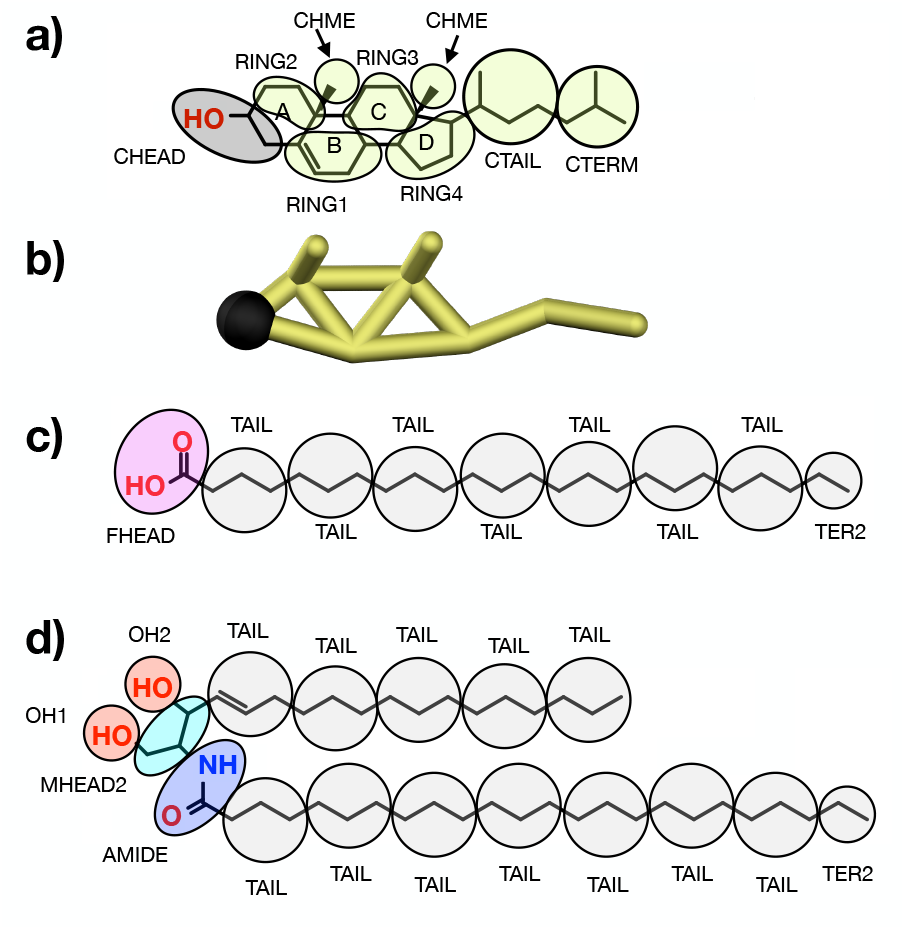
(a) CG mapping scheme for CHOL where the A, B, C, and D rings of the skeletal structure are labeled, and (b) CG representation of CHOL. CG mapping scheme for (c) FFA C_24_, and (d) CER NS C_24_. In all cases, CG beads are spherically symmetric. CG mappings are colored to be consistent with simulation renderings in this work: CHEAD is black, CBODY (i.e., RING1-4, CHME, CTAIL, CTERM) yellow, FHEAD purple, TAIL and TER2 gray, OH1 and OH2 red, MHEAD2 cyan, and AMIDE blue.

Simulations using CG models allow longer timescales and larger system sizes to be studied than those using atomistic models, due to a reduction in the number of interactions that must be computed at each timestep and a softening of the interactions, allowing for larger integration timesteps. While numerous CG studies have been performed of phospholipids, similar studies for SC lipids and CERs in particular are in their infancy. The popular CG MARTINI model has been applied to a few SC lipids (CER NS^27,31–34^, CER AP^28^, CER NP,^35^ CHOL^36,37^, and FFA^36^). However, the generic nature of the MARTINI force field parameters, which circumvents the need for extensive optimization, also limits MARTINI*e*s ability to accurately reproduce experimental and atomistic SC lipid membrane structures. To address this limitation, Hadley and M^c^Cabe developed the first CG model specifically optimized for SC lipids. Using the iterative Boltzmann inversion method (IBI), they optimized a CG forcefield to accurately reproduce the structure of atomistically simulated systems of pure FFA and CHOL as well as mixtures thereof. MacDermaid et al. have also developed CG models for CER NS, CER EOS, CHOL, and FFA based on the Klein model, in which generic functional groups are parameterized by fitting to experimental density and surface tension data for small molecules.

More recently, M^c^Cabe and co-workers have developed CG force fields for CER NS,^38^ FFA,^38^ water,^39^ and the corresponding cross interactions,^30^ using the multistate iterative Boltzmann inversion (MS-IBI) method, combined with simulated wetting, which removes the limitations of state transferability of the IBI method.^40^ These force fields have been shown to provide close agreement with their atomistic targets in terms of both bulk fluid and ordered structural properties, while also capturing the correct balance between hydrophobic and hydrophilic interactions necessary for self-assembly. These force fields have subsequently been used to study the self-assembly of SC lipids into multi-lamellar structures, enabling the examination of conditions more representative of experiment and probing the preferred conformation of CERs in the lipid lamella.^30,38^ For example, simulations performed using these MS-IBI forcefields have shown approximately 15% of the lipids were found to adopt extended conformations in the central layers (i.e., those not in contact with water) in pure CER NS systems, which increased to approximately 25% for systems with a 1:1 ratio of CER NS C_24_:FFA C_24_.^30,38^

We note that methods have been proposed to reverse-map or back-map CG systems to the atomistic level in order to generate atomistic structures not accessible from atomistic simulations alone and determine properties that require atomistic detail, such as hydrogen bonding.^22–24^ However, these methods have primarily been applied to fluid membranes, with only limited application to models of the SC,^28,41^ likely due to challenges associated with generating stable configurations for the dense, highly ordered CER-based membranes.

Here, we first present a MS-IBI based CG model for CHOL using the mapping scheme for CHOL proposed by Hadley and M^c^Cabe.^42^ The model is tested and validated through the study of a minimal model of the SPP phase of the SC comparing CG and atomistic simulations of bilayers. We then explore the effects of CHOL concentration on self-assembled multilayer structures containing mixtures of CER NS, CHOL and FFA. In particular, the tendency of CER NS to form hairpin rather than extended conformations in multi-layer lipid lamella is studied, along with analysis of the three-dimensional structural arrangements of the individual lipids within the multilayer structures. The CG simulation results are also used to construct atomistic multilayer systems that mimic the CG conformations were constructed using a simplified back-mapping approach, allowing for an accurate atomistically detailed examination of the structural ordering and hydrogen bonding of multilayer membranes.

## Computational Methods

In order to simulate a model SPP phase, we use a minimal 3-component model containing the ceramide N-(tetracosanoyl)-sphingosine (NS) with a saturated acyl chain of C_24_, designated CER NS C_24_ mixed with CHOL, and FFA C_24_. In experiments, this mixture formed lamellar phases resembling the SPP at molar ratios of 1:1:1 and 1:0.45:1 for CER NS C_24_:CHOL:FFA C_24_.^19^ CG models for CER NS, FFA, and water were developed in previous work.^30,38,39^ The schemes for mapping atoms in CER NS C_24_ and FFA C_24_ to CG beads are shown in Figure 1. CER NS C_16_ and FFA C_16_ were also simulated by eliminating the TER2 and two TAIL beads of the fatty acid chain.

### CHOL Model

The CG model for CHOL is shown in Figure 1a,b.^42^ In this mapping, proposed by Hadley and M^c^Cabe,^42^ the CHOL molecule is described as a 4-ring structure, denoted as A-D from left to right in Figure 1a, with 9 separate interaction sites, of which there are 5 unique sites or bead types for the nonbonded interactions. The CHOL head bead (CHEAD) represents the hydroxyl group and two of the carbon atoms in the A ring. The remaining carbon atoms in the 4-ring structure are divided into 4 RING beads. The CHOL tail attached to the D ring is divided into two distinct beads, CTAIL and CTERM. To capture the molecular roughness of one face of the CHOL molecule, the methyl groups attached to rings are each mapped to a CHME bead (i.e., chiral methyl).^42^ For the remainder of the discussion, the RING, CTAIL, CTERM, and CHME beads are collectively referred to as CBODY.

### MS-IBI Optimization

The MS-IBI method,^40^ which is an extension of the commonly used IBI methodology,^43^ was used to optimize the CG CHOL force field and cross interactions with other lipids.^40,44^ In its typical implementation, IBI performs self-consistent optimization of a CG interaction potential, whereby the CG interaction potential is modified at each step by comparing the radial distribution function (RDF) of the CG simulation to the RDF of the “target,” typically from an atomistic simulation of the system being reproduced. MS-IBI uses the same general procedure, but instead allows the CG interaction potential to be optimized based upon the agreement with multiple distinct targets; these targets may be for the same molecules or different molecules with a subset of shared chemical topologies, at distinct thermodynamic state points/structures, and/or at the same state point but simulated using different ensembles (e.g., constant volume compared to constant pressure simulations). This is accomplished by considering the correction to the CG interaction potential to be the sum of the deviations between RDFs for the multiple targets, as shown in Equation 1,

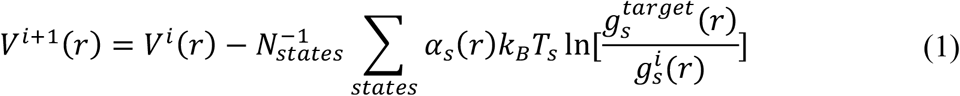

where *V*^*i*^*(r)* and *V*^*i*+1^*(r)* are the potential energy functions at the current and next iterations, respectively, for a given pair of atom types; 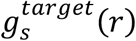 and 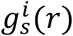 are the radial distribution functions between a given pair of atom types calculated respectively from the target atomistic system mapped to the CG level and the current iteration; α_*s*_*(r)* is a weighting factor for state *s, N*_*states*_ is the total number of states, *T*_*s*_ the temperature of the target state, and *K*_*B*_ the Boltzmann constant. For the case of a single state point, equation 1 reduces to the standard IBI procedure.

The relative weight (α_*s*_*(r)*) of each of the states allows greater emphasis to be placed on matches with targets of greater importance. In addition, α_*s*_ acts as a damping factor to stabilize the optimization procedure by preventing large update terms, which may slow down the rate of convergence. Additionally, for MS-IBI, the weighting functions are defined such that they linearly reduce from their maximum value from α_*s*_ *(0) =* α_,*max*_, to α *(r*_*max*_*) = 0* to suppress artifacts in the interaction potential that can arise due to intermediate and long-range correlations in the RDF following the work in reference.^45^ The values of α_*max*_ are chosen via guidance from prior optimization work and refined via guess and check iteration.

The nonbonded interactions for CG CHOL were parametrized using the MS-IBI approach to match several target states from atomistic simulations in order to improve state transferability of the model, as compared to the original parameters determined by Hadley and McCabe using single-state IBI.^46,47^ To capture a range of behaviors, the target states for optimizing the CHOL force field included bulk fluid phases, monolayers, bilayers, and dehydrated multilayers, as detailed in the supplemental information. Surface wetting simulations were also performed for select interactions to measure contact angles and aid in gauging the accuracy of the strength of the hydrophobic interactions in the CG model.^38^ Complete details of the optimization process are provided in the supplemental information.

### Simulation Methodology

All atomistic simulations employed the CHARMM36 force field^46^ with additional parameters from Guo et al.^47^ for CER NS and from Cournia et al.^48^ for CHOL taken Atomistic simulations are performed using the GROMACS simulation engine version 2018^49^ with a 1 fs timestep. Van der Waals interactions were truncated at 12 Å with a smooth switching function applied between 10 and 12 Å. Short-range electrostatics were truncated at 12 Å and long-range electrostatics calculated using the particle-mesh Ewald (PME) method. NVT and NPT simulations employed the Nose-Hoover thermostat with a coupling constant of 1 ps.^50^ For hydrated lipid bilayers, separate but identical thermostats were applied to the water and the lipids. NPT simulations used a Parrinello-Rahman barostat^51^ with a coupling constant of 10 ps. Semi-isotropic coupling (in the *xy*-dimensions, along the bilayer plane) were used for bilayer systems, whereas isotropic coupling was used for the bulk fluid systems. For ordered systems, such as bilayers, the random walk MD (RWMD) procedure^22^ was used to allow lipids to decorrelate from their initial orientation and in-plane morphology. The exact simulation protocol is detailed in the supplemental information.

The CG simulations were performed using the HOOMD-Blue simulation engine, version 2.9.3.^52^ A timestep of 10 fs was used in all of the CG simulations. Full details of the simulation parameters used and the accuracy of the individual CG RDFs as compared to their atomistic targets mapped to the CG level are in the supplemental information. Simulations of the self-assembly of CG model membranes were initialized with randomized configurations using the mBuild software.^53^ Bilayer systems contained 1,000 lipid molecules and 4- and 6-leaflet systems contained 2,000 lipid molecules. All systems contained 10 water beads per lipid (10,000 and 20,000 water beads respectively in total for the bilayer and multilayer systems), which translates to 40 water molecules per lipid. Four independent replicates each starting from randomized initial configurations were simulated for each self-assembled CG system studied. To accelerate self-assembly the box annealing protocol described in Moore et al.^30^ was employed. Finally, production runs for the self-assembled systems were carried out in the NPT ensemble for 200 ns at 1 bar and 305 K using the Martyna-Tobias-Klein thermostat-barostat with the same coupling constants as the atomistic simulations.^54^ Additional details of the system set up are provided in the supplemental information. Input files for simulations were prepared using the mBuild software as part of the Molecular Simulation and Design Framework (MoSDeF).^53^

### Analysis

Analysis of the lipid systems was performed utilizing the MDTraj^55^ and Freud^56^ software packages. Bilayers and multilayer systems were characterized using the area per lipid (APL), bilayer thickness (d), interdigitation, tilt angle, and nematic order parameter (S_2_). The fraction of ceramides in the hairpin and extended conformations and the number and location of water beads were measured for the inner leaflets of multilayer systems. To further describe in-plane morphology, the coordination number between specific lipid types was determined. For atomistic systems, neutron scattering length density (NSLD) profiles and hydrogen bonds were also calculated. Details of how these quantities were determined are in the supplemental information. Results are reported as mean ± standard deviation for four replicate simulations initialized with different randomized configurations

## Results and Discussion

### CHOL force field validation

The CG CHOL force field obtained from the MS-IBI optimization is able to accurately describe pure CHOL. The atomistic and CG simulations are in good agreement for bulk fluid properties; for example, the density of pure CHOL at 550K and 1 atm is reproduced with high accuracy (0.73 ± 0.01 g/mL (CG) versus 0.73 ± 0.01 g/mL (atomistic)), as is that for an equimolar mixture of CHOL and FFA C_24_ at 1 atm and 500K (0.81 ± 0.03 g/mL (CG) versus 0.79 ± 0.01 g/mL atomistic). High temperatures were chosen for these systems to ensure that the lipids remain in a fluid state. At 298K, a density of 0.87 ± 0.01 is observed for CG simulation of pure CHOL, which is lower than the experimental density of 1.02 g/mL measured for pure CHOL in the crystalline state at room temperature.^57,58^ Good agreement is also obtained between the atomistic target and CG RDFs (Fig. 2). In terms of cross-interactions with CER NS and FFA, the CG force field is in close agreement for ordered states, when considering the RDFs between the CHOL headgroup (CHEAD) and the headgroups in CER and FFA, but deviates more for bulk states, where the CG model predicts stronger association between the head groups than is seen in the atomistic simulations. Because the focus of this work is on the formation of ordered phases, which are given a higher weighting factor in the MS-IBI derivation, these bulk phase deviations are considered acceptable. All of the potentials and RDFs from the MS-IBI optimization of the CHOL CG forcefield and its cross interactions with other SC lipids are provided in the supplemental information.

**Figure 2:**
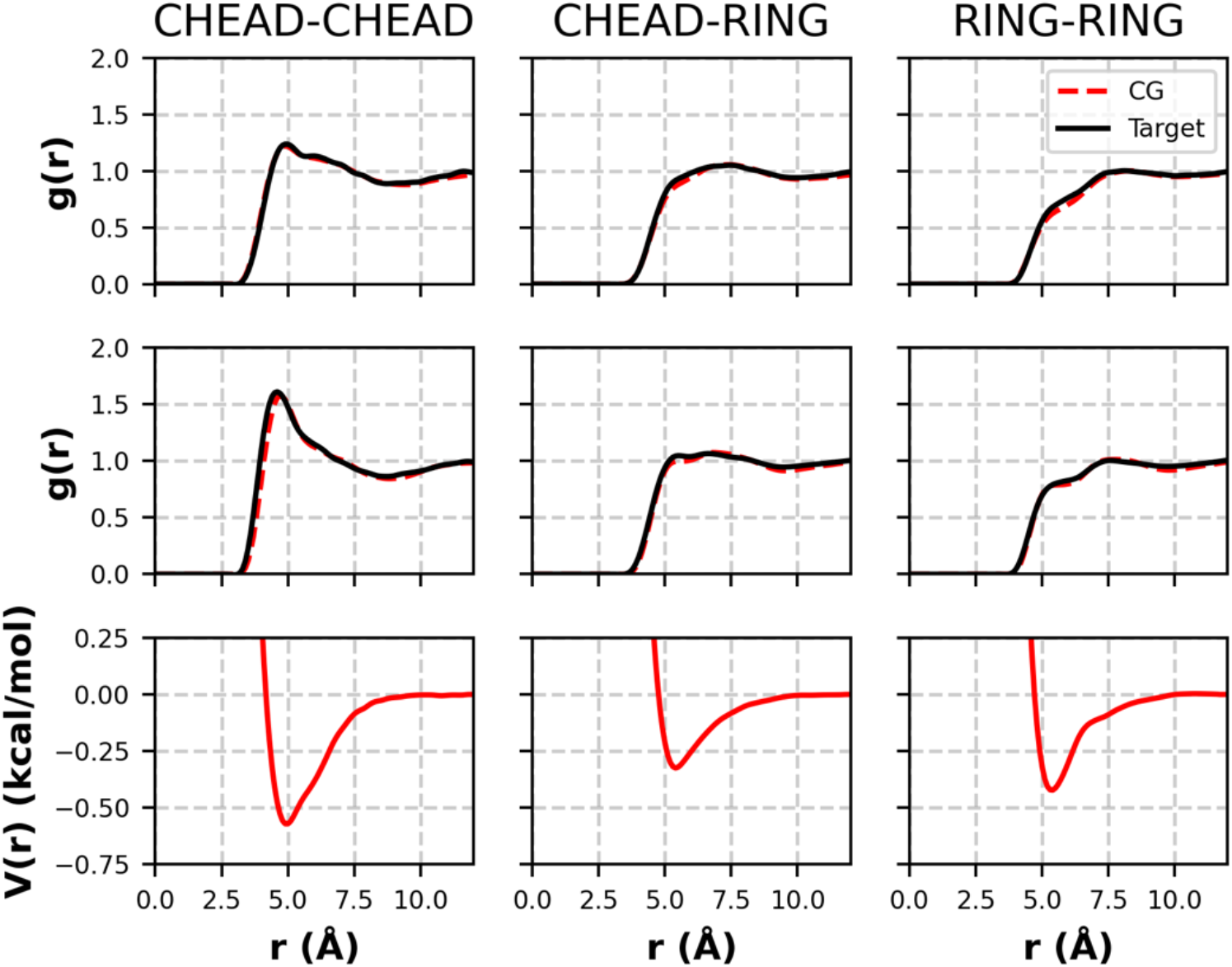
RDFs and pair potentials from the pure CHOL force field optimization. Top: target (black) and CG (red) RDFs at the 550 K NPT state; middle: target (black) and CG (red) RDFs from the 400K NVT state; bottom: pair potential that yields the CG RDFs above.

In bilayer simulations of two or three lipids, the CG force field reproduces with good accuracy the structural properties of the pre-assembled atomistic bilayers in both pre-assembled and self-assembled CG simulations at 305K (Table 1). Three system were studied: CER NS C_24_/CHOL, allowing the CHOL/CER NS cross interactions to be tested, CHOL-FFA C_16_, allowing the CHOL/FFA interactions to be tested, and CER NS C_24_/CHOL/FFA C_24_, the system of particular interest in this study. Because the pre-assembled CG simulations were initialized using the final configuration of the atomistic simulation mapped to the CG level, the differences between the CG and atomistic structure for pre-assembled bilayers is primarily driven by the CG force field rather than the equilibration procedure involved in self-assembled systems.

**Table 1:**
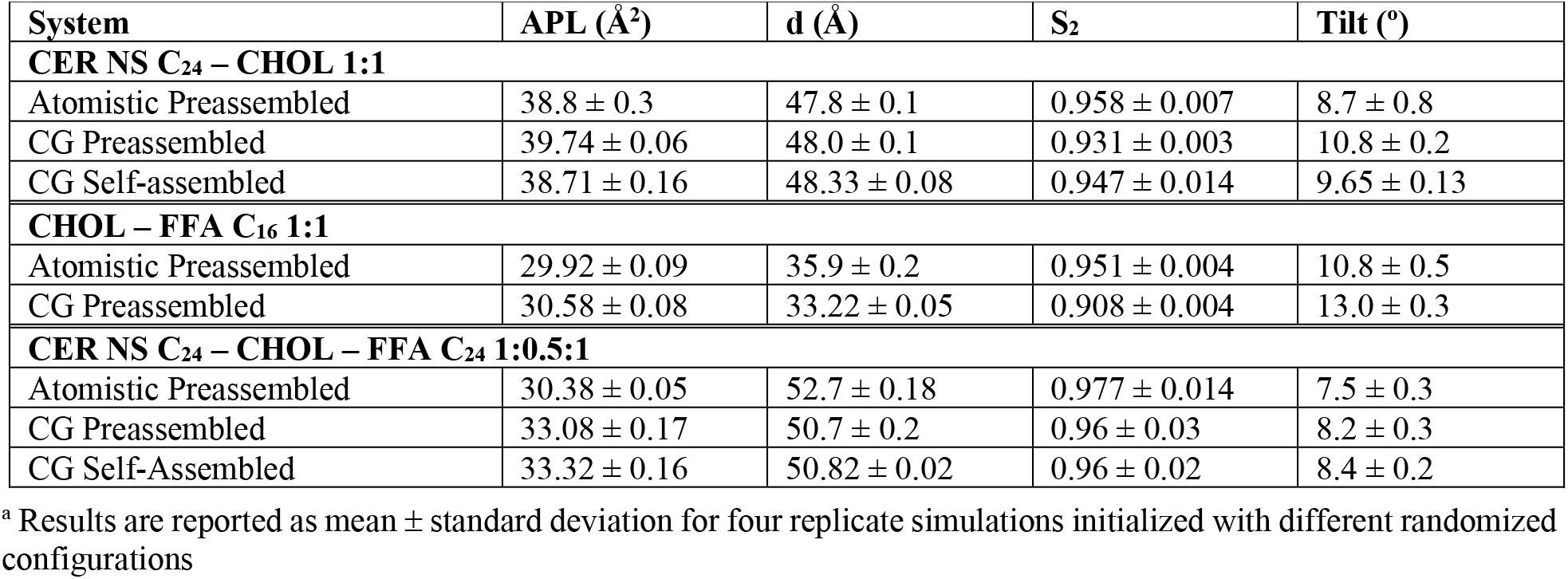
Structural properties of the atomistic and CG systems studied at 305K^*a*^.

For pre-assembled bilayers composed of CER NS C_24_/CHOL and CHOL/FFA C_16_, both in 1:1 ratios, the CG area per lipid (APL) is within 2.5% of the atomistic APL, showing that the CG model captures the in-plane packing of the lipids, similar to prior results for pure CER NS and CER NS/FFA mixtures. The thickness of the CG CER NS C_24_/CHOL bilayer shows almost perfect agreement with the atomistic bilayer, while the CG model of CHOL/FFA C_16_ underpredicts the atomistic bilayer thickness by *∼*7.5%. Lafleur et al.^59,60^ studied equimolar CHOL/FFA C_16_ systems experimentally via NMR and observed a bilayer phase between 330 K and 340 K with a height of 32 Å, which is close to the 35.7 Å height observed for the CG bilayers at 330 K, which is ∼8.5% higher than the experimentally measured height. An APL of 30.9 Å2 was observed for our CG model, which is ∼8% lower than the APL obtained for the same system via atomistic simulation.^61^ The change in APL for the equimolar CHOL-FFA C_16_ system from 305 to 330 K was much smaller for the CG model than the atomistic model (30.58 to 30.9 and 29.92 to 33.3 Å respectively). This suggests that the temperature dependence of the CG model is much weaker than the atomistic. Hadley and M^c^Cabe had also used their IBI-derived CG force field to self-assemble equimolar mixtures of CHOL and FFA and obtained a bilayer height of 30 Å which is slightly lower than the experimentally measured height. They also measured an APL of 33.3 Å at 330 K, which is in near perfect agreement with atomistic simulations. However, it is important to note that the IBI model developed by Hadley and McCabe used the equimolar CHOL and FFA bilayer at 330 K as the only target state for CHOL/FFA cross-interactions, whereas our model did not use this state.

In both the CER NS C_24_-CHOL and CHOL-FFA C_16_ systems, the nematic order of the lipid tails indicates they are highly ordered, although the CG model underpredicts the atomistic results. The CG model overpredicts the tilt angle for both systems, although only slightly. Comparison to a pre-assembled bilayer of the 3 components, CER NS C_24_, CHOL, FFA C_24_ in a 1:0.5:1 mixture also shows close agreement in terms of structural properties, as reported in Table 1.

Turning to the self-assembled systems, again, close agreement is observed with the atomistic results in terms of structural properties. Comparison of the in-plane structure, as quantified via CHOL-CHOL, CHOL-CER NS, and CHOL-FFA coordination numbers, also shows good agreement with the atomistic simulations. Note, these atomistic simulations use the RWMD procedure in order to relax and decorrelate lipids from their initially, assumed configurations and morphologies. Additionally, these 3 component lipid simulations provide an unbiased validation of the CG force field because they were not included in the target optimization.

### Self-Assembly of lamellar structures

Self-assembly simulations have been performed with varying CHOL concentration using the validated MS-IBI CG force fields discussed above for 3 component mixture of CER NS, CHOL and FFA to explore the effect of lipid composition on structural properties. Differences in the behavior of a bilayer sandwiched between water layers compared with a bilayer between other bilayers were explored in self-assembly simulations of 2-, 4- and 6-leaflet systems.

#### Comparison of bilayer and multilayer structures

As noted earlier, experimental SC lipid systems exhibit multilamellar structures, typically containing more than 20 layers.^61^ Thus, almost all lipids in these systems reside in interior bilayers without direct contact to bulk water. Experimental results, therefore, such as X-ray scattering profiles and FTIR and NMR spectra, represent the behavior of these interior lipids, which are fundamentally different from the single hydrated bilayer studied in most simulations. First, interior lipids in experimental lipid systems have very low hydration, only 1-2 water molecules per lipid in the head group region, leading to more interactions between head groups.^15^ In contrast, simulated hydrated lipid bilayers contain large amounts of water, which produces fully hydrated headgroups and reduces head group interactions. Second, because there is a large difference in the amount of water, CERs located in interior bilayers can adopt an extended conformation, which is impossible for a single hydrated bilayer because contact between water and CER tails is highly unfavorable.

To investigate the differences between bilayer and multilayer systems, mixtures of CER NS C_24_/CHOL/FFA C_24_ in a 1:0.5:1 molar ratio were self-assembled in to 2-, 4-, and 6-leaflet structures (Fig. 3). In the following analysis, the head groups of the exterior leaflets are in contact with the bulk water interface whereas the head groups of the interior lipid leaflets are in contact head groups of another leaflet. The self-assembled multilayer systems naturally contain low lipid hydration for the interior lipids: 1.5 water molecules per lipid for the 4-leaflet system and 2.6 for the 6-leaflet system. This is in good agreement with ∼2 waters per lipid based on neutron scattering measurements^15^ observed in the aforementioned experimental model SPP systems containing a mixture of CERs, CHOL, and FFAs of varying tail lengths. In addition, a significant fraction of interior CERs adopt an extended conformation (∼35-36%) in the simulated structures.

**Figure 3:**
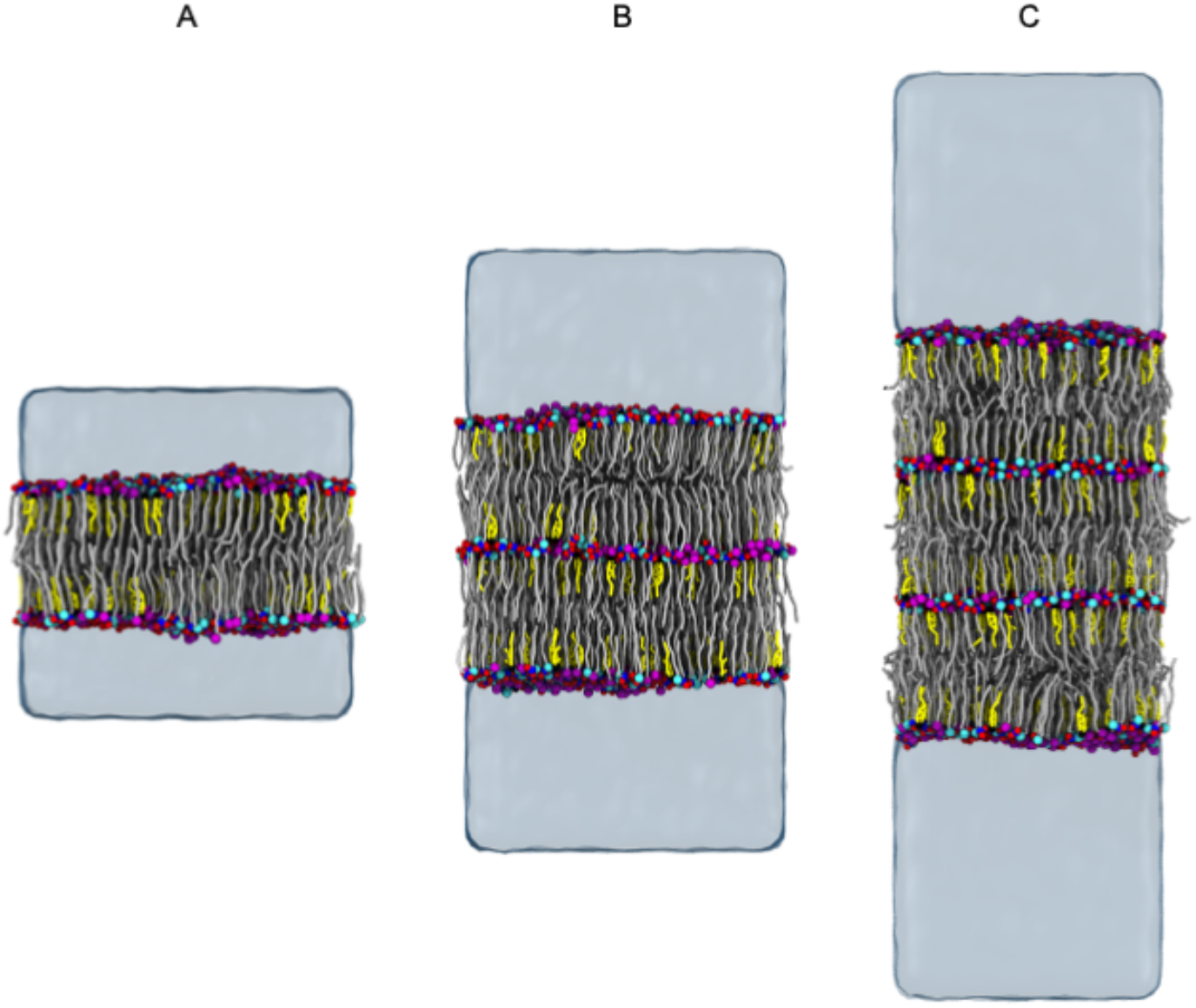
Simulation snapshots of self-assembled mixtures of CER NS C_24_/CHOL/FFA C_24_ with a 1:0.5:1 molar ratio in (A) bilayer, (B) 4-leaflet, and (C) 6-leaflet configurations. The lipid molecules are colored as defined in Figure 1. Water is represented as a transparent surface.

The combination of low hydration and the presence of extended CERs in the interior layers of the multilayer systems affects structural properties as shown in Table 2, which reports the average across all leaflets, except for the fraction of extended CERs and the water molecules per lipid, which are calculated from the leaflets without bulk water contact. The decrease in APL for the 4- and 6-leaflet systems compared to the 2-leaflet systems, can be attributed to the increase in extended CERs, which pack more efficiently. At the same time, the tilt angle for the 6-leaflet system is larger than for the 2- or 4-leaflet systems. Figure S6, which shows the distribution of angles between the acyl and sphingosine chains, indicates that the angle between the tails of extended CERs is ∼170º rather than the fully linear 180º. This slightly “V”-shaped conformation may arise to optimize chain packing similar to the V-shaped arrangements proposed for CER NP C_24_ mixed with CHOL and FFA C_24_,^62^ as well as those observed for pure CER NP crystals. Furthermore, the increased fraction of extended CERs in this slight V-shaped conformation may impart the higher tilt angle observed in the 6-leaflet system. The average bilayer height increased by ∼2-3 Å as the number of layers increases, which is consistent with the reduced lipid interdigitation. The bilayer height of 53.81 Å for 6 layers closely agrees with the experimentally measured 54 Å height of the SPP phase, likely due to the similarities in the lipid hydration and the presence of extended CERs.^15,20^

**Table 2:**
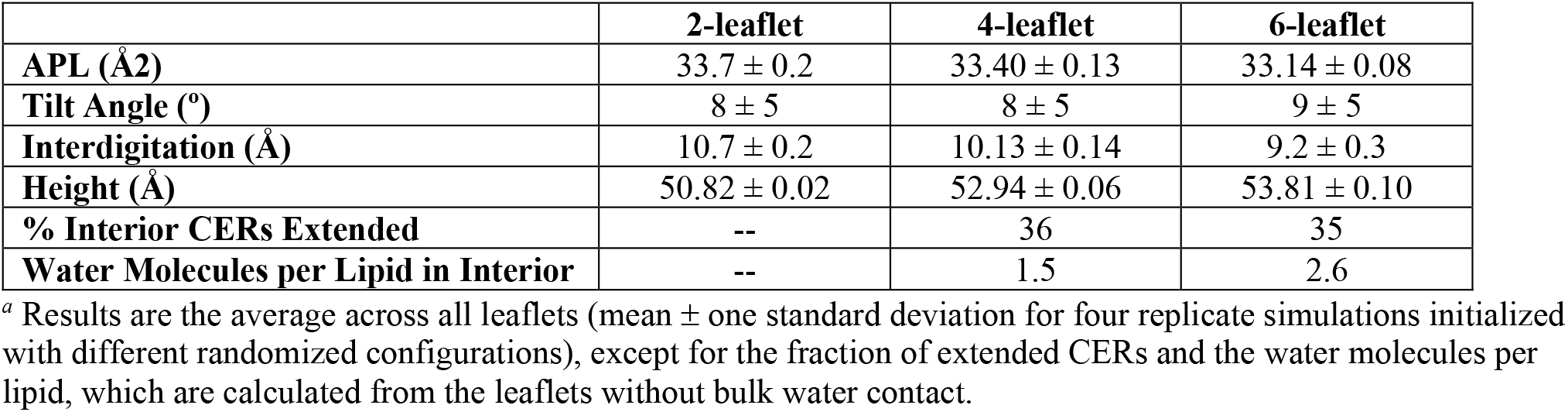
Average structural properties of 2-, 4-, and 6-leaflet systems of CER NS C_24_/CHOL/FFA C_24_ with a molar ratio 1:0.5:1 ^*a*^

#### Influence of CHOL content on self-assembled CG systems

To examine the role of CHOL in the SC, the mole ratio of CHOL in the minimal SPP systems containing equimolar CER NS C_24_ and FFA C_24_ was varied from 0 to 1 relative to CER NS in self-assembly simulations (Fig. 4). As expected, extended CER NS molecules are abundant in the inner bilayers of the 6-leaflet systems and absent in the outer leaflets (Figure 5a). However, the fraction of extended CER NS molecules varies as a function of the CHOL content, with equimolar CHOL:CER NS demonstrating the highest fraction and also the highest area per lipid tail (APT). Perhaps the increased concentration of bulky CHOL tails and consequent increase in APT is countered by an increase in extended CERs, which pack more efficiently.

**Figure 4:**
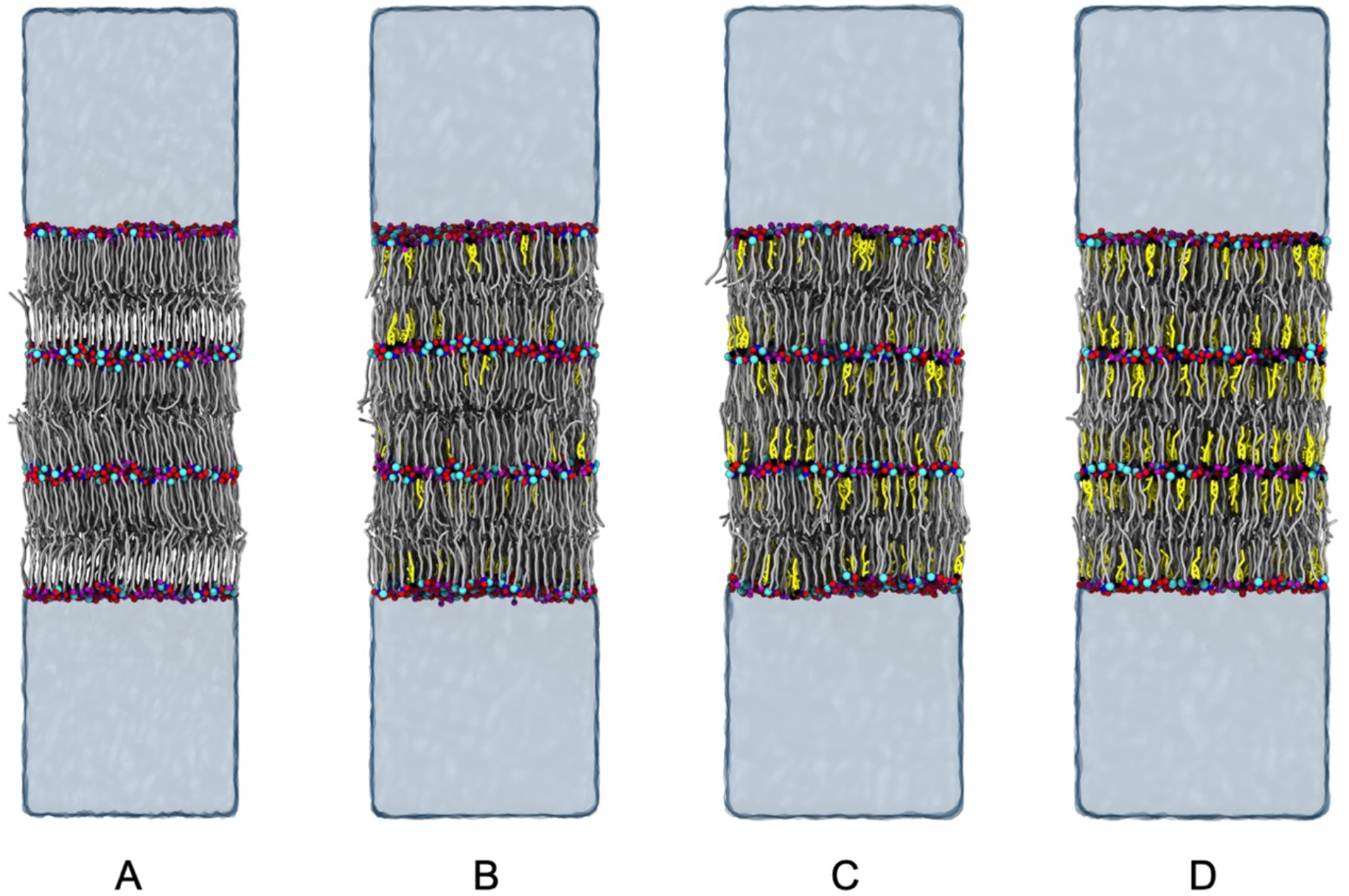
Snapshots from simulations of the 6-leaflet system of CER NS C_24_/CHOL/FFA C_24_ with mole ratios of a) 1:0:1, b) 1:0.2:1, c) 1:0.5:1, and d) 1:1:1. The lipid molecules are colored as defined in Figure 1. Water is represented as a transparent surface.

**Figure 5:**
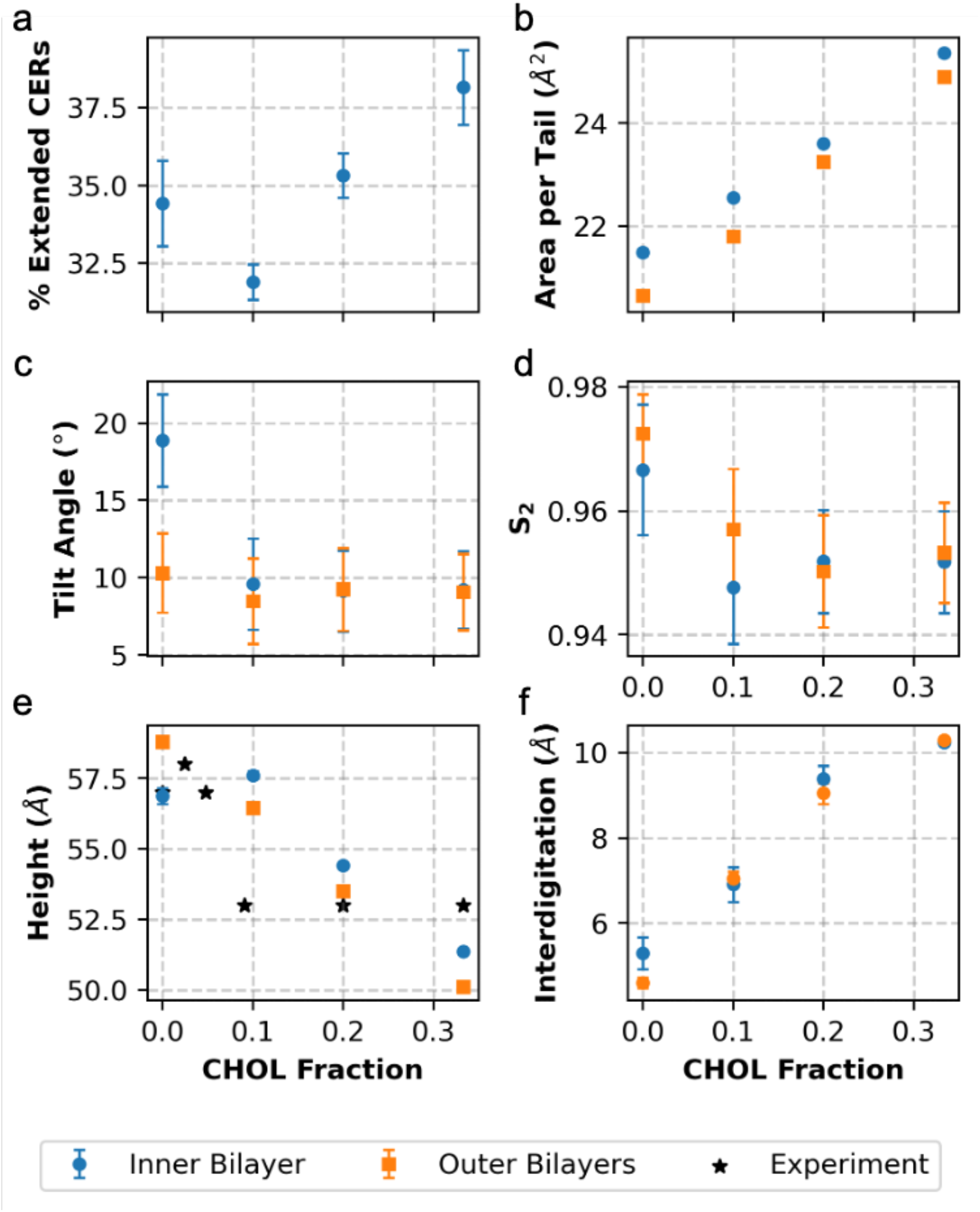
Average structural properties for the inner (blue circles) and outer (orange squares) bilayers of the 6-leaflet system with varying amounts of CHOL in an equimolar mixture of CER NS C_24_:FFA C_24_: a) t CERs extended, b) tilt angle, c) APT, d) S_2_, e) bilayer height, and f) interdigitation). Results are reported as mean ± one standard deviation from four replicate simulations initialized with different randomized configurations. Experimental height measurements (stars in e) from Mojumdar et al.^20^ are comprised of a mixture of CER types, although predominantly CER NS C_24_ (60 molt) CHOL, and FFAs of varying tail length from C_16_ to C_26_.

Figure 5 shows the trends in the structural properties of the central bilayer in the CER NS C_24_/CHOL/FFA C_24_ systems as a function of CHOL concentration. From the figure, we can see that the area per tail increases as the fraction of CHOL increases. This is consistent with experimental measurements in which hydrocarbon lipid tails typically have a cross-sectional area of 20 Å^2^ while cholesterol molecules have a cross-sectional area of roughly 38 Å^2^.^63^ There is a significant decrease in the S_2_ and tilt angle upon addition of CHOL into the bilayer, indicating that the increased CHOL content disrupts the hydrocarbon tail packing. However, further increases in CHOL content did not have a significant effect on the S_2_ and tilt angle. Also plotted in Figure 5 is a comparison of the same properties for the outer bilayers of each system. The structure of these interfacial leaflets, which would be characteristic of single bilayer systems, have several distinct differences as compared the dehydrated inner region. In particular, the outer bilayers have a smaller tilt angle than the outer bilayers for the 1:0:1 system, which may be attributed to fewer ceramides in the extended conformation, as seen when compared to bilayers in Table 2. A layer-by-layer analysis demonstrates differences in the number of lipids at the water-lipid interface, as compared to the inner leaflets with a lipid-lipid interface. Similarly, the APL and tilt angle differ within these inner leaflets as compared to those in contact with water. As explained earlier, the presence of extended CERs increases the APL and decreases the tilt as observed when comparing multilayer and bilayer systems of CER NS C_24_/CHOL/FFA C_24_ in a 1:0.5:1 ratio. The most significant difference is the ability of the multilayer structures to form both hairpin and extended conformations. As can be seen from the figure, the number of extended conformations within these central layers increases from ∼34 to 38% as the cholesterol content is increased. Finally, we note that comparison to experimental repeat distances when available shows close agreement (see Figure 5d) in both trends and quantitative values, though we note that the experimental systems contain a mixture of CERs NS, NP, AP, AS, and 7 FFAs tail lengths ranging from C16-C26.

In order to examine the preferential arrangement and aggregation of lipid tails in the central bilayer, a cluster analysis was performed to identify the number and size of lipid tail clusters. In Figure 6a we note that the normalized number of CHOL clusters decreases as the fraction of CHOL increases. This indicates that there is a higher fraction of large clusters formed. This is also supported by Figure 6b, which shows that there is an increase in the cluster size as the CHOL content increases. One should also note that there is a sharp increase in the cluster size between the 1:0.5:1 and 1:1:1 systems. This supports the experimental observation that found phase separation of CHOL occurs at CHOL fractions above 0.2 for SPP systems;^20^ in our system the CHOL stays within the membrane, rather than forming a separate phase, but clearly phase separates within the layer. One can also see that there is an increase in the normalized number of clusters and a decrease in the mean cluster size for FFA and CER tails as the fraction of CHOL increases, which suggests that CHOL may improve the miscibility of FFA and CER tails in the membrane.

**Figure 6:**
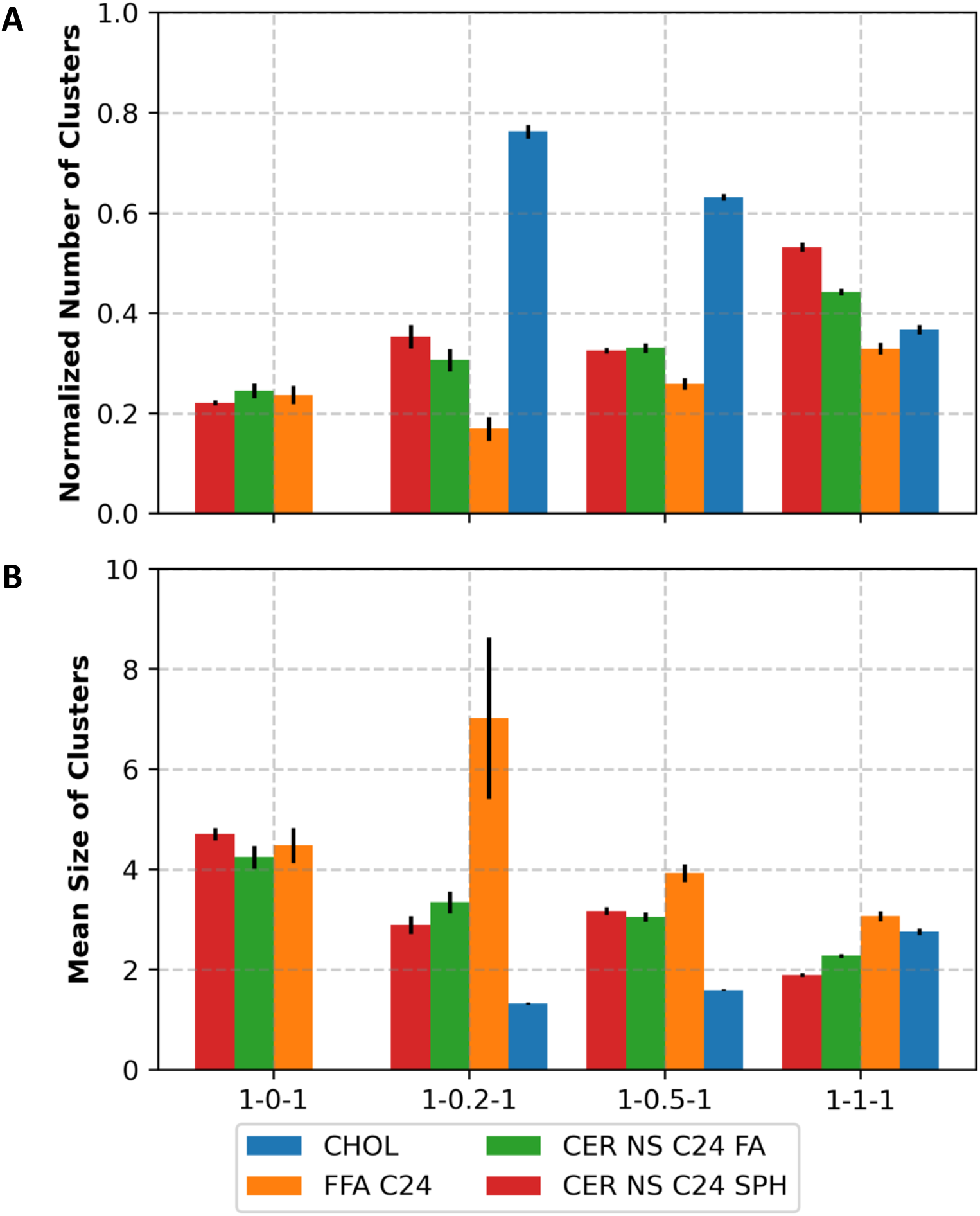
Cluster analysis of lipid tails in the central leaflet. A) the number of clusters normalized by the number of lipids for each lipid type. A value of 1.0 indicates that all clusters contain only a single molecule, meaning that the lipids of that type are fully dispersed. Lower values indicate that more multi-lipid clusters have formed. B) the mean cluster size is shown. All values are averaged over a 2000 frames trajectory spanning 20 ns of equilibrated simulation.

To further characterize the preferential arrangement of CHOL molecules in these systems, the coordination number of the CHOL tails with each of the lipid tail types was calculated. One should note due to the sparsity of CHOL-CHOL pairs, the coordination number cannot be precisely calculated as the first peak of the CHOL-CHOL RDF is very short. From the results shown in Figure 7, we can see that between the sphingosine and acyl CER tails, CHOL prefers to neighbor the similar-length sphingosine tails. This is corroborated by experimental NMR spectra, which suggests that the sphingosine chain is more likely to neighbor CHOL molecules in a mixture of CER NS C24, CHOL, and FFA C24 in a 1:0.45:1 ratio.^19^ However, these experiments were not able to quantify the extent of this preference and assumed in their proposed model that all of the sphingoid chains of CER neighbor CHOL molecules.

**Figure 7:**
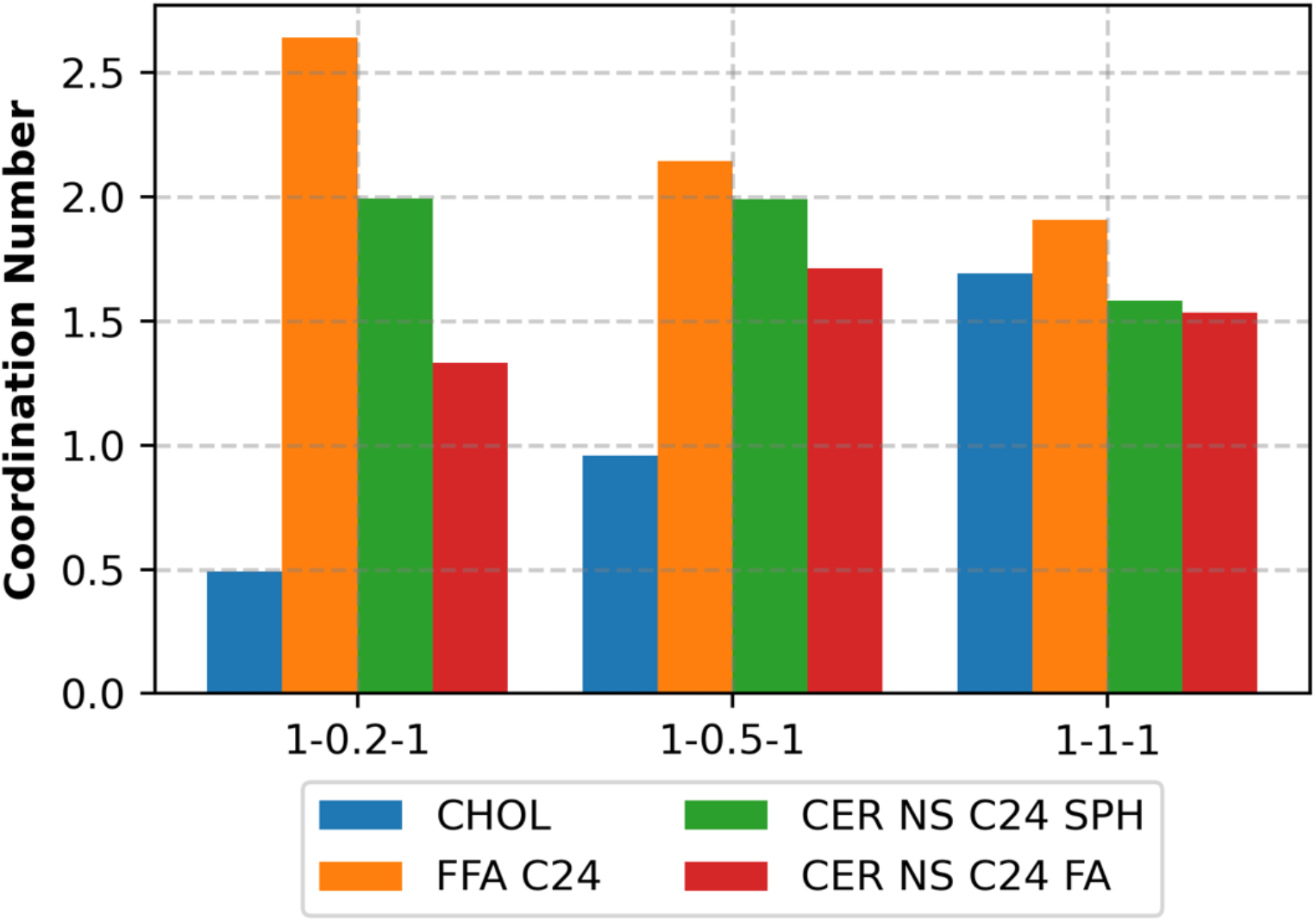
CHOL-X coordination numbers, where X is either CHOL, FFA C_24_, the sphingosine chain of CER NS C_24_ (SPH), or the fatty acid chain of CER NS C_24_ FA.

From the data shown in Figure 7, we can see that although there is a slight preference for CHOL to neighbor sphingoid chains, CHOL still has a significant number of CER acyl chain neighbors. It is also interesting to note that, contrary to the model hypothesized by Engberg et al., CHOL prefers to neighbor FFA molecules.^19^ This may be because FFA molecules have higher mobility compared to CERs due to their single-tailed nature, which may more effectively accommodate packing defects induced by the bulky CHOL tails. This is further supported by the Voronoi analysis provided in the SI, which indicate that 5-neighbored FFAs mitigate hexagonal packing defects introduced by 7-neighbor CHOL molecules.

### Recovering Atomistic Details

As previously discussed, the CG model and force field used in this work is capable of reproducing, with good accuracy, many of the key structural metrics of the corresponding atomistic bilayers. However, the loss of detail inherent to the CG model necessitates the use of atomistic-level configurations to be able to compare against experimental neutron scattering length density (NSLD) measurements as well as to accurately calculate quantities such as hydrogen bonding and the localization of atomic species. Here, a simple procedure is used to reintroduce atomistic level details into the CG multilayer configurations generated via self-assembly. Rather than utilizing a reverse-mapping approach such as in Wassenaar et al.,^64^ which requires constraints and careful equilibration due to high energy states and overlaps associated with inserting atoms into the CG configuration, we instead employ a simpler approach that utilizes information from the self-assembled CG simulation to initialize a pre-assembled atomistic configuration. In this approach, the morphology (i.e., in-plane arrangement of lipids) and CER conformations (i.e., hairpin or extended) are determined from the self-assembled CG structure and the atomistic lipids are arranged accordingly. With this configuration generated, the same general procedures used for the pre-assembled atomistic bilayers are employed to relax the system. Since morphology is derived from the unbiased self-assembled simulation, RWMD is not required since the main goal is to allow lipids to perform a local energy minimization while retaining the morphology dictated by the self-assembled configuration, akin to reverse-mapping.

Here we focus on generating an atomistic configuration for the system shown in Fig. 4c, namely a 6 layer 1:0.5:1 (CER NS: CHOL:FFA) mixture containing 2000 total lipids, hydrated with 26,666 water molecules. As is typical for initializing atomistic bilayers, the in-plane area of the initial configuration is set larger than the anticipated equilibrium area to avoid overlaps between the lipids; in this case, the initial in-plane box dimensions are larger than the final dimensions by a factor of ∼1.2. All lipids in the atomistic multilayer have their in-plane rotational orientation randomized, similar to the bilayer simulations above and in prior work,^22^ with the individual lipid tilt angle set to match the average value from the CG simulation. To relax this configuration, first, a steepest-descent energy minimization is performed to remove high energy states; this is followed by a 10 ns NVT step at 305K to relax molecular structure; next an NPT step is performed for 100 ns at 305K and 1 atm to allow the box shape (i.e., in-plane area) to relax and for lipids to further minimize their energy; a final 20 ns NPT production step at 305K and 1 atm is performed in order to gather simulation data for analysis. We note that, while the self-assembled CG configuration had a small number of water molecules associated with the headgroups within the inner leaflets (see Table 2), these water molecules were not considered when constructing the atomistic system.

A simulation snapshot of the relaxed atomistic multilayer configuration for a 1:0.5:1 (CER NS: CHOL:FFA) mixture at 305K and 1 atm is shown in Figure 8a, with lipids colored according to the key in Figure 8b; we note that extended CERs are colored differently than hairpin CERs to highlight their presence. Considering the regions in contact with water separately from those not in contact, there is very small variation from the nominal 1:0.5:1 ratio of CER NS:CHOL:FFA, where the layers in contact with water have ratio 1:0.52:1.09 (raw totals: 262:136:288) and those not in contact with water 1:0.49:0.95 (raw totals: 538:264:512), with 35% of the CERs (187) in an extended conformations in the central region. The tilt of the atomistic system is on average 9.9 ±8.2° compared to 9.2±5.3° for the CG configuration. We note that average tilt angle from the CG system was used to set the initial tilt angle of the lipids in atomistic system, and thus we might expect such agreement; however, due to starting with a configuration that was expanded by a factor of ∼1.2 in the in-plane dimensions, lipids were provided with sufficient freedom to adjust this angle if the initial was unfavorable. The average APT for the atomistic system is calculated to be 22.63 ± 0.03 Å^2^ which is slightly smaller than 23.36 ± 0.05 Å^2^ for the CG system.

**Figure 8.**
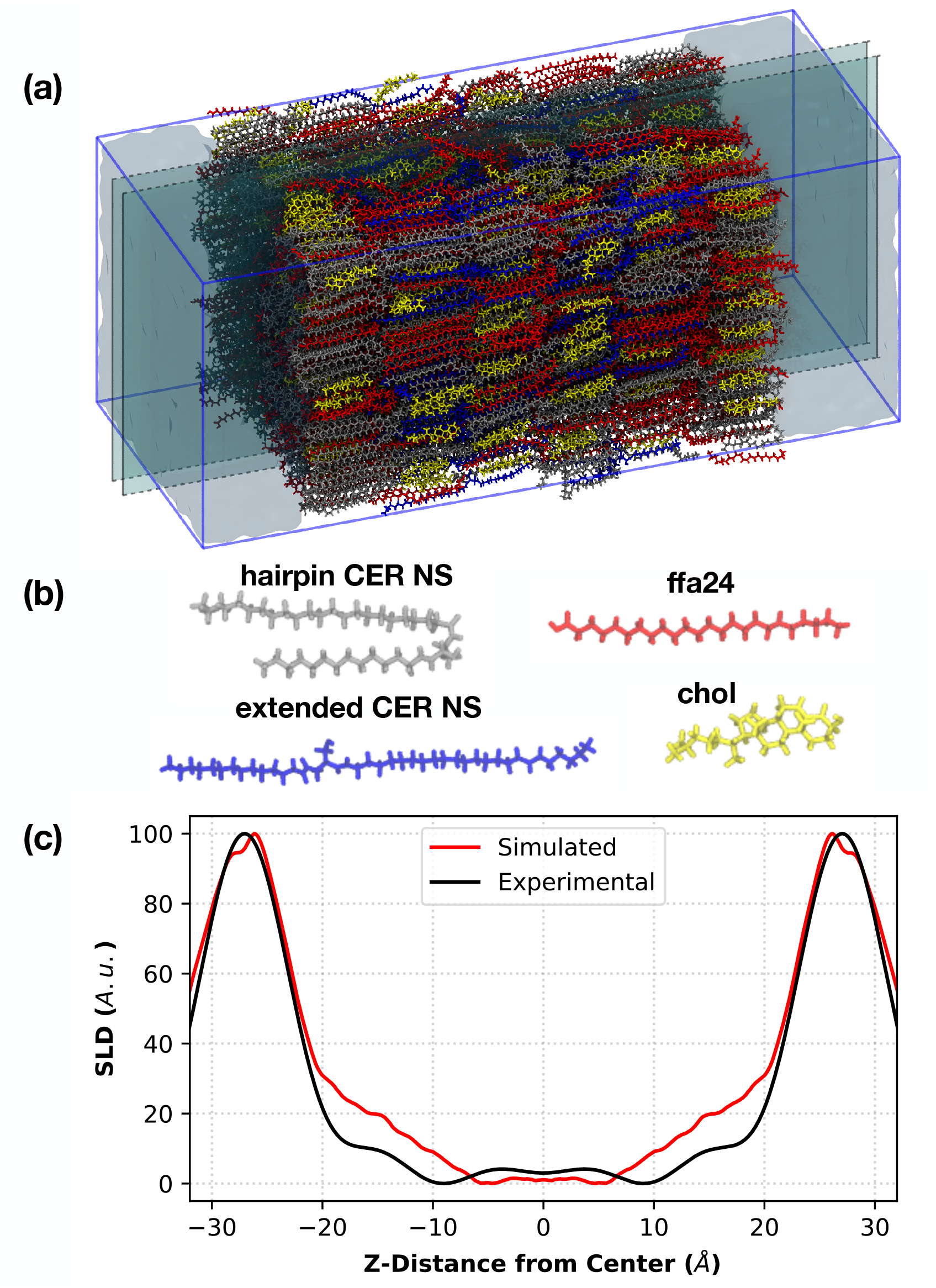
(a) Simulation snapshot of a reverse-mapped atomistic 1:0.5:1 CER NS C_24_/CHOL/FFA C_24_ multilayer, with lipids colored according to (b). A 1 nm thick slice is highlighted in (a) and shown in detail in Figure 9a. (c) Protiated NSLD profiles for the same reverse-mapped 6-leaflet system from the atomistic simulation and experiment reproduced from Groen et al.^15^

As a means of validating the configuration, the NSLD profile is calculated and shown in Figure 9, focusing on the two central most layers, compared to an experimental NSLD for a system containing a mixture of CERs NS, NP, AP, and AS, CHOL, and FFAs ranging from C_16_ to C_26_; while the experimental composition is more complex, we note that CER NS C24 is the dominant CER species at 60 mol%, making it a reasonable point of comparison. Good agreement between the simulated system and experiment is observed, where the distance between the main peaks are ∼52 Å and 53 Å for the simulated and experimental profiles, respectively. Moore et al.^22^ and Wang and Klauda^23^ both made comparisons to the same NSLD experimental data, but considering a 1:1:1 (CER NS:CHOL:FFA) mixture in a bilayer configuration. The bilayer configuration demonstrated a slightly reduced profile width (Moore, et al. report 50 Å, Wang and Klauda 49 Å), which may be expected given the increased concentration of shorter length CHOL molecules, enabling increased interdigitation; most notably though, the peaks in the NSLD for the bilayer are significantly narrower than those seen in either the experiment or the multilayer simulation, owing to the fact these regions in bilayer simulations are at a water-lipid, rather than a lipid-lipid interface.

**Figure 9:**
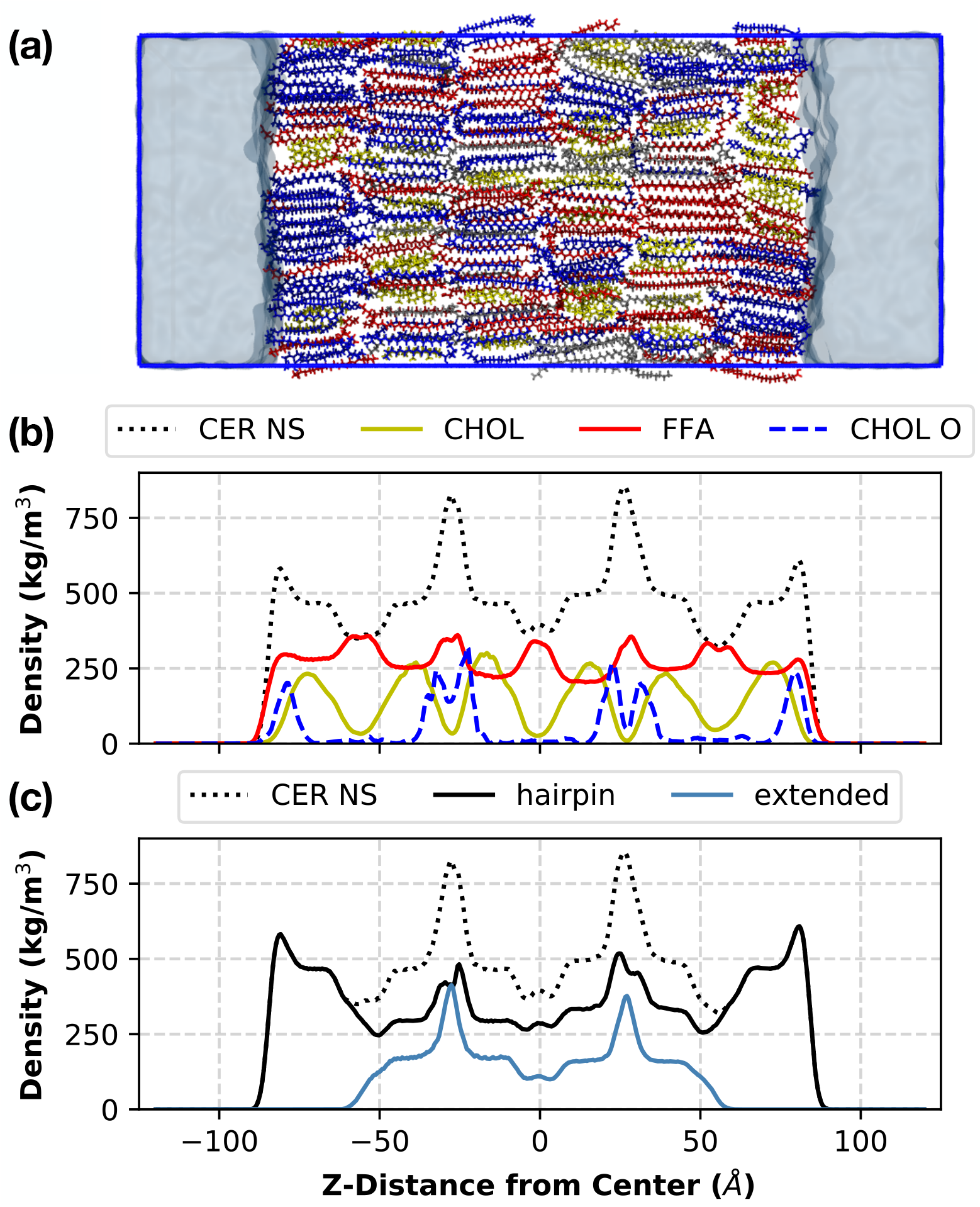
(a) Slice from the center of the simulation cell for the 1:0.5:1 CER NS C_24_/CHOL/FFA C_24_ multilayer shown in Fig.8a, again colored following Figure 8b. (b) density profile of CER NS, CHOL, and FFA species. The density profile of the oxygen in the CHOL headgroup is also plotted, with the numerical values scaled by a factor of 10 in order to make them visible on the plot. (c) Density histograms for CER NS, plotting hairpin and extended CERs separately.

To demonstrate the ordering of the lipids, Figure 9a renders a slice of the system (as highlighted by the two planes in Figure 8a) using the color scheme shown in Fig 8b. To quantify the distribution of the species in Figure 9a, Figures 9b and c plot the density histogram of the lipid components of this system. The z-distance from the center (i.e., x-axis) in Figures 9b and 9c correspond to the spatial dimensions shown in 9a. Consistent with experimental observations, FFA chains show increased density within the central region of a pair of leaflets (as seen at z-positions at ∼ −60, 0, and 60Å in Figure 9b), due to interdigitation of the chains. Also consistent with experiment, CHOL appears to sit deeper in the leaflet, away from the headgroups of the CERs and FFA. Experimentally this distance has been measured to be ∼4Å from the edge of the unit cell, at position ±22.9Å; examination of the density profile of the headgroup oxygen in CHOL, labeled as CHOL O in Figure 9b, we identify a peak location at ±22.8Å in very close agreement with experiment (as noted in the caption to Figure 9b, this histogram of CHOL O is scaled by a factor of 10 such that it can be seen on the same scale). Figure 9c, plots the overall density histogram of CER NS, along with histograms corresponding to those molecules in hairpin and extended conformations separately. At ∼ ±25 angstroms the hairpin CERs shows two small peaks, corresponding to two layers formed by opposing CER headgroups; only a single peak is observed for the extended conformation, occurring roughly at the center between these two peaks, rather than preferentially aligning with one of the layers of hairpin CER headgroups.

Lipid head groups in the SC form a network of intermolecular hydrogen bonds, which have been shown to affect the stability and permeability of SC lipid membranes.^65^ As such to further examine behavior, hydrogen bonding is examined. The average number of hydrogen bonds between each pair of lipid types has been characterized in Table 3. Lipid-lipid hydrogen bonds are measured separately for those in the interior of the membrane and at the exterior (i.e., in the leaflets in contact with water). Considering first the overall number of hydrogen bonds, scaled by the number of lipids in each region (i.e. interior or exterior), we observe that lipids in the interior have approximately 3 times the number of lipid-lipid hydrogen bonds than the exterior, on average, with just over one hydrogen bond per lipid in the interior. We would expect interior lipids to have more hydrogen bonds, given that there can be both intra- and inter-layer lipid-lipid hydrogen bonding; the exterior layer only has intra-layer lipid-lipid interactions due to the interface with water (lipids in the exterior have 2.00 ± 0.04 hydrogen bonds per lipid with water). Isolating an interior leaflet to only capture intra-layer lipid-lipid hydrogen bonding reveals 0.584 ± 0.023 hydrogen bonds per lipid, slightly larger than 0.396 ± 0.020 seen in the outer leaflet. As such, this shows that roughly half of lipid-lipid hydrogen bonds in the interior occur between lipids in separate leaflets (i.e., where leaflet headgroups are in contact). Focusing on the number of hydrogen bonds containing CERs, on average, each CER in the inner leaflets will have roughly one hydrogen bond with another CER, in contrast to the outer leaflet where one out of every 2 CERs will have a hydrogen bond with other CERs. Far fewer hydrogen bonds exist between CER and CHOL, where for inner leaflets only 1 in every 4 CERs will have a hydrogen bond with CHOL, and this number is halved in the outer leaflets. CER and FFA in the inner leaflets are likely to have just under one hydrogen bond per CER, although this number is cut by ∼1/3 when considering the outer leaflets. Virtually no CHOL-CHOL hydrogen bonding is observed, regardless of its location in the inner or outer leaflets. Hydrogen bonds are found between FFA and CHOL, where, again, the outer leaflets have ∼1/3 the number of bonds than the inner leaflets. When considering FFA, there are virtually no FFA-FFA hydrogen bonds in the outer leaflets, as compared to approximately 1 hydrogen bond for every 2 FFA lipids. Isolating only intra-layer hydrogen bonding between FFA lipids, we find 0.113 ± 0.012, and thus the bulk of the hydrogen bonding occurs between FFA lipids in separate leaflets (a factor of ∼3 times more inter-layer than intra-layer). Table 4 considers the fraction of hydrogen bonds formed by extended and hairpin conformations separately in the inner leaflets. When considering CER-CER hydrogen bonds, the largest fraction of hydrogen bonds are between extended and hairpin, followed closely by hairpin-hairpin. Only 15% of CER-CER hydrogen bonds form between extended lipids. When considering hydrogen bonding between the CERs and with CHOL and FFA, approximately 35% of the hydrogen bonds are with extended CERs, matching the total fraction of extended CERs in the system, suggesting that the hydrogen bonding per conformation are proportionate to their fraction in the system. Due to the differences in hydrogen bonding in the inner leaflets versus the interfacial leaflets, as well as the strong influences of hydrogen bonding on barrier properties, one would expect that permeability measurements through bilayer and self-assembled multilayer systems would yield differing results. Because of this, one should be cautious when interpreting simulated permeability measurements of bilayers and comparing them with experimental permeabilities.

**Table 3:**
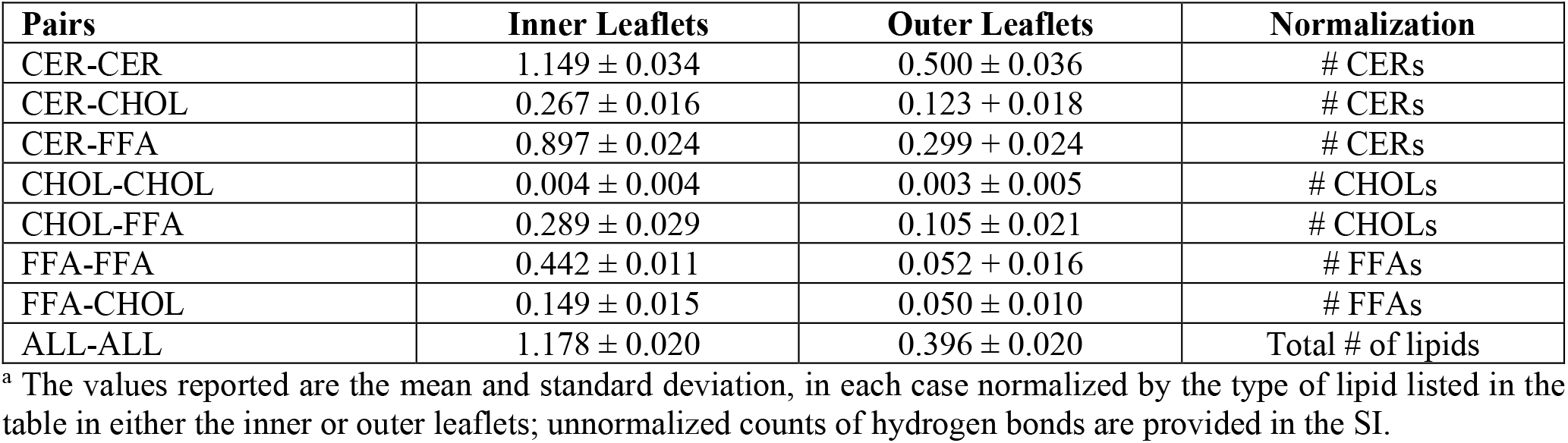
Normalized Number of Lipid-Lipid Hydrogen Bonds by Pairs of Molecule Types^a^

**Table 4:**
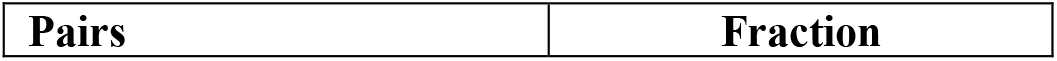
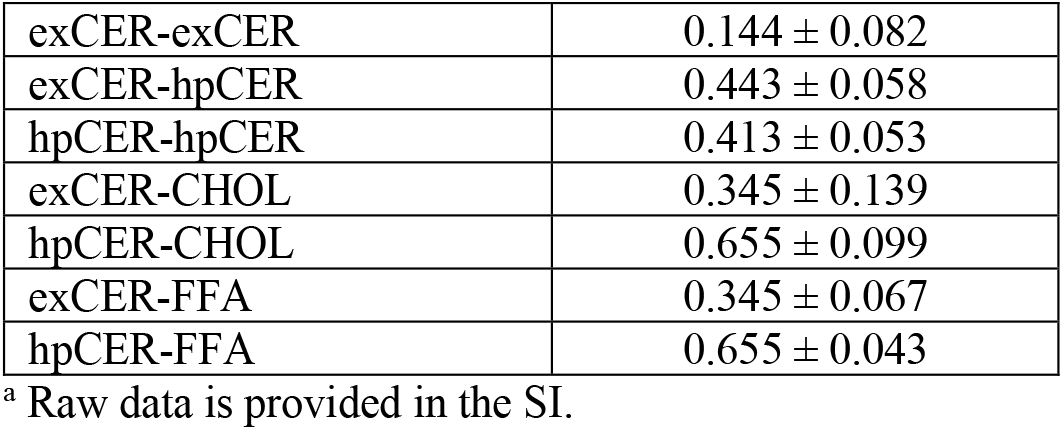
Fraction of Lipid-Lipid Hydrogen Bonds by Pairs of Molecule Types comparing hairpin and extended CERs in the Inner Leaflets^a^

## Conclusions

In this work, we present a novel CG forcefield for CHOL, derived using the MS-IBI method,^40^ which is compatible with our previously developed MS-IBI-based CG forcefields for CER NS and FFA.^38^ Together, these provide a set of force fields that can be used to examine systems exhibiting SPP-like ordering. In this work, the CHOL-CHOL self-interactions and CHOL-CER and CHOL-FFA cross interaction potentials were optimized using a diverse set of atomistic target states ranging from bulk fluids to pure bilayers to mixed bilayers, finding close agreement between the structural properties of CG and atomistic systems. The CHOL force field was further validated by comparing bilayer structural properties of preassembled atomistic, preassembled CG, and self-assembled CG structures for mixtures of CHOL with CER NS and FFA, not used as target states for optimization, again finding good agreement and thus indicating that the CG model accurately reproduces atomistic bilayer structures. To probe more realistic structures, self-assembled multilayer structures were simulated for mixtures of CER NS C24:CHOL:FFA C24 1:0.5:1, comparing bilayer structures to 4-leaflet and 6-leaflet systems. While only small differences in the average APL, tilt angle, S2 order parameter, and height were observed, a significant number of extended CERs (∼35-36% or total CERs in interior layers) are present in the inner layers of the multilayer structures, which affects the morphology and arrangement of lipids in-plane as well as lipid-lipid interactions spanning adjacent layers. Since experimental studies, such as neutron scattering and FTIR, average properties over dozens of lamellae, the results are largely dependent on the structure of interior layers, and thus the central bilayer of 6-leaflet structures provides the best comparison to experimental measurements. Self-assembled 6-leaflet CER NS C24:CHOL:FFA C24 1:X:1 systems were simulated, where the CHOL molar ratio, X, ranged from 0 to 1, demonstrating that interdigitation, fluidity, and number of packing defects increase with increasing CHOL content. The results also indicate that the size of FFA and CER clusters reduces with increasing CHOL, suggesting that CHOL may improve the miscibility of these two components.

To further compare with experiment and examine atomistic level properties, a simplified reverse-mapping procedure was used to generate an atomistic 6-leaflet CER NS C24:CHOL:FFA C24 1:0.5:1 system, using the associated CG self-assembled structure as reference. This reverse-mapped structure preserved the conformation of CERs (i.e. hairpin or extended) and in-plane morphology of the associated self-assembled structure. Compared to previous work,^22,23^ which used only pre-assembled bilayer systems, the simulated NSLD curves using the inner layer of the reverse mapped structure showed much closer agreement with experimental NSLD curves from a model SPP system,^15^ highlighting the importance of considering multi-layer structures. Furthermore, it was observed that approximately half of the lipid-lipid hydrogen bonds in the inner portion of the multilayer system exist between lipids in separate leaflets (i.e., inter-layer hydrogen bonding), which may differentiate barrier properties through the inner layers from the outer layers or bilayer structures. This multiscale approach allows for the exploration of atomistic self-assembled multilayer configurations that capture both hairpin and extended CER conformation and unbiased morphologies, rather than arbitrarily arranged pre-assembled configurations, allows for the simulation of systems that are more closely representative of experimental. With the parameterization of additional CER headgroups for the CG model, this approach can be applied to other experimentally realized model systems^10,16,62^ and provide molecular-level explanations for the observed experimental behavior.

## Acknowledgments

The project described was supported by Grant Number R01AR072679 from the National Institute of Arthritis and Muscoskeletal and Skin Diseases. D.X. also acknowledges support from the National Science Foundation through grant number DMR-1852157. Computational resources were provided by the National Energy Research Scientific Computing Center (NERSC) operated under Contract DE-AC02-05CH11231and the Advanced Computing Center for Research and Education at Vanderbilt University.

## Supporting Information

### Atomistic Simulation Methods

All atomistic simulations are performed with the GROMACS simulation engine^49^ using a 1 fs timestep. Van der Waals interactions were cut off at 12 Å, and set to smoothly decay to zero between 10 -12 Å. The particle-mesh Ewald summation method^66^ was used to handle long-ranged electrostatic interactions, with a real-space cutoff of 12 Å. The Nose-Hoover thermostat^50^ with a time coupling constant of 1 ps was used to control the temperature. The Parinello-Rahman barostat was used for pressure control, with a coupling constant of 10 ps and a compressibility of 4.5e-5 bar^-1^.^51^ For constant-pressure bulk fluid simulations, pressure was controlled isotropically. For constant-pressure bilayer simulations, pressure was controlled separately in the bilayer normal and lateral directions. All lipids were modeled with the atomistic CHARMM36 force field^67^ supplemented by parameters for the CER NS headgroup where necessary.^68^ Water was modeled with the TIP3P water model.^69^

### Simulation of Atomistic Target States

A variety of different atomistic simulations were performed to gather target data. Pure, bulk fluid CHOL simulations contained 64 CHOL molecules, and are run at four different statepoints, as listed in Table S1. CHOL-water systems contained 1250 water molecules, and the corresponding number of CHOL molecules to give the concentrations listed in Table S1. Bulk fluid lipid mixtures contained 64 copies of each lipid. These systems are initialized with lipids randomly distributed throughout a large simulation box and simulated for 5 ns at 800K to randomize the positions of the molecules. The system temperature is then reduced to the target temperature while condensing down to a realistic liquid density over 5 ns. For constant-pressure, bulk fluid systems, a Parinello-Rahman barostat using a coupling constant of 10 ps and a compressibility of 4.5e-5 bar^-1^. The systems are relaxed for 5 ns, before collecting data over 10 ns. For constant-volume, bulk fluid systems, the systems are relaxed at the target density for 5 ns before collecting target data over the final 10 ns. Mixed-lipid bilayers are taken from simulations described in Moore et al. 2018.^22^ The dehydrated bilayers are constructed by removing the water from the corresponding bilayer structures, replicating the bilayer in the z direction, and relaxing with the random walk molecular dynamics (RWMD) protocol described in Moore et al. 2018.^22^ The CHOL-FFA C16:0 monolayer is constructed by placing an equal number of CHOL and FFA molecules on a hexagonal lattice with an area per lipid (APL) of 33.1 Å^2^. The atoms corresponding to the headgroups of each lipid are fixed, and the system relaxed for 10 ns at 405 K. The system temperature is then lowered to 305 K over the course of 20ns and allowed to relax for 10 ns. Target data is then collected for 50 ns.

Simulations of droplets on surfaces are performed for tuning various interactions in the CG force field. These include a water droplet on the tail side of a CHOL monolayer surface, and CER headgroups on the headgroup side of a CHOL monolayer. These systems are initialized by carving a sphere out of a bulk fluid of the droplet species (i.e., either water or CER headgroups), and placing it on the specified surface. The droplet is allowed to relax; data is collected from the final 5 ns after the contact angle had reached a steady-state value.

**Table S1:**
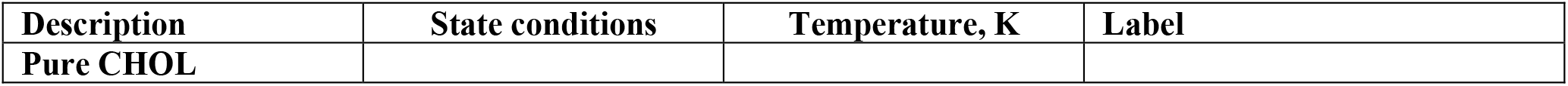
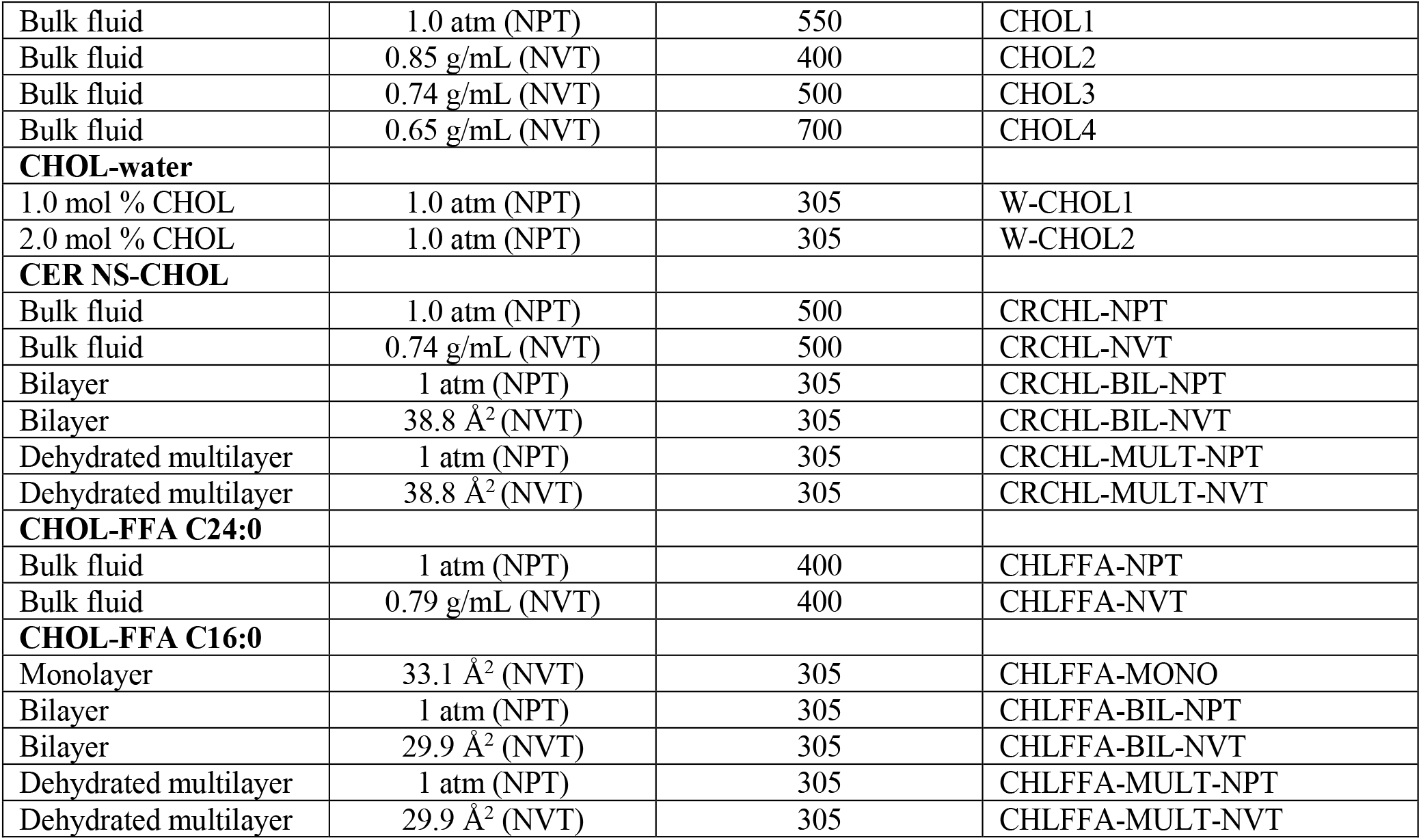
Target states used in the MS IBI optimization of CHOL self- and cross-interactions

**Table S2:**
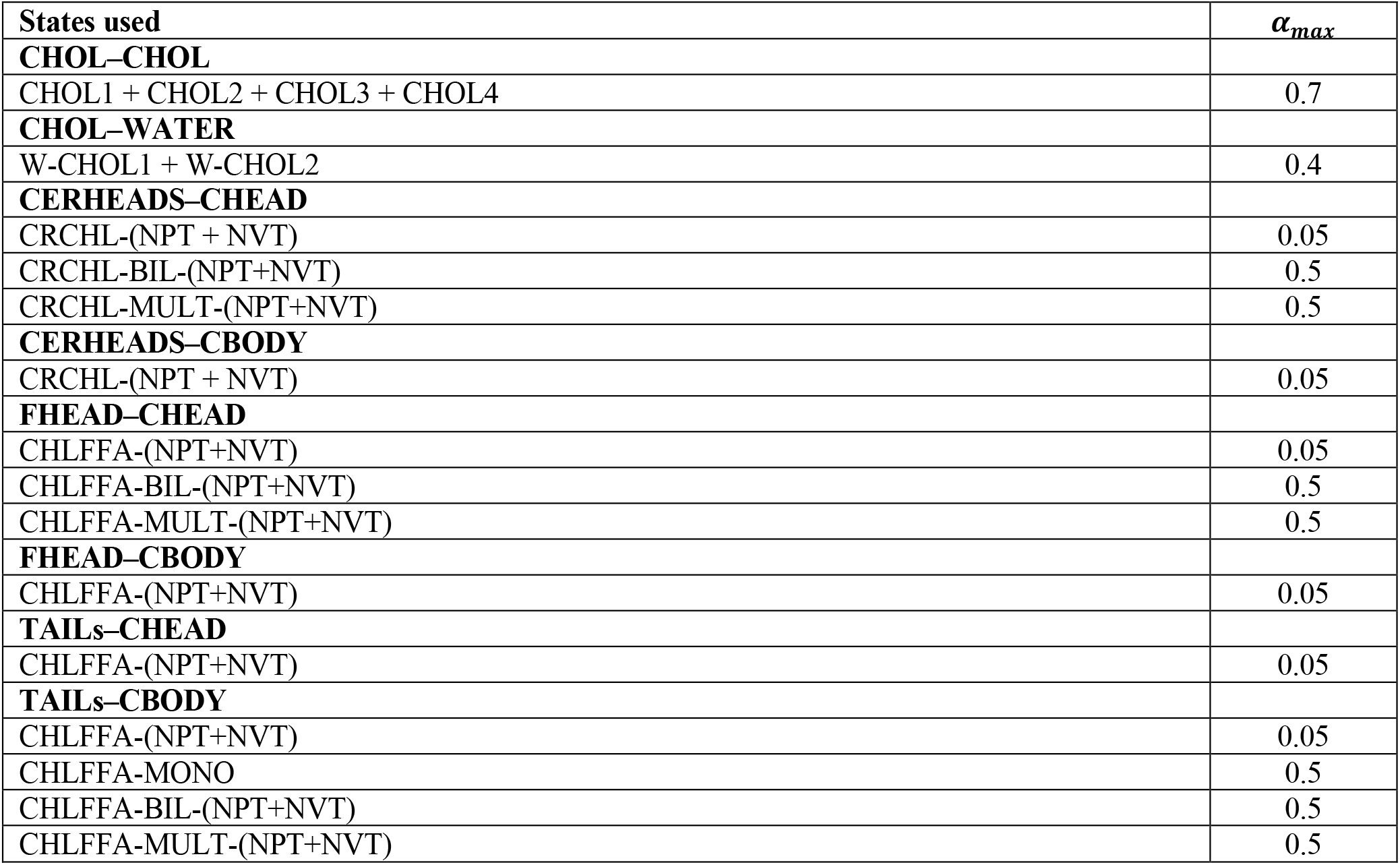
States used to optimize each pair potential and the weight α_*max*_ given to each state. Pairs are denoted by bold text, with the states used to optimize a given (set of) interaction(s) listed below the interaction name. The state condition names correspond to the label given in Table S1.

## Coarse-Grained Simulation Methods

All CG simulations are performed using the GPU-accelerated HOOMD-Blue simulation engine with a 10 fs timestep.^52^ Systems using the constant-temperature (NVT) ensemble use a Nosé-Hoover thermostat with a coupling constant of 1 ps,^54,70^ and constant-pressure (NPT) CG simulations use the Martyna-Tobias-Klein barostat-thermostat.^54,71,72^ Three distinct sets of CG simulations are performed for force field optimization, force field validation, and self-assembly.

### Force Field Optimization and Validation

The first set is run during the MS-IBI optimizations to generate the RDF for each iteration. These use the final configuration from the corresponding atomistic simulation for the initial configuration, except configurations are replicated twice in each direction for bulk systems and three times in each lamellar lateral direction for the lamellar systems. These simulations use the force field derived in Moore et al. 2018,^30^ except for interactions involving CHOL beads, which are optimized during this stage. Topological constraints, i.e., bonds/angles, are determined following the procedure in Milano, et al.^73^ Briefly, distributions of the bond distances and/or angles are collected from atomistic simulations mapped to the CG level, with harmonic bond/angle parameters fit to best reproduce these distributions; 11 bond and 23 angle types are present in the CG model, and each have unique stiffnesses and equilibrium separations, where the parameters used for each bond/angle type are provided in the Table S3 of the SI.

**Table S3:**
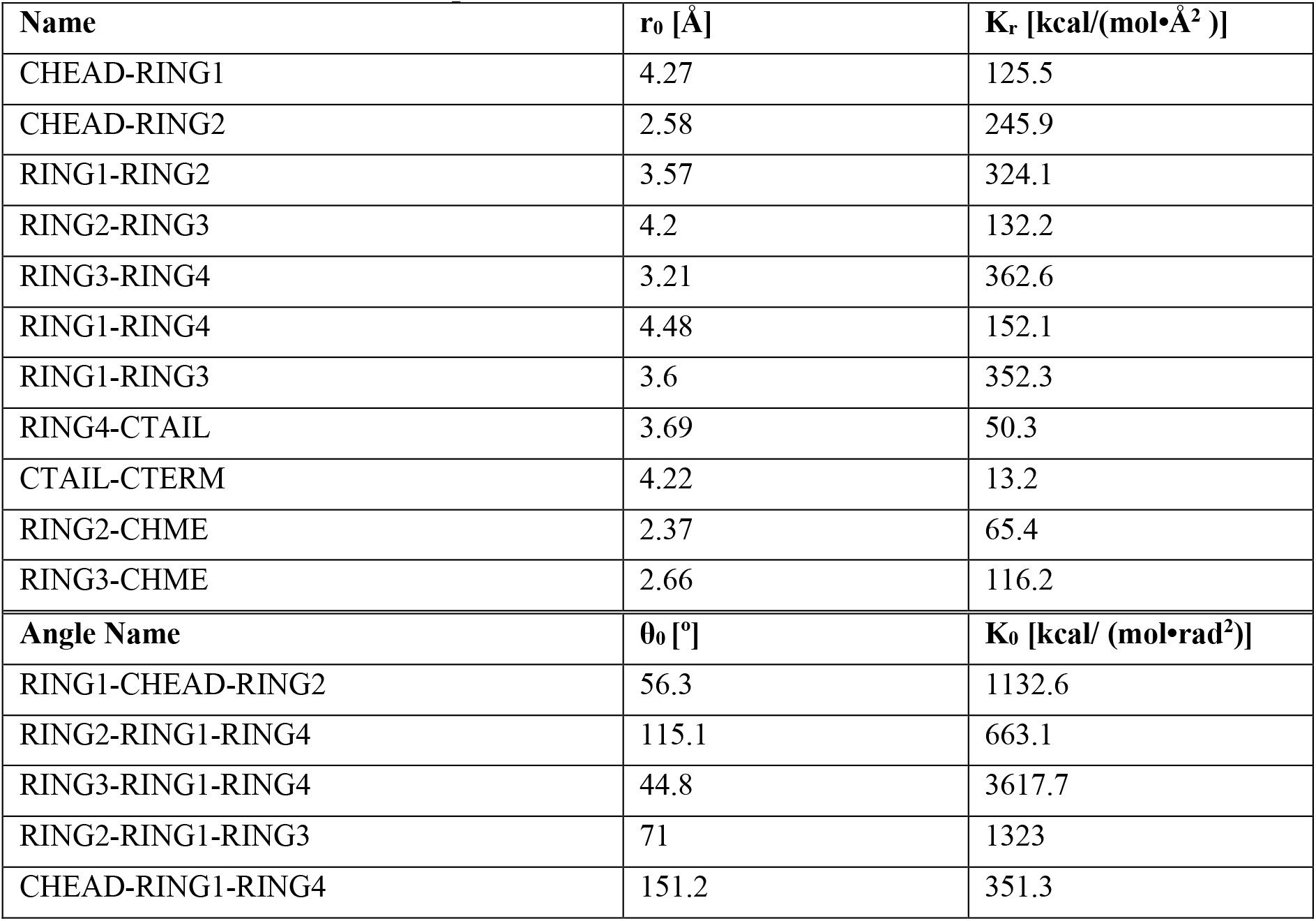
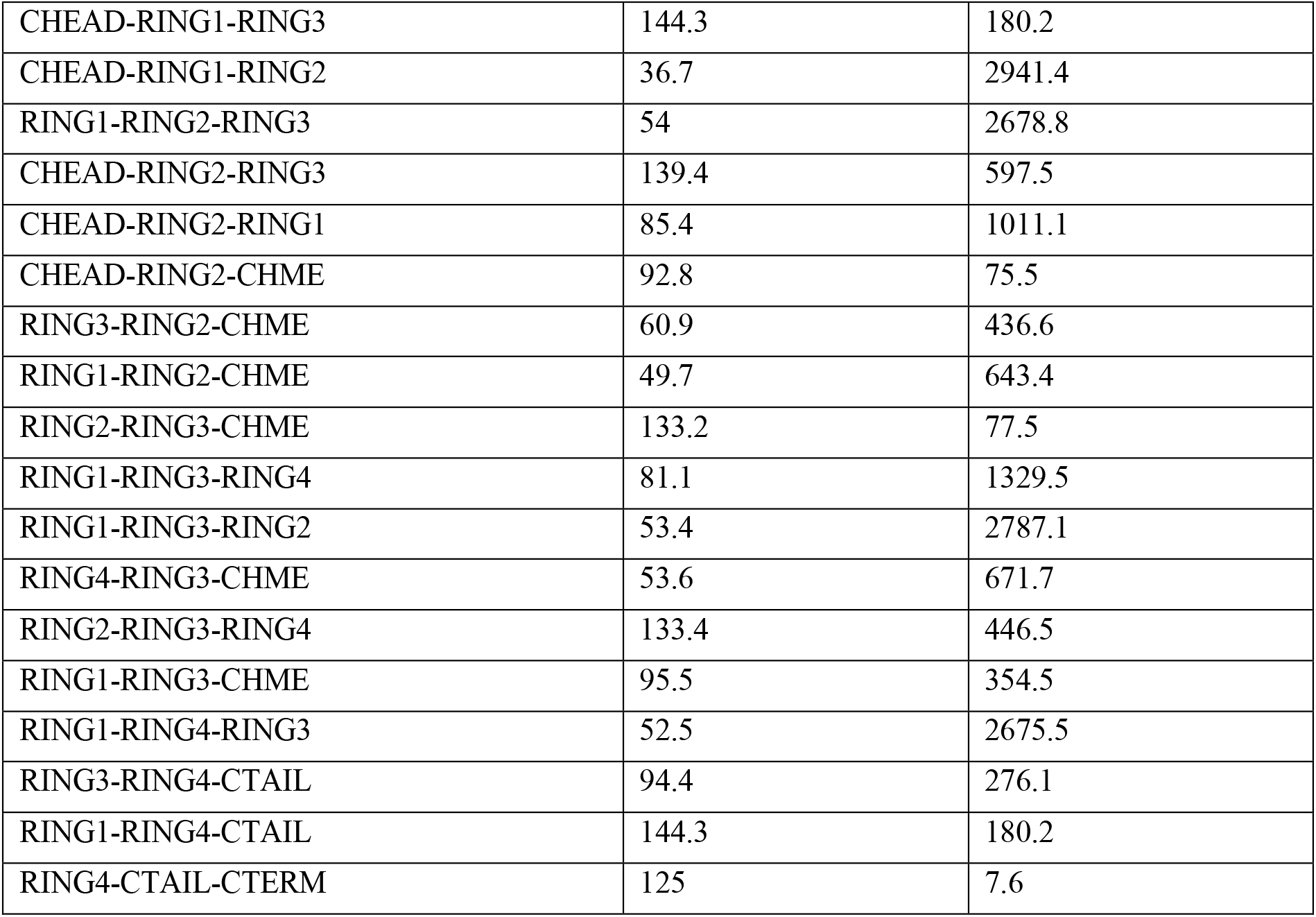
Bonded interaction parameters for CHOL.

The next set of simulations use the newly derived CHOL interactions to validate the force field and tune, where necessary, specific interactions. These include the surface wetting simulations used to tune specific interactions. These simulations are performed analogously to their atomistic counterparts, except systems are simulated for longer times to ensure stability of the structures. If it is found that the force field needs to be tuned based on the surface wetting behavior, these simulations are repeated with the modified force field. This cycle is repeated until no further modifications are required (i.e., the CG wetting behavior matches the atomistic wetting behavior).

### Self-Assembly Simulations

Finally, after the CG force field is modified and validated, the self-assembly of mixtures of the various lipids is simulated. Systems used for self-assembly are initialized in disordered configurations, with the lipids and water in separate domains, and the lipid phase spanning the box in the *x* and *y* directions, but not in *z*; this setup is used to ensure that the lipid phase only forms a 2D periodic structure. The size and shape of the simulation box is initialized with a target density and aspect ratio amenable to the formation of a periodic structure (i.e., estimated from the pre-assembled configurations). In all systems, the initial density of the water phase is set to 1.0 g/cc, and the lipid density is set to 0.8 g/cc, which is approximately the expected density based on NPT atomistic simulations of the bulk fluid state at 305 K. Given the lipid density and composition, the lipid volume is specified, and the cross-sectional area of the box is set based on the estimated APL and number of leaflets. To remove high-energy overlaps, very brief NVE simulations are first performed, with a limit on the displacement of each bead at each timestep (0.1 Å). Next, a brief simulation at 305 K and 1 atm with an isotropic barostat is performed to allow the total density of the system to equilibrate while keeping the aspect ratio of the box constant. After density equilibration, the barostat is turned off, and the systems undergo an expansion/compression process, as described in Moore et al. 2018.^30^ Following the expansion/compression, the semi-isotropic barostat is turned on, and the system annealed from 450 K to 305 K over 100 ns to remove any defects formed during the expansion/compression phase. After annealing, the simulation is continued for at least another 200 ns, and data for analysis is collected for at least 100 ns.

## Analysis Methods

### CG mapping

CG mapping is used to convert atomistic trajectories to the CG level. This is done using the procedure outlined in Moore et al.^38^ Briefly, the CG beads are placed at the center of mass of their constituent atoms for lipid molecules. For water, a k-means algorithm is used to identify clusters of 4 water molecules (i.e. k = N_waters_ / 4) for which a CG water bead is placed at the center of mass of the water molecules in each cluster. The python script used for CG mapping is available on GitHub (https://github.com/uppittu11/cg_mapping).

### Structural properties

The APT, tilt angle, S_2_, and fraction of extended CERs are calculated according to the methods outlined in the SI of reference ^38^. The area per tail is calculated by dividing the number of tails in the leaflet by the cross-sectional area of the simulation box and multiplied by the cosine of the average tilt angle of the leaflet to project the area onto a plane perpendicular to the direction of the tail. The tilt angle is defined as the angle between the tail director vector and the z-axis, where the director is the eigenvector corresponding to the minimum eigenvalue of the inertia tensor. S2 is calculated by taking the eigenvalue of the nematic tensor (see reference 74 for more details). The fraction of extended CERs is calculated by measuring the angle between the COM of the acyl chain, the amide group, and the COM of the sphingosine chain. If this angle is *≥* 90º the CER is considered to be in the extended conformation and if this angle is < 90º the CER is considered to be in the hairpin conformation. Finally, the bilayer thickness is taken as the peak-to-peak distance in the mass density profile. For multilayer systems, the height for each layer is calculated (i.e. the distance between each set of adjacent peaks).

### Coordination Number

In order to get the coordination numbers between the different lipid tails (as in Figure 7) a trajectory of the lipid tails is generated by placing particles at the center of mass of each lipid tail (herein referred to as the “COM trajectory”). RDFs between lipid tail types is determined by calculating the RDFs between the sets of particles representing each of these tails. Finally, coordination number is taken as the area under the curve of the first peak of the calculated RDF. This analysis is done separately for each leaflet.

### Lipid Clustering Analysis

Clustering is calculated using Freud,^56^ which defines clusters as a network of nearest neighbors for each snapshot of the COM trajectory. This analysis is done separately for each leaflet.

### Voronoi diagrams

The lipid tail COMs are extracted for each leaflet. The coordinates of the COMs are projected onto the x-y plane. 2D Voronoi tessellations are determined for the resulting coordinates using Freud^56^ and Voro++ packages.^75^ Each polytope is labeled with either the lipid tail type or number of sides and visualized using Matplotlib.^76^

## CG Force Field Optimization

### Pure CHOL and CHOL–WATER

Overall, excellent agreement was found between the target and CG RDFs. In any given CHOL system, there will be far more RING-RING interactions, so this pair is used as a representative pair for the force field; the optimized pair potential, along with the target and CG RDFs at two of the states used in the optimization, are shown in Figure S1. As expected based on the excellent agreement in the RDFs, the CG and atomistic models show good agreement in the density of the NPT target state, with values of 0.732 ± 0.005 g/mL (CG) and 0.73 ± 0.01 g/mL (atomistic).

**Figure S1:**
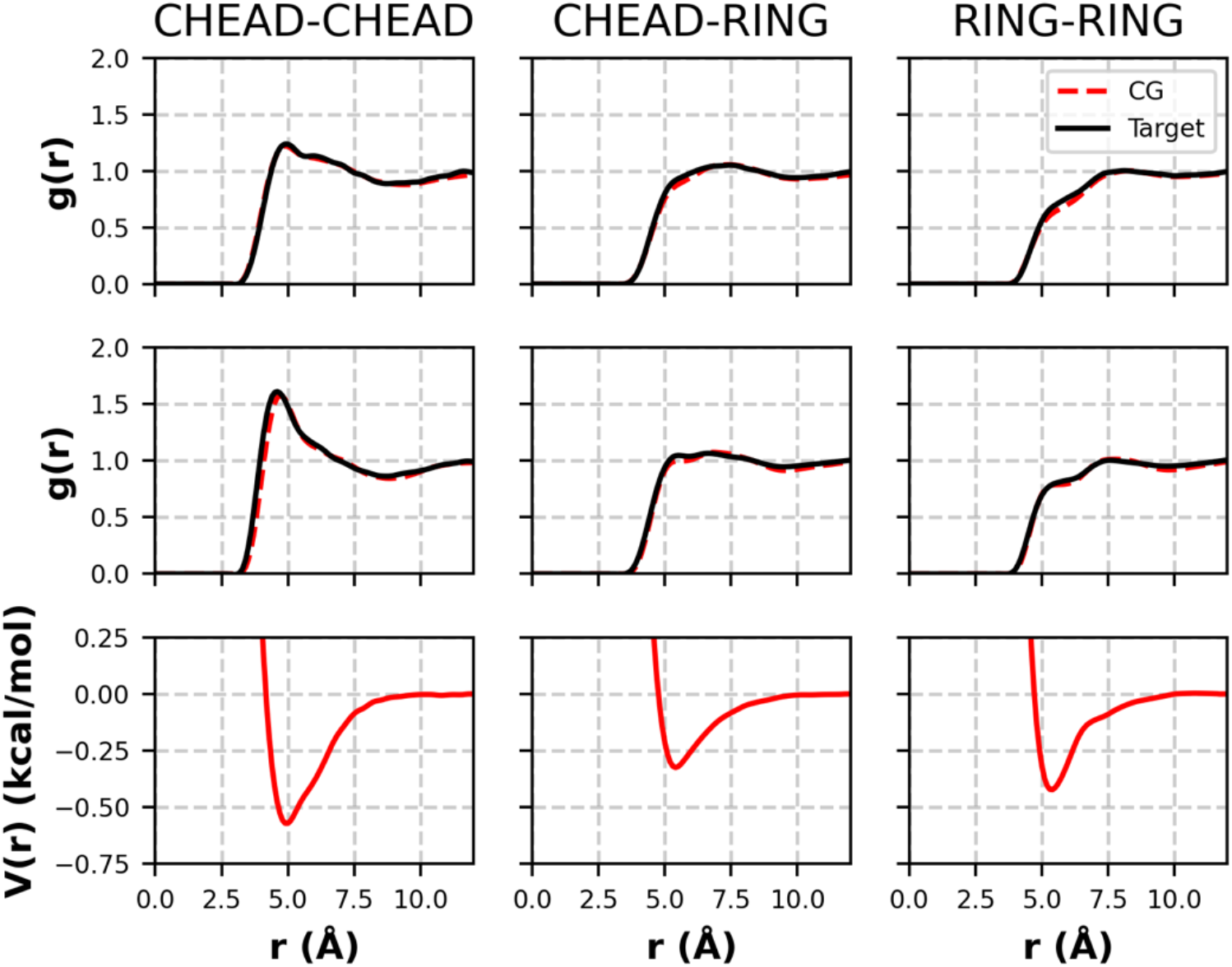
RDFs and pair potential from the pure CHOL force field optimization. Top: target(black) and CG (ref) RDFs from the 400K NVT state; middle: target and CG RDFs at the 550 K NPT state; bottom: pair potential that yields the CG RDFs above.

Overall, good agreement was found between the CG and target RDFs for interactions between CHOL and water. However, as discussed in Moore et al., RDF matching does not ensure that the partitioning behavior will be replicated.^38^ Therefore, to test the hydrophobicity of the CHOL beads, the wetting behavior of water on a CHOL monolayer was investigated. Using the optimized force field, the water tended to spread on the CBODY surface, forming a contact angle of 82º. This result indicates that the CBODY surface is more hydrophilic than the corresponding atomistic surface, which gave a contact angle of 127º. The CBODY–WATER interactions were scaled to be less attractive, and a scaling value of 0.4 was found to give good agreement with the atomistic contact angle. In contrast, water was found to readily wet the CHEAD surface, and therefore the CHEAD–WATER interaction was not modified from its optimized form.

After the CHOL–CHOL and CHOL–WATER pair potentials were optimized and modified, they were used in the optimization of the mixed-lipid interactions. Two sets of pair potentials were optimized: CHOL–FFA and CER–CHOL. Although CER and FFA tails both consist of TAIL beads, the interactions between the TAIL and CHOL beads were only optimized with data from CHOL–FFA mixtures for simplicity. As in Moore et al. (2016),^38^ interactions were optimized using target data from states where specific interactions are thought to play an important structural role, e.g., target data from lamellar states is not used in the interactions between lipid headgroups and tails. In all cases, more weight (through the α term of the MS IBI pair potential update formula) was given to the lamellar states (α = 0.5) than the bulk fluid states (α = 0.05). The results of these sets of optimizations are discussed separately below.

### CHOL–FFA

The TAIL–CBODY RDFs are generally reproduced with a fairly high degree of accuracy, which is important since there will be more of these interactions in any given system containing CHOL. The TAIL–RING interactions are likely dominant in these system because of the number of each of these beads, therefore, this pair is a good representation of the TAIL–CBODY interactions. The TAIL–RING RDFs are shown in Figure S2, where it can be seen that the target RDFs are well-reproduced with the CG model.

**Figure S2:**
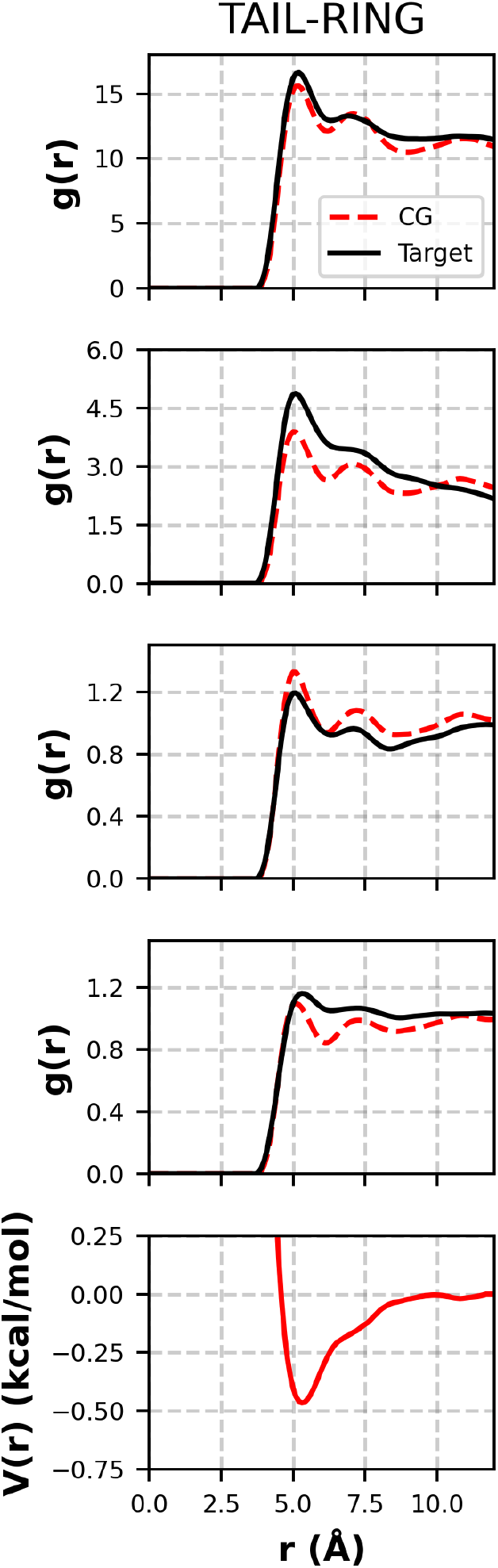
TAIL–RING RDFs and pair potential. From top to bottom: RDFs from the monolayer state; RDFs at the NPT bilayer state; RDFs at the stacked bilayer state; RDFs at the bulk fluid NPT state; and the pair potential that yields the CG RDFs above.

The CHEAD–FHEAD RDFs are decently captured at the lamellar states, as shown in Figure S3. Interestingly, the best fit is found at the stacked bilayer state, even compared to the bilayer state. This is likely a result of the relative locations of the first peaks: at the bilayer state, the first peak in the RDF is at a larger separation compared to the stacked bilayer and bulk states. To compensate, the pair potential shifts the first RDF peak to smaller separation values. Like in previous optimizations, the fits at the lamellar states are better than at the bulk states as a result of the lamellar states being given a larger weight in the optimization process. The RDFs between the mixed hydrophilic–hydrophobic beads, e.g., TAIL–CHEAD (shown in Figure S3), are captured with less accuracy than in previous Moore et al. (2016).^38^ This may be a result of the shape of CHOL, which is more complex than the simpler chain-like CER and FFA molecules. Despite the discrepancies, the density of the bulk fluid NPT state was found to be 0.81 ± 0.03 g/mL, which is in good agreement with the atomistic value of 0.79 ± 0.01 g/mL. Additionally, in a lamellar configuration, there are likely to be very few of these types of interactions. Therefore, these deviations in the RDF are deemed acceptable considering the target objective of this force field.

**Figure S3:**
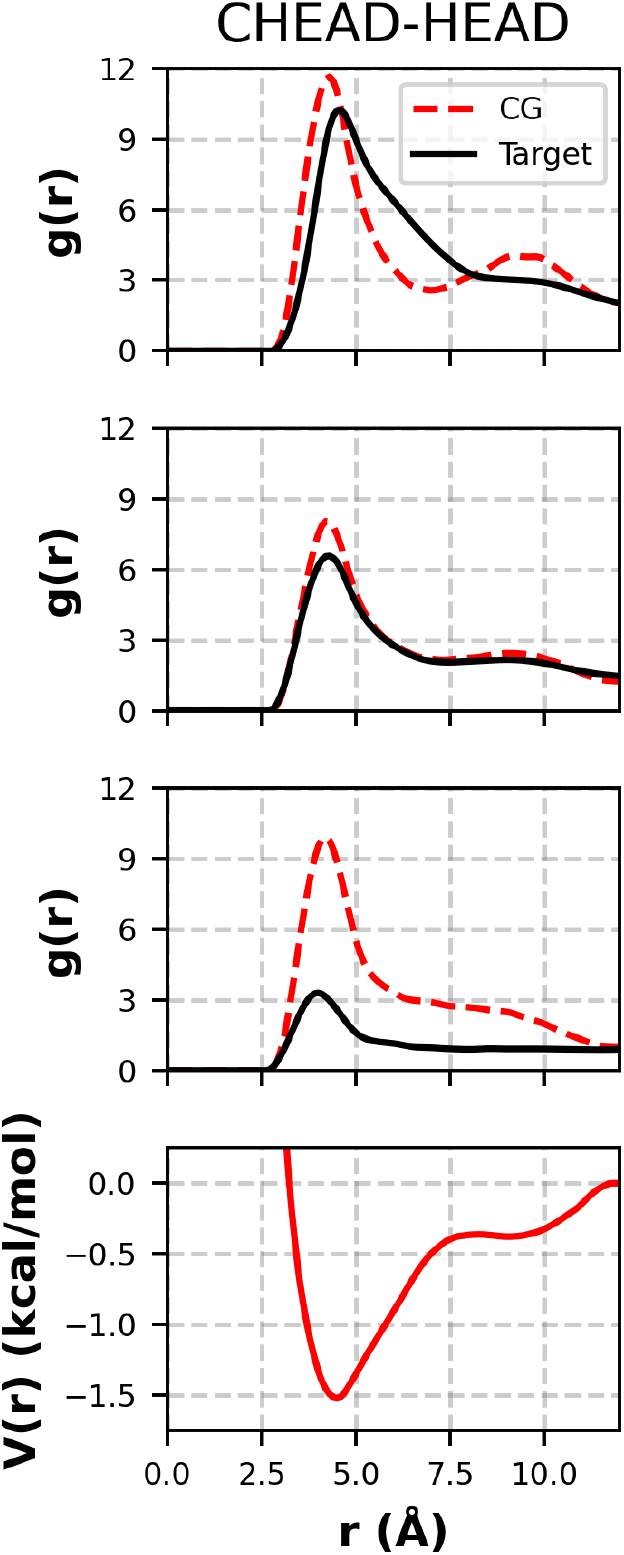
CHEAD–FHEAD RDFs and pair potential. From top to bottom: RDFs from the bilayer state; RDFs from the stacked bilayer state; RDFs from the bulk fluid NPT state; and the pair potential that yields the CG RDFs above.

### CER NS–CHOL

Only the CERHEADs–CHEAD and CERHEADs–CBODY interactions are considered here, since the TAILs–CHOL interactions were optimized using target data from CHOL–FFA systems. In general, as shown in Figure S4, the CERHEADs–CHEAD interactions are accurately reproduced at the lamellar states.

**Figure S4:**
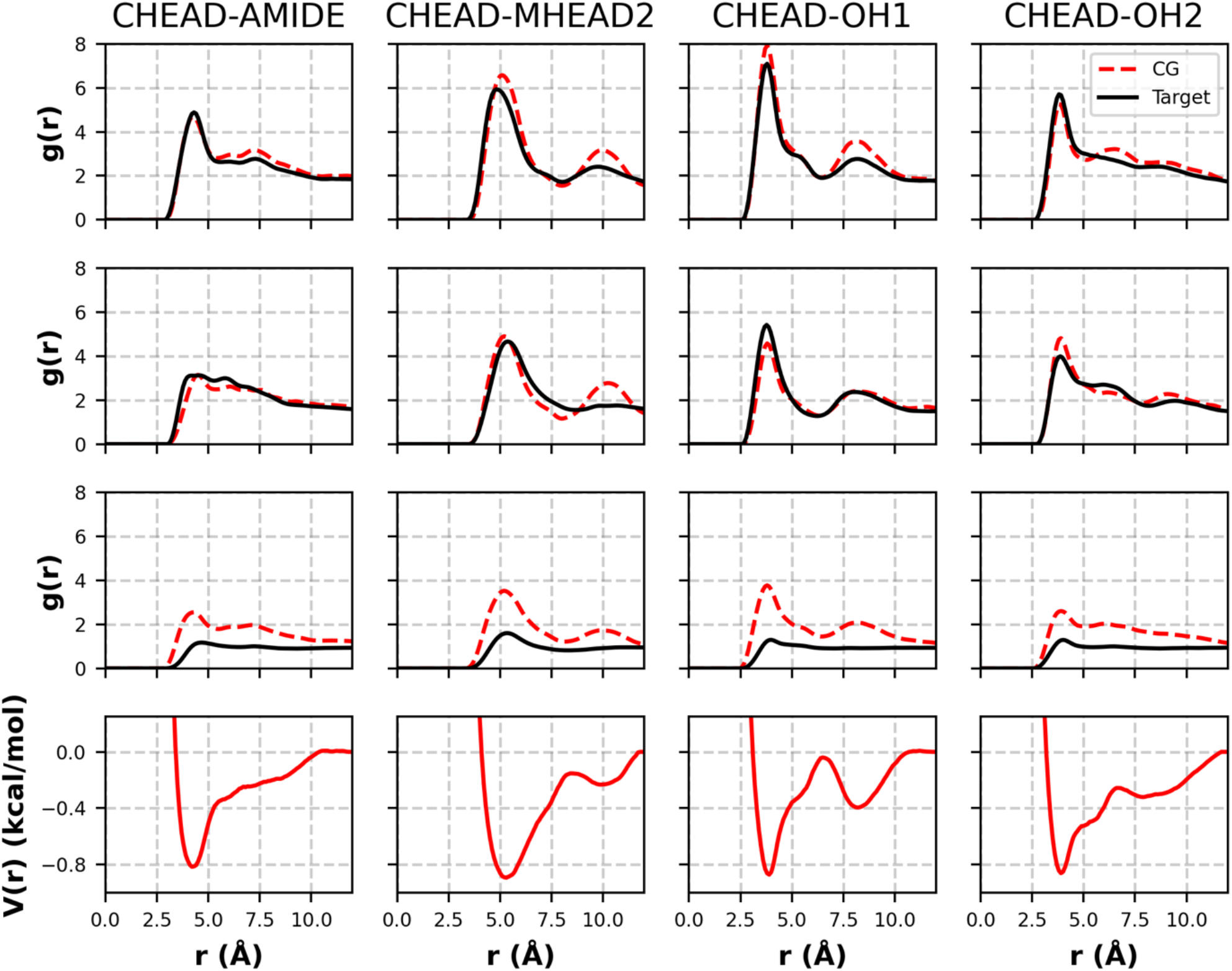
CERHEADS–CHEAD RDFs and pair potential. From top to bottom: RDFs at the stacked bilayer state; RDFs at the bilayer state; RDFs at the bulk fluid NPT state; and the pair potentials that yield the RDFs above. Target RDFs are shown by solid black lines, and CG RDFs are shown as the dashed red lines. Each column corresponds to a given pair, listed at the top of that column.

As seen previously, the fit is much better at these states compared to the bulk fluid states, as a result of the higher weight given to the lamellar states in the optimization process. Despite the accurate RDFs and stable lamellar phases, preliminary self-assembly simulations found that CER– CHOL mixtures did not form bilayer or multilayer structures, in contrast to the stable bilayers simulated with the atomistic model. To investigate, the wetting behavior of CER headgroups on a CHOL headgroup surface was examined. Here, the surface was a CHOL monolayer where the CTERM beads were fixed, and the droplet was composed of CER headgroups (i.e., no tails) and interacted with the CHEAD side of the monolayer. The CER headgroups fully wetted the CHEAD surface with the atomistic model. However, with the CG model, the CER headgroups tended to form a droplet on the surface, indicating an imbalance in the CERHEADs–CERHEADs and CERHEADs–CHEAD interactions. This imbalance was corrected by systematically making the CERHEADs–CHEAD interactions more attractive until the CER headgroups wetted the surface with the atomistic model. It was found that a scaling factor of 1.5 was required to give the correct wetting behavior; therefore, the derived interactions scaled by 1.5 were used for the final CG force field. The CERHEADs–CBODY interactions, which were optimized using target data from only the bulk fluid states, generally show a good match with the target RDFs.

## CG Model Validation

Several bilayer systems were self-assembled and compared to the atomistic simulation data from Moore et al. Three ratios of CER:CHOL:FFA and 5 ratios of CER NS C24:CER NS C16 were considered for a total of 15 distinct systems. Table S4 contains a list of all of the systems simulated and their compositions. Bilayers containing 1000 lipids and 10000 water molecules using these compositions were self-assembled from randomized initial configurations using the procedure outlined in the *Self-assembly simulations* section of the SI.

**Table S4:**
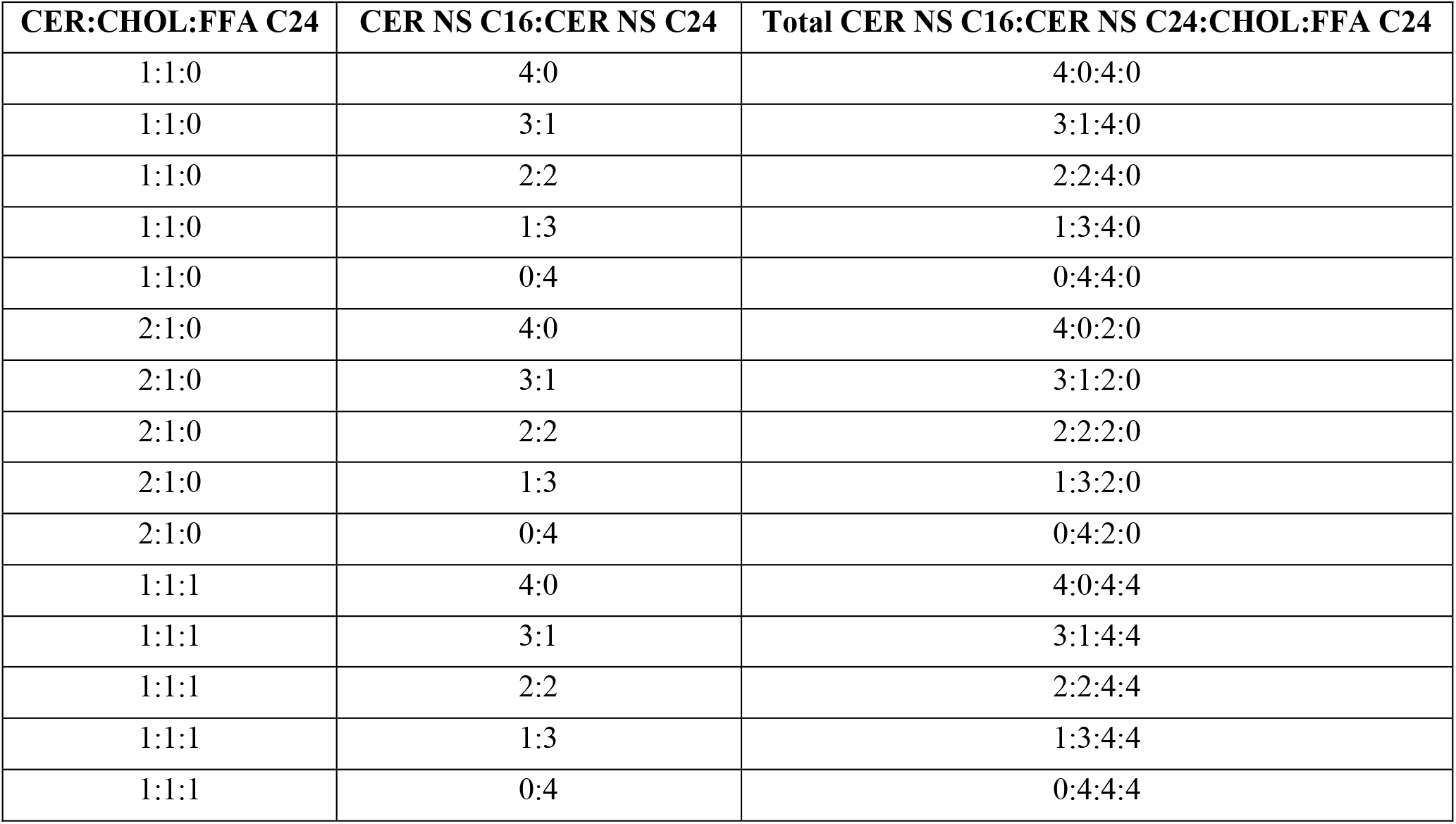
Molar ratios of systems used for validation.

Figure S4 compares structural properties of the CG self-assembled systems to the atomistic simulation results obtained by Moore et al.^22^ Prior to analysis, the atomistic trajectories are mapped to the CG resolution using the method described in the Analysis Methods section of the SI. The values in table S4 for CG systems are averaged from four replicate systems initialized from different randomized configurations. From these results, we can see close agreement between the CG and atomistic values. One should note that several CHOL molecules were “stuck” in the center of the atomistic bilayers simulated by Moore et al.^22^ This resulted in a slightly lower area per tail, due to the decrease in bulky CHOL molecules at the water interface, and increased the height, as the CHOLs in the center of the bilayer act as spacers between the two leaflets.

**Figure S4:**
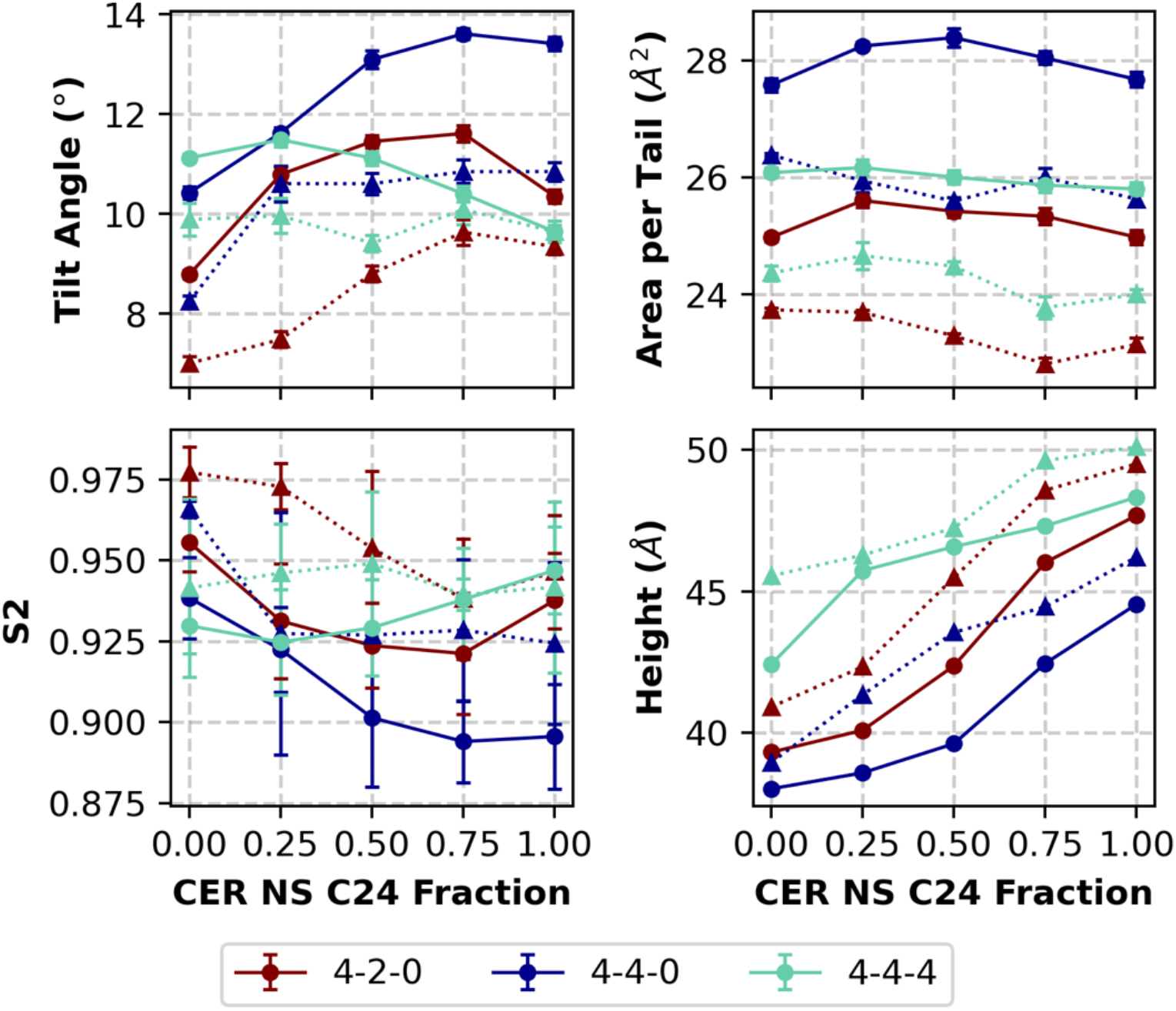
Structural parameters of CER NS:CHOL:FFA C24 systems with 4:2:0, 4:4:0 and 4:4:4 ratios, for which the ratio of CER NS C16:CER NS C24 is varied from pure CER NS C16 to pure CER NS C24 in five evenly spaced increments (i.e. 4:0, 3:1, 2:2, 1:3, and 0:4). Results from self-assembled CG bilayers are represented by circles and connected by a solid line, whereas results from preassembled atomistic bilayers (taken from Moore et al.) are represented by triangles and connected by a dashed line.

## Tail Angle Distributions for CER NS C24/CHOL/FFA C24 Multilayers with Varying CHOL content

A histogram of the angles between the sphingosine and acyl chains of interior CERs are plotted in figure S5.

**Figure S6:**
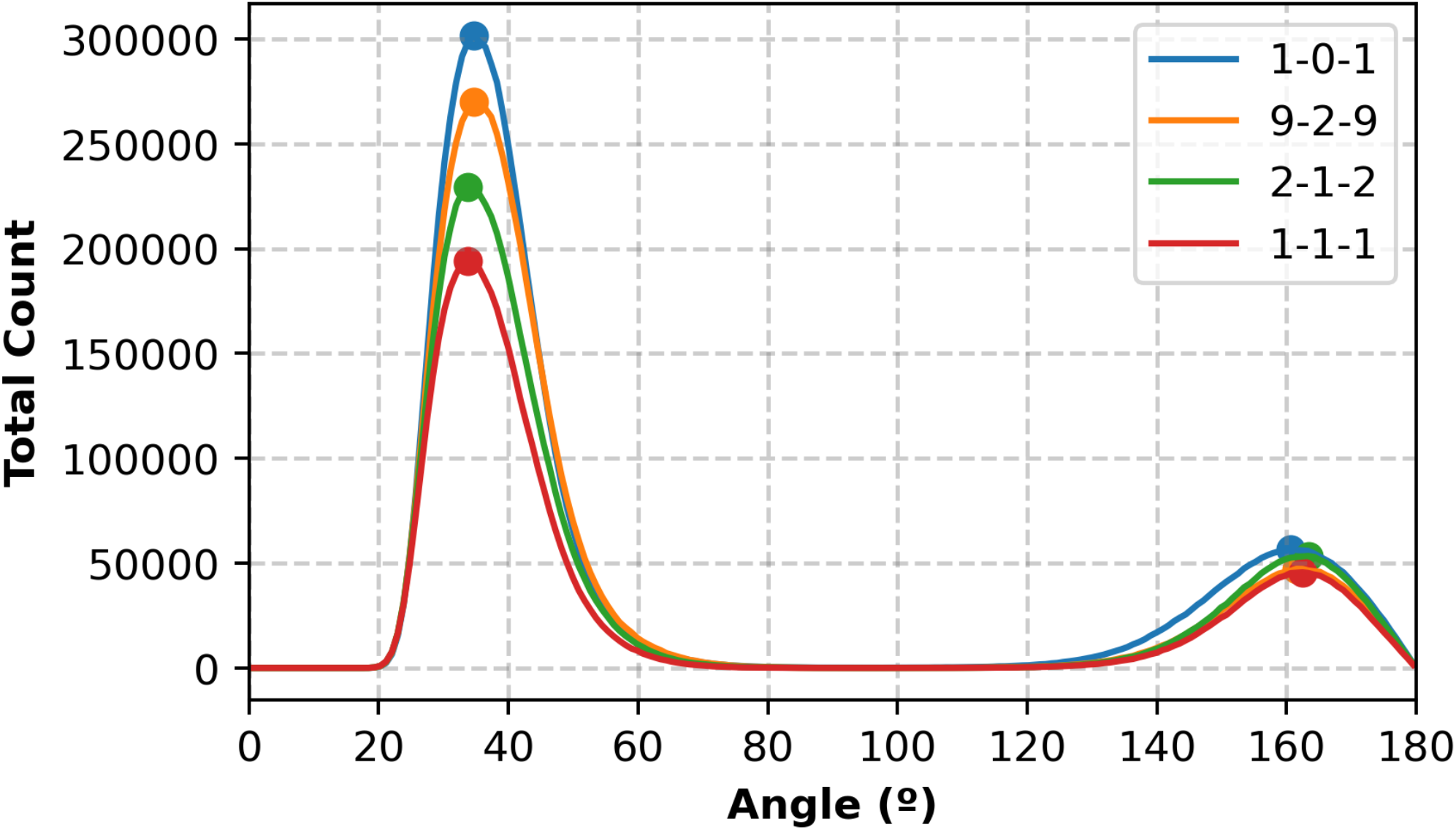
Histograms of the angle between the acyl and sphingosine chains of CER NS molecules in the interior of CER NS C24/CHOL/FFA C24 6-leaflet multilayers in 1:0:1, 1:0.2:1, 1:0.5:1, and 1:1:1 ratios.

## Voronoi Analysis of CER NS C24/CHOL/FFA C24 Multilayers with Varying CHOL Content

Voronoi analysis of the interior leaflets of the CER NS C24/CHOL/FFA C24 multilayers was conducted using Freud as described in the “Analysis Methods” section of the SI. Voronoi diagrams of the last frame of first trial for each lipid ratio is shown in figures S6. Each polygon is colored by the number of sides in figure S6. Figure S7 shows the average number of sides by lipid type. Here we can see that CHOL molecules have greater than 7 sides due to their bulky tail, whereas FFA molecules have less than 7 sides due to their narrow tail. While the average number of sides stays constant for CHOL as the CHOL content increases, more defects are introduced in the packing of the other lipid tails since the average number of sides deviates from 6.

**Figure S6:**
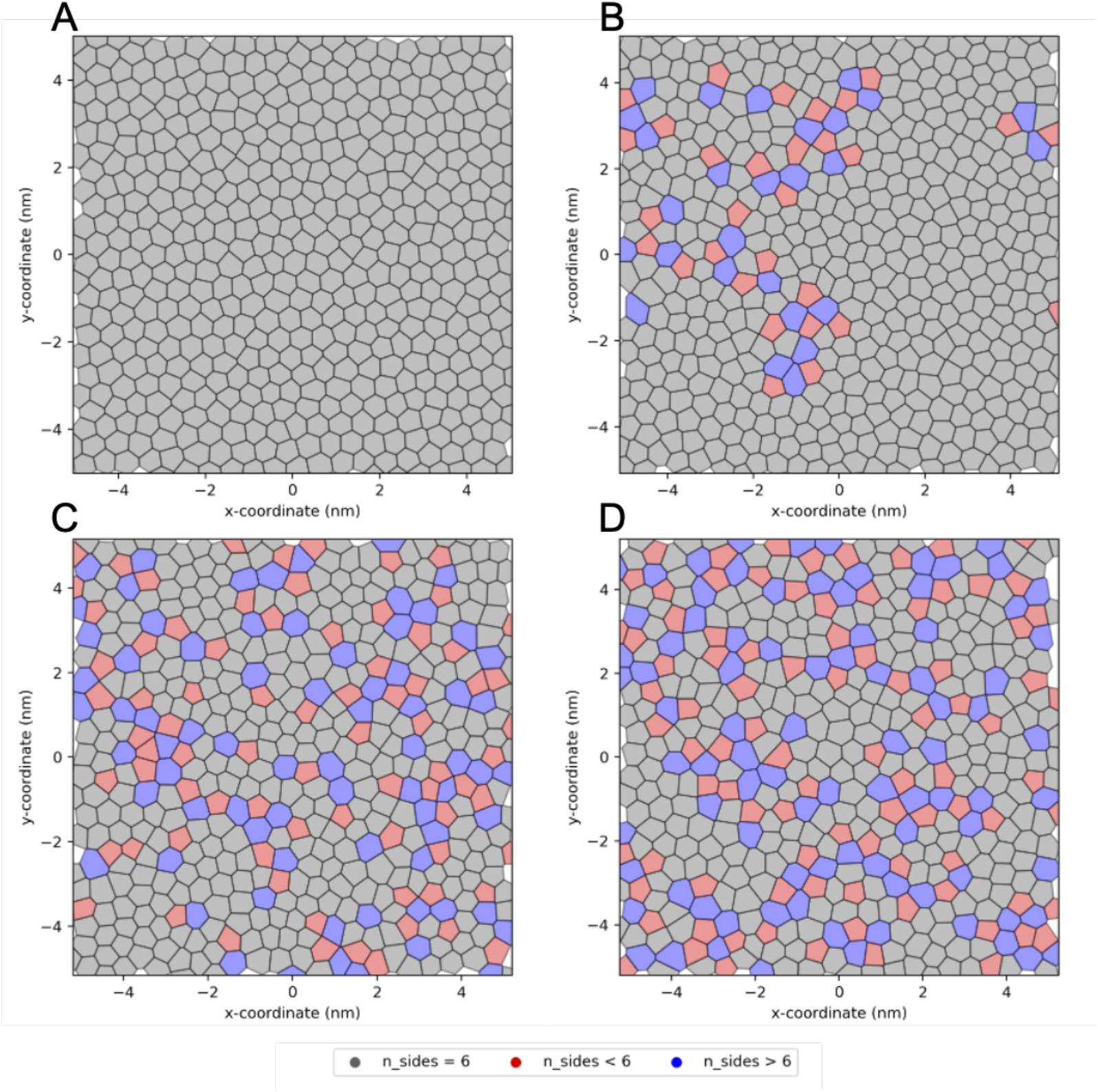
Voronoi diagrams of an interior leaflet of a 6-leaflet CER NS C24/CHOL/FFA C24 system in a) 1:0:1, b) 1:0.2:1, c) 1:0.5:1, and d) 1:1:1 ratios. The polytopes are colored by the number of sides, where those with 6 sides are grey, less than 6 sides are red and greater than 6 sides are blue.

**Figure S7:**
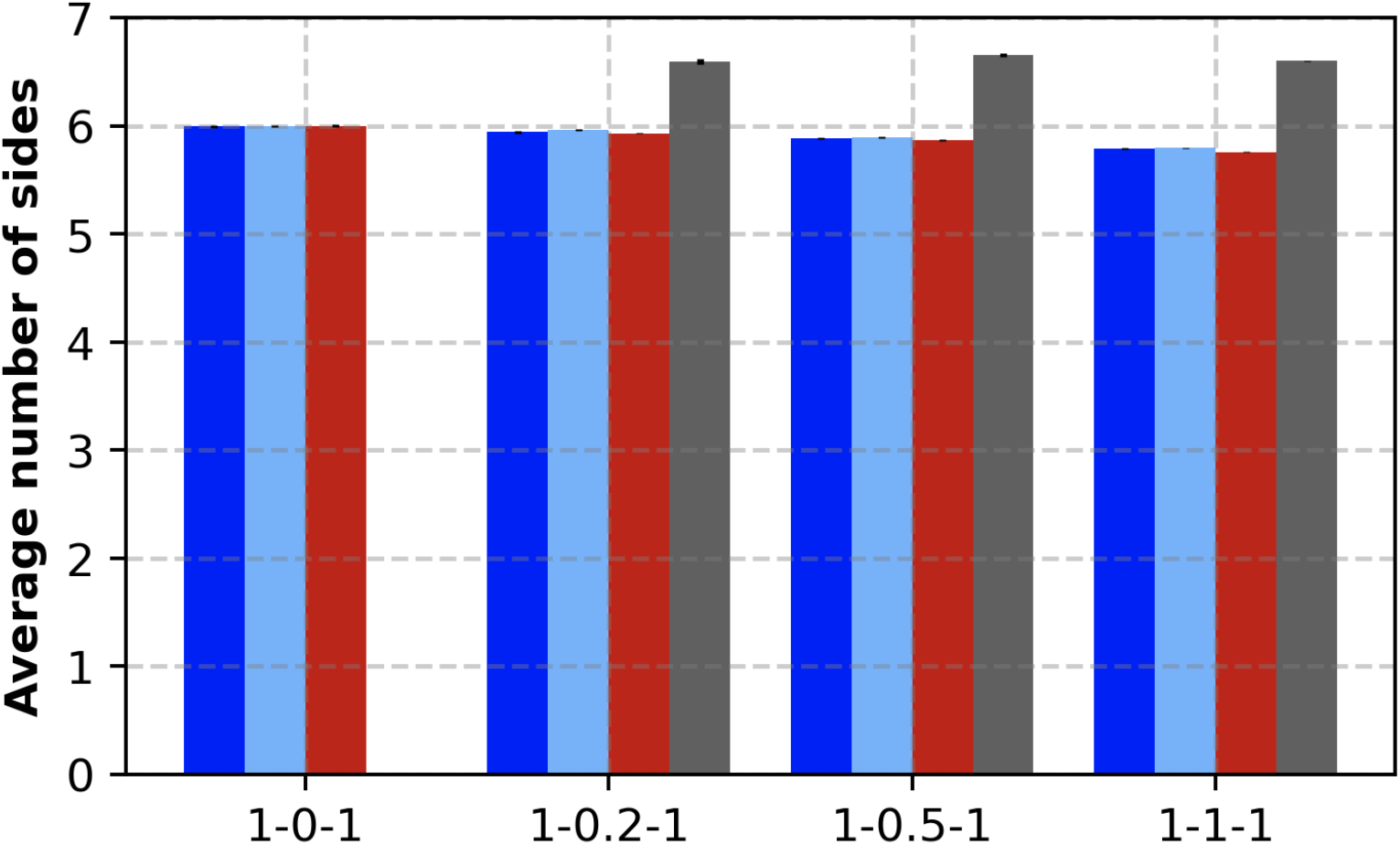
Based on Voronoi diagrams, the average number of sides for each lipid type is plotted above, where dark blue represents the CER NS Acyl chain, light blue represents the CER NS sphingosine chain, red represents FFA, and grey represents CHOL.

## Reverse Mapped Structures

Table S5 compares structural properties of the reverse-mapped atomistic structures to the CG self-assembled structures. Many of the small differences between the CG and reverse-mapped configurations can be explained by the significant presence of lipids stuck in the tail region of the bilayer. For example, the APT is smaller for the reverse-mapped system as the lipids stuck in the tail region are not in-plane with the other lipids, the tilt angle is higher as lipids are sitting parallel to the bilayer plane in the tail region, S_2_ is smaller as the lipids stuck in the tail region are in alignment with the other tails, and the height is larger as the lipids in the tail region push the leaflets apart.

**Table S5:**
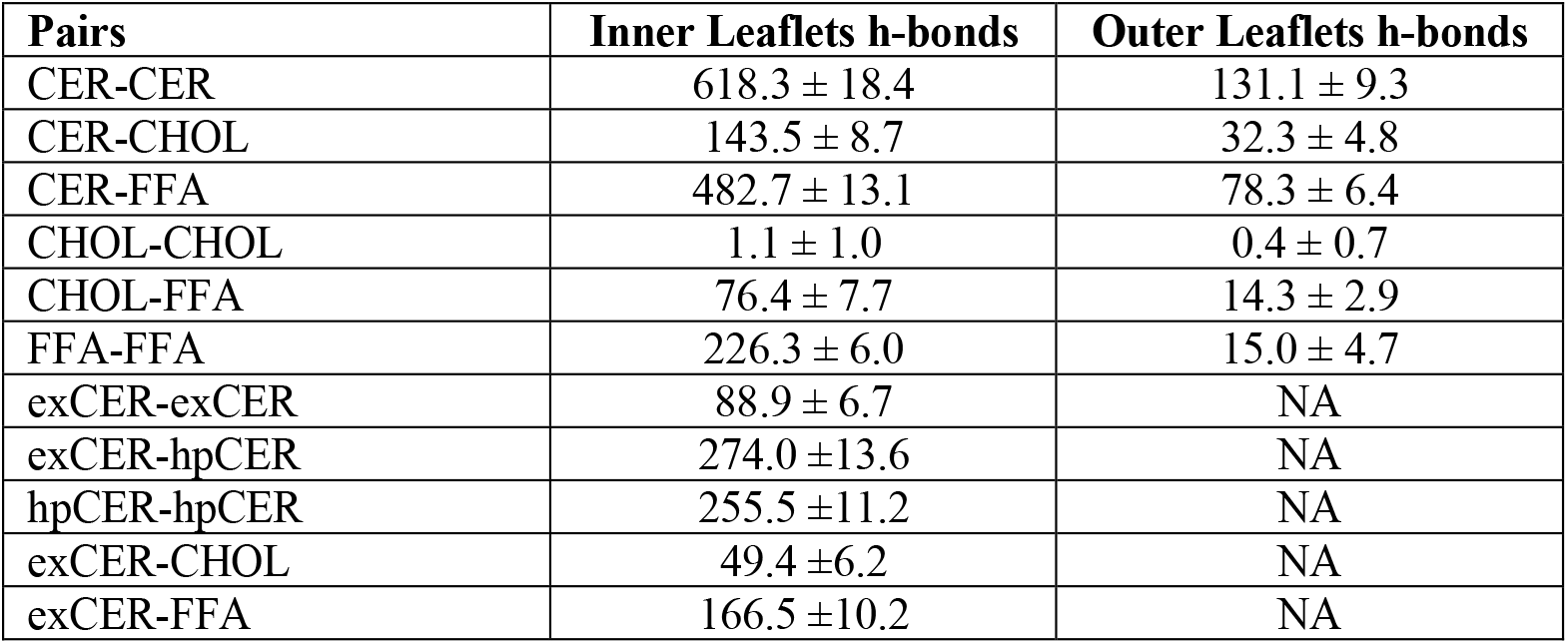
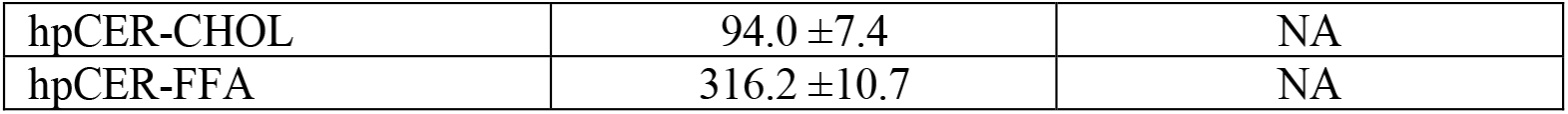
Mean and standard deviation of total number of hydrogen bonds in reverse-mapped atomistic structure. For the inner leaflets there are [538,264, 512] and for outer [262,136,288] [CER, CHOL, FFA24] lipids. For the interior, 187 CERs are I the extended (exCER) and 351 in the hairpin (hpCER) configurations.

| Pairs | Inner Leaflets h-bonds | Outer Leaflets h-bonds |
| --- | --- | --- |
| CER-CER | 618.3 ± 18.4 | 131.1 ± 9.3 |
| CER-CHOL | 143.5 ± 8.7 | 32.3 ± 4.8 |
| CER-FFA | 482.7 ± 13.1 | 78.3 ± 6.4 |
| CHOL-CHOL | 1.1 ± 1.0 | 0.4 ± 0.7 |
| CHOL-FFA | 76.4 ± 7.7 | 14.3 ± 2.9 |
| FFA-FFA | 226.3 ± 6.0 | 15.0 ± 4.7 |
| exCER-exCER | 88.9 ± 6.7 | NA |
| exCER-hpCER | 274.0 ± 13.6 | NA |
| hpCER-hpCER | 255.5 ± 11.2 | NA |
| exCER-CHOL | 49.4 ± 6.2 | NA |
| exCER-FFA | 166.5 ± 10.2 | NA |
| hpCER-CHOL | 94.0 ± 7.4 | NA |
| hpCER-FFA | 316.2 ± 10.7 | NA |

**Table S6:**
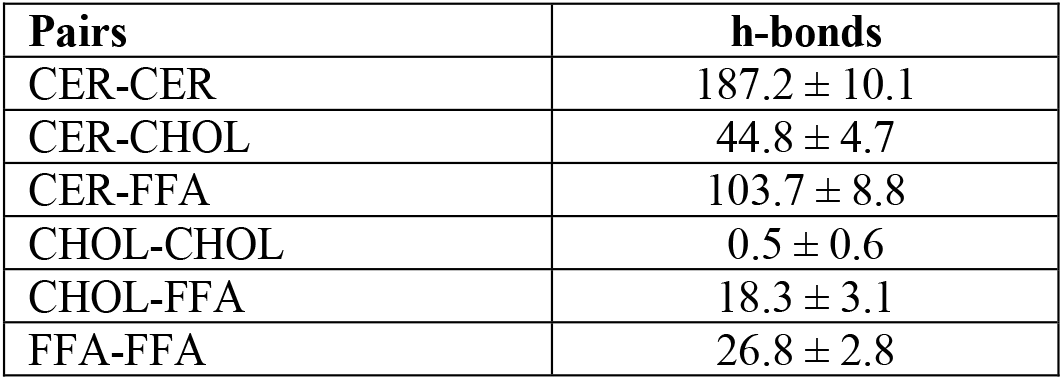
Mean and standard deviation of total number of hydrogen bonds in reverse-mapped atomistic structure in the inner most two layers in order to capture only intra-layer lipid-lipid hydrogen bonding. This inner slice includes 278 CER, 139 CHOL, and 236 FFA molecules. Of the CERs, 104 are in extended conformations and 174 in hairpin.

| Pairs | h-bonds |
| --- | --- |
| CER-CER | 187.2 ± 10.1 |
| CER-CHOL | 44.8 ± 4.7 |
| CER-FFA | 103.7 ± 8.8 |
| CHOL-CHOL | 0.5 ± 0.6 |
| CHOL-FFA | 18.3 ± 3.1 |
| FFA-FFA | 26.8 ± 2.8 |

