## Supplemental Information for "Development of coarse-grained model for a minimal stratum corneum lipid mixture"

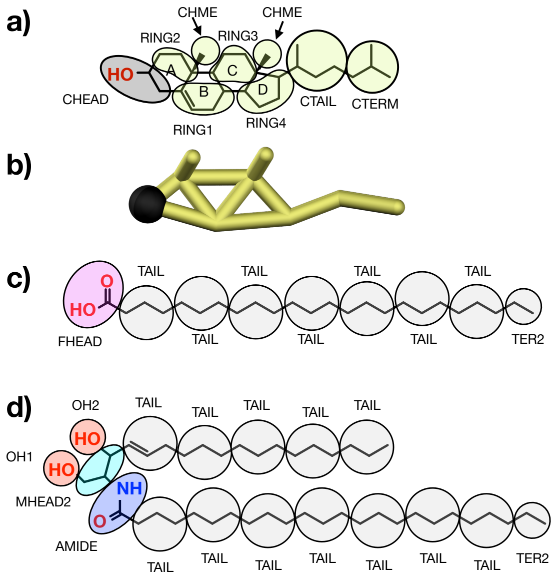


**Figure 1:** *(a) CG mapping scheme for CHOL where the A, B, C, and D rings of the skeletal structure are labeled, and (b) CG representation of CHOL. CG mapping scheme for (c) FFA $C_{24}$, and (d) CER NS $C_{24}$. In all cases, CG beads are spherically symmetric. CG mappings are colored to be consistent with simulation renderings in this work: CHEAD is black, CBODY (i.e., RING1-4, CHME, CTAIL, CTERM) yellow, FHEAD purple, TAIL and TER2 gray, OH1 and OH2 red, MHEAD2 cyan, and AMIDE blue.*

$$V^{i+1}(r) = V^{i}(r) - N_{states}^{-1} \sum_{states} \alpha_s(r) k_B T_s \ln[\frac{g_s^{target}(r)}{g_s^{i}(r)}] \quad (1)$$

where $V^i(r)$ and $V^{i+1}(r)$ are the potential energy functions at the current and next iterations, respectively, for a given pair of atom types; $g_s^{target}(r)$ and $g_s^i(r)$ are the radial distribution functions between a given pair of atom types calculated respectively from the target atomistic system mapped to the CG level and the current iteration; $\alpha_s(r)$ is a weighting factor for state $s$, $N_{states}$ is the total number of states, $T_s$ the temperature of the target state, and $k_B$ the Boltzmann constant. For the case of a single state point, equation 1 reduces to the standard IBI procedure.

The relative weight ($\alpha_s(r)$) of each of the states allows greater emphasis to be placed on matches with targets of greater importance. In addition, $\alpha_s$ acts as a damping factor to stabilize the optimization procedure by preventing large update terms, which may slow down the rate of convergence. Additionally, for MS-IBI, the weighting functions are defined such that they linearly reduce from their maximum value from $\alpha_s\ (0) = \alpha_{max}$ , to $\alpha\ (r_{max}) = 0$ to suppress artifacts in the interaction potential that can arise due to intermediate and long-range correlations in the RDF following the work in reference.[45] The values of $\alpha_{max}$ are chosen via guidance from prior optimization work and refined via guess and check iteration.

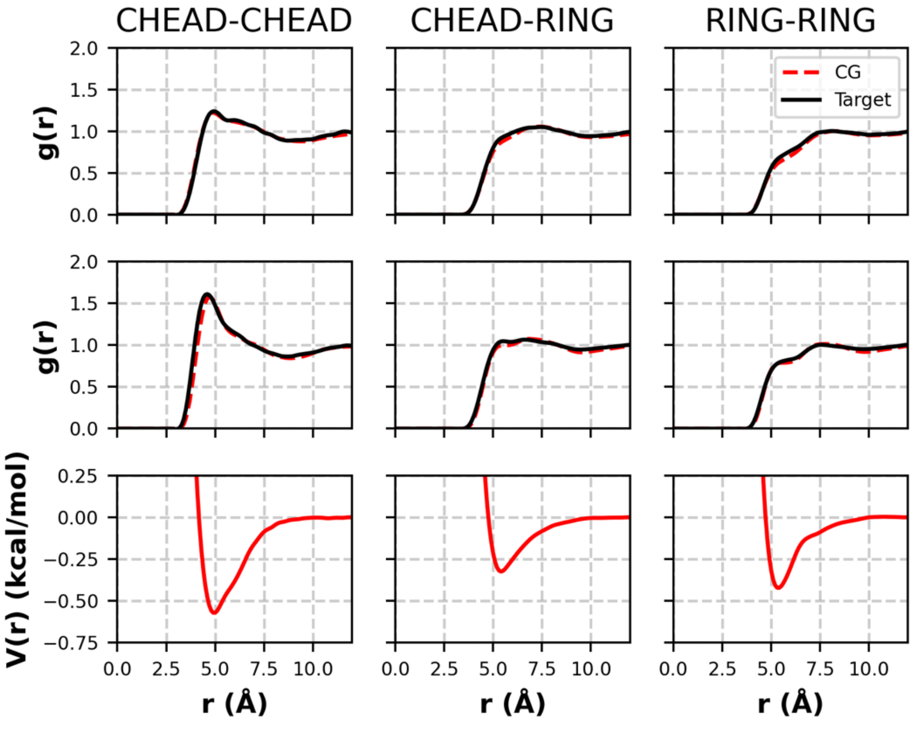


**Figure 2**: *RDFs and pair potentials from the pure CHOL force field optimization. Top: target (black) and CG (red) RDFs at the 550 K NPT state; middle: target (black) and CG (red) RDFs from the 400K NVT state; bottom: pair potential that yields the CG RDFs above.*

**Table 1:** Structural properties of the atomistic and CG systems studied at 305K[a].

| System | APL (Å$^2$) | d (Å) | $S_2$ | Tilt ($^o$) |
|---|---|---|---|---|
| **CER NS $C_{24}$ – CHOL 1:1** | | | | |
| Atomistic Preassembled | 38.8 ± 0.3 | 47.8 ± 0.1 | 0.958 ± 0.007 | 8.7 ± 0.8 |
| CG Preassembled | 39.74 ± 0.06 | 48.0 ± 0.1 | 0.931 ± 0.003 | 10.8 ± 0.2 |
| CG Self-assembled | 38.71 ± 0.16 | 48.33 ± 0.08 | 0.947 ± 0.014 | 9.65 ± 0.13 |
| **CHOL – FFA $C_{16}$ 1:1** | | | | |
| Atomistic Preassembled | 29.92 ± 0.09 | 35.9 ± 0.2 | 0.951 ± 0.004 | 10.8 ± 0.5 |
| CG Preassembled | 30.58 ± 0.08 | 33.22 ± 0.05 | 0.908 ± 0.004 | 13.0 ± 0.3 |
| **CER NS $C_{24}$ – CHOL – FFA $C_{24}$ 1:0.5:1** | | | | |
| Atomistic Preassembled | 30.38 ± 0.05 | 52.7 ± 0.18 | 0.977 ± 0.014 | 7.5 ± 0.3 |
| CG Preassembled | 33.08 ± 0.17 | 50.7 ± 0.2 | 0.96 ± 0.03 | 8.2 ± 0.3 |
| CG Self-Assembled | 33.32 ± 0.16 | 50.82 ± 0.02 | 0.96 ± 0.02 | 8.4 ± 0.2 |

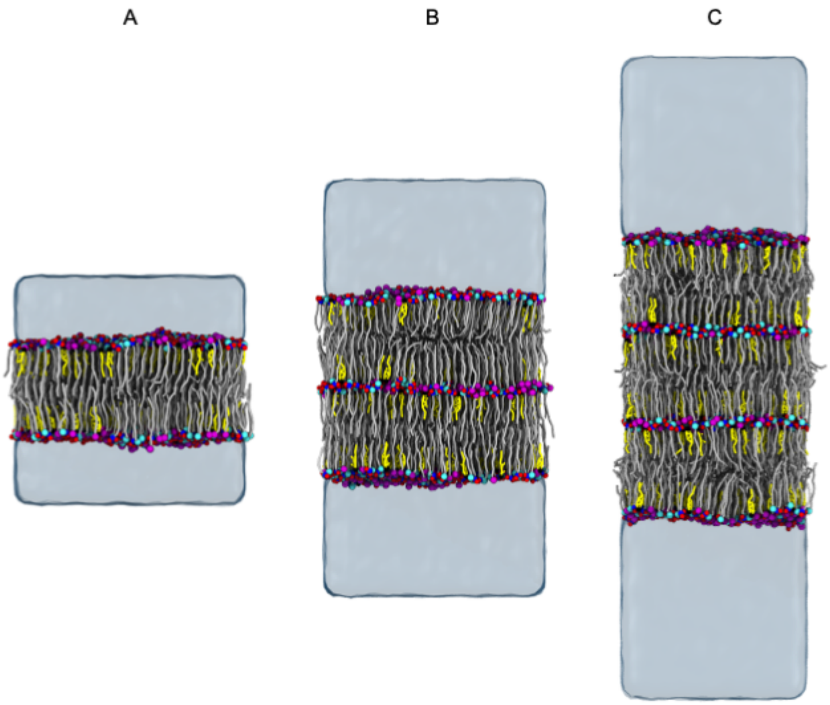


**Figure 3:** Simulation snapshots of self-assembled mixtures of CER NS $C_{24}$/CHOL/FFA $C_{24}$ with a 1:0.5:1 molar ratio in (A) bilayer, (B) 4-leaflet, and (C) 6-leaflet configurations. The lipid molecules are colored as defined in Figure 1. Water is represented as a transparent surface.

The combination of low hydration and the presence of extended CERs in the interior layers of the multilayer systems affects structural properties as shown in Table 2, which reports the average across all leaflets, except for the fraction of extended CERs and the water molecules per lipid, which are calculated from the leaflets without bulk water contact. The decrease in APL for the 4- and 6-leaflet systems compared to the 2-leaflet systems, can be attributed to the increase in extended CERs, which pack more efficiently. At the same time, the tilt angle for the 6-leaflet system is larger than for the 2- or 4-leaflet systems. Figure S6, which shows the distribution of angles between the acyl and sphingosine chains, indicates that the angle between the tails of extended CERs is ~170º rather than the fully linear 180º. This slightly "V"-shaped conformation may arise to optimize chain packing similar to the V-shaped arrangements proposed for CER NP $C_{24}$ mixed with CHOL and FFA $C_{24}$,[62] as well as those observed for pure CER NP crystals. Furthermore, the increased fraction of extended CERs in this slight V-shaped conformation may impart the higher tilt angle observed in the 6-leaflet system. The average bilayer height increased by ~2-3 Å as the number of layers increases, which is consistent with the reduced lipid interdigitation. The bilayer height of 53.81 Å for 6 layers closely agrees with the experimentally measured 54 Å height of the SPP phase, likely due to the similarities in the lipid hydration and the presence of extended CERs.[15,20]

**Table 2:** Average structural properties of 2-, 4-, and 6-leaflet systems of CER NS $C_{24}$/CHOL/FFA $C_{24}$ with a molar ratio 1:0.5:1 [a]

| | 2-leaflet | 4-leaflet | 6-leaflet |
|---|---|---|---|
| **APL (Å2)** | 33.7 ± 0.2 | 33.40 ± 0.13 | 33.14 ± 0.08 |
| **Tilt Angle (º)** | 8 ± 5 | 8 ± 5 | 9 ± 5 |
| **Interdigitation (Å)** | 10.7 ± 0.2 | 10.13 ± 0.14 | 9.2 ± 0.3 |
| **Height (Å)** | 50.82 ± 0.02 | 52.94 ± 0.06 | 53.81 ± 0.10 |
| **% Interior CERs Extended** | -- | 36 | 35 |
| **Water Molecules per Lipid in Interior** | -- | 1.5 | 2.6 |

[a] Results are the average across all leaflets (mean ± one standard deviation for four replicate simulations initialized with different randomized configurations), except for the fraction of extended CERs and the water molecules per lipid, which are calculated from the leaflets without bulk water contact.

*Influence of CHOL content on self-assembled CG systems*

To examine the role of CHOL in the SC, the mole ratio of CHOL in the minimal SPP systems containing equimolar CER NS $C_{24}$ and FFA $C_{24}$ was varied from 0 to 1 relative to CER NS in self-assembly simulations (Fig. 4). As expected, extended CER NS molecules are abundant in the inner bilayers of the 6-leaflet systems and absent in the outer leaflets (Figure 5a). However, the fraction of extended CER NS molecules varies as a function of the CHOL content, with equimolar CHOL:CER NS demonstrating the highest fraction and also the highest area per lipid tail (APT). Perhaps the increased concentration of bulky CHOL tails and consequent increase in APT is countered by an increase in extended CERs, which pack more efficiently.

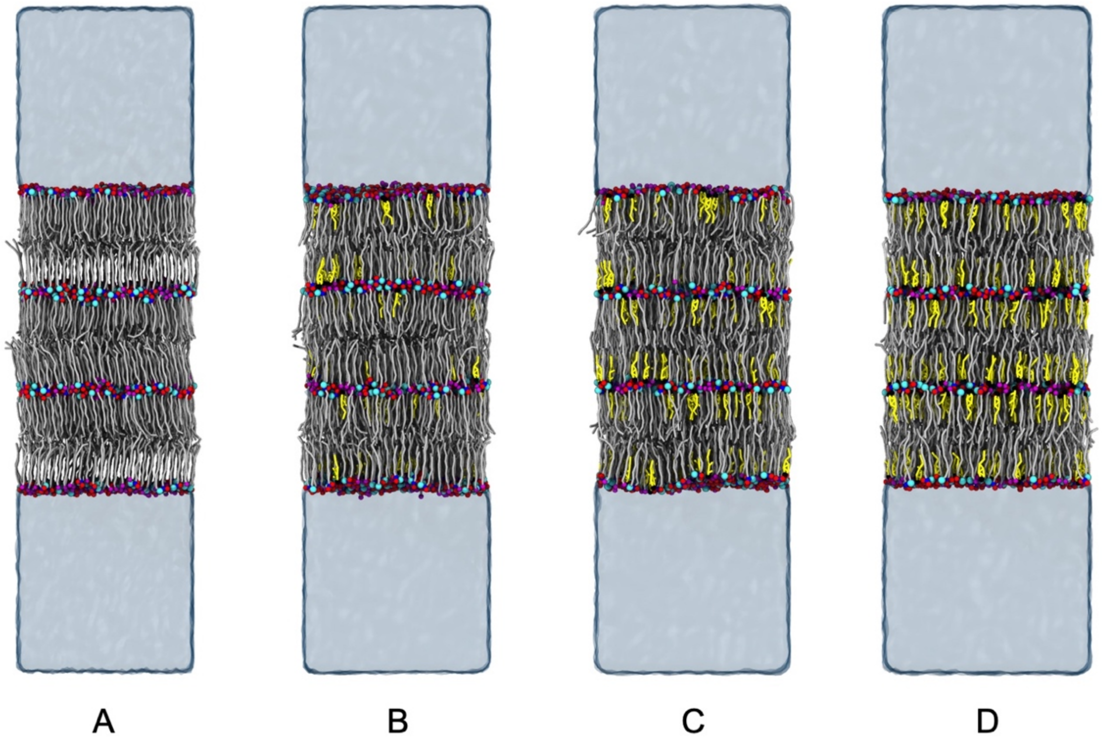


**Figure 4:** *Snapshots from simulations of the 6-leaflet system of CER NS $C_{24}$/CHOL/FFA $C_{24}$ with mole ratios of a) 1:0:1, b) 1:0.2:1, c) 1:0.5:1, and d) 1:1:1. The lipid molecules are colored as defined in Figure 1. Water is represented as a transparent surface.*

Figure 5 shows the trends in the structural properties of the central bilayer in the CER NS $C_{24}$/CHOL/FFA $C_{24}$ systems as a function of CHOL concentration. From the figure, we can see that the area per tail increases as the fraction of CHOL increases. This is consistent with experimental measurements in which hydrocarbon lipid tails typically have a cross-sectional area of 20 $Å^2$ while cholesterol molecules have a cross-sectional area of roughly 38 $Å^2$.[63] There is a significant decrease in the $S_2$ and tilt angle upon addition of CHOL into the bilayer, indicating that the increased CHOL content disrupts the hydrocarbon tail packing. However, further increases in CHOL content did not have a significant effect on the $S_2$ and tilt angle. Also plotted in Figure 5 is a comparison of the same properties for the outer bilayers of each system. The structure of these interfacial leaflets, which would be characteristic of single bilayer systems, have several distinct differences as compared the dehydrated inner region. In particular, the outer bilayers have a smaller tilt angle than the outer bilayers for the 1:0:1 system, which may be attributed to fewer ceramides in the extended conformation, as seen when compared to bilayers in Table 2. A layer-by-layer analysis demonstrates differences in the number of lipids at the water-lipid interface, as compared to the inner leaflets with a lipid-lipid interface. Similarly, the APL and tilt angle differ within these inner leaflets as compared to those in contact with water. As explained earlier, the presence of extended CERs increases the APL and decreases the tilt as observed when comparing multilayer and bilayer systems of CER NS $C_{24}$/CHOL/FFA $C_{24}$ in a 1:0.5:1 ratio. The most significant difference is the ability of the multilayer structures to form both hairpin and extended conformations. As can be seen from the figure, the number of extended conformations within these central layers increases from ~34 to 38% as the cholesterol content is increased. Finally, we note that comparison to experimental repeat distances when available shows close agreement (see Figure 5d) in both trends and quantitative values, though we note that the experimental systems contain a mixture of CERs NS, NP, AP, AS, and 7 FFAs tail lengths ranging from C16-C26.

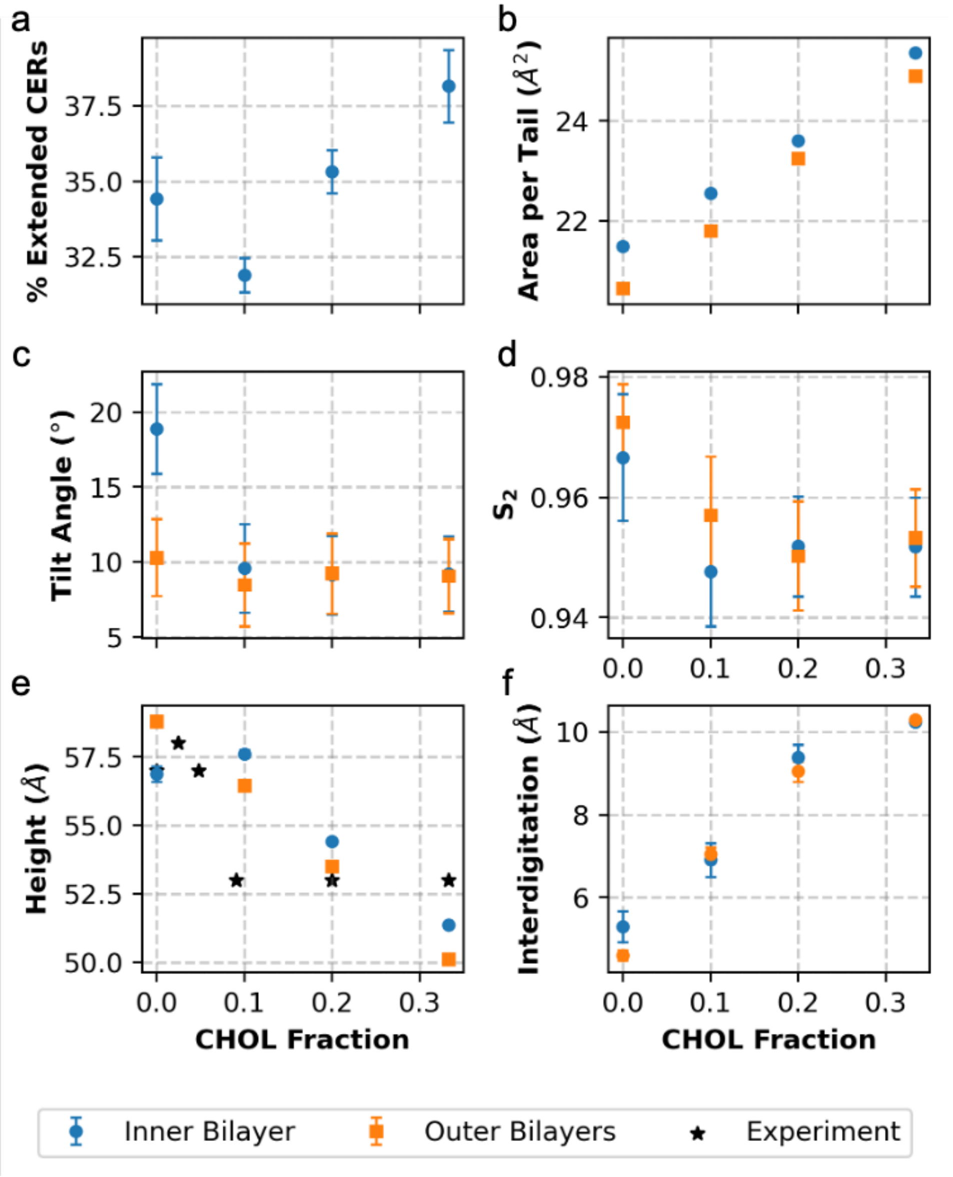


**Figure 5:** *Average structural properties for the inner (blue circles) and outer (orange squares) bilayers of the 6-leaflet system with varying amounts of CHOL in an equimolar mixture of CER NS $C_{24}$:FFA $C_{24}$: a) % CERs extended, b) tilt angle, c) APT, d) $S_2$, e) bilayer height, and f) interdigitation). Results are reported as mean ± one standard deviation from four replicate simulations initialized with different randomized configurations. Experimental height measurements (stars in e) from Mojumdar et al.*[20] *are comprised of a mixture of CER types, although predominantly CER NS $C_{24}$ (60 mol%) CHOL, and FFAs of varying tail length from $C_{16}$ to $C_{26}$.*

In order to examine the preferential arrangement and aggregation of lipid tails in the central bilayer, a cluster analysis was performed to identify the number and size of lipid tail clusters. In Figure 6a we note that the normalized number of CHOL clusters decreases as the fraction of CHOL increases. This indicates that there is a higher fraction of large clusters formed. This is also supported by Figure 6b, which shows that there is an increase in the cluster size as the CHOL content increases. One should also note that there is a sharp increase in the cluster size between the 1:0.5:1 and 1:1:1 systems. This supports the experimental observation that found phase separation of CHOL occurs at CHOL fractions above 0.2 for SPP systems;[20] in our system the

CHOL stays within the membrane, rather than forming a separate phase, but clearly phase separates within the layer. One can also see that there is an increase in the normalized number of clusters and a decrease in the mean cluster size for FFA and CER tails as the fraction of CHOL increases, which suggests that CHOL may improve the miscibility of FFA and CER tails in the membrane.

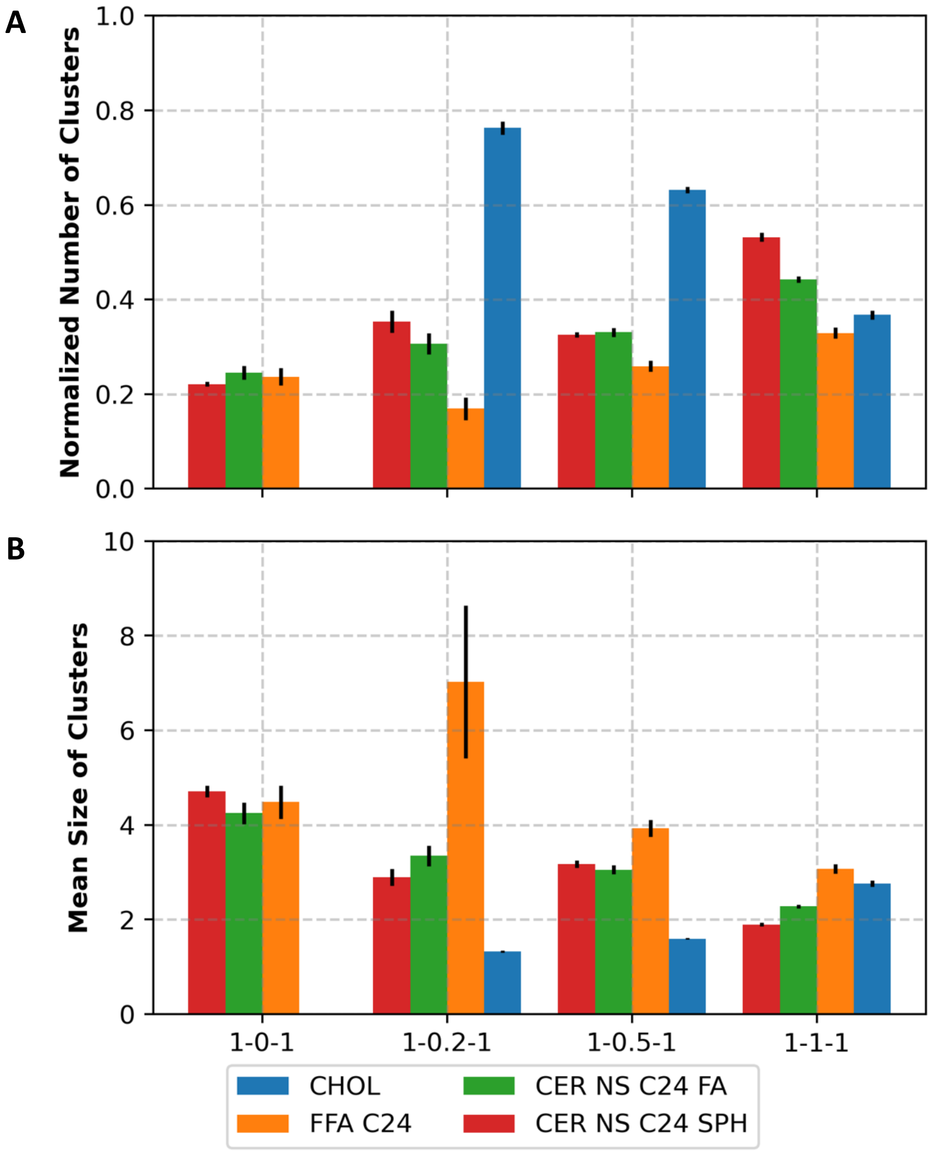


**Figure 6:** *Cluster analysis of lipid tails in the central leaflet. A) the number of clusters normalized by the number of lipids for each lipid type. A value of 1.0 indicates that all clusters contain only a single molecule, meaning that the lipids of that type are fully dispersed. Lower values indicate that more multi-lipid clusters have formed. B) the mean cluster size is shown. All values are averaged over a 2000 frames trajectory spanning 20 ns of equilibrated simulation.*

To further characterize the preferential arrangement of CHOL molecules in these systems, the coordination number of the CHOL tails with each of the lipid tail types was calculated. One should note due to the sparsity of CHOL-CHOL pairs, the coordination number cannot be precisely calculated as the first peak of the CHOL-CHOL RDF is very short. From the results shown in

Figure 7, we can see that between the sphingosine and acyl CER tails, CHOL prefers to neighbor the similar-length sphingosine tails. This is corroborated by experimental NMR spectra, which suggests that the sphingosine chain is more likely to neighbor CHOL molecules in a mixture of CER NS C24, CHOL, and FFA C24 in a 1:0.45:1 ratio.[19] However, these experiments were not able to quantify the extent of this preference and assumed in their proposed model that all of the sphingoid chains of CER neighbor CHOL molecules.

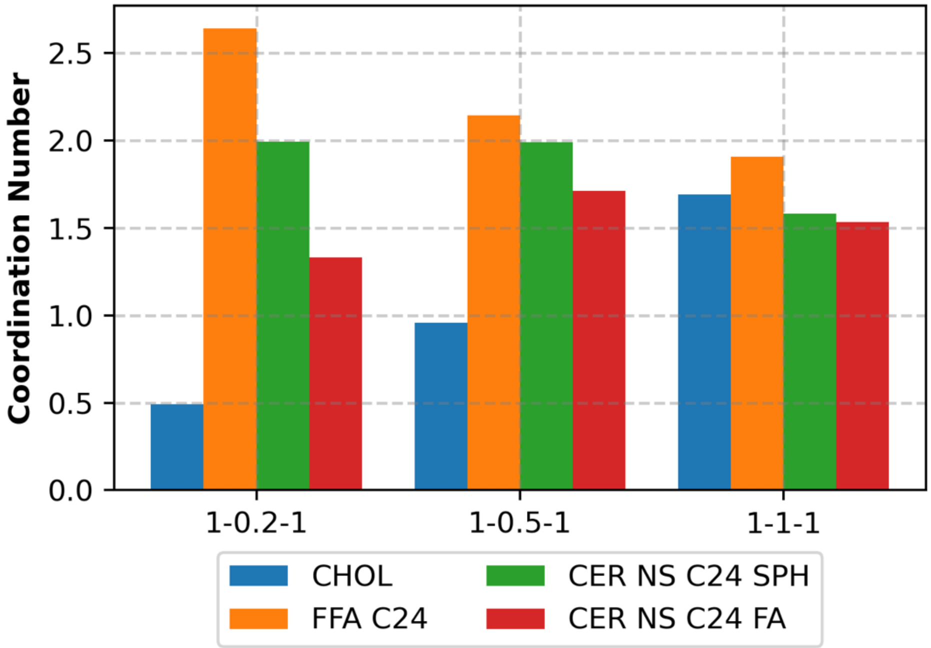


**Figure 7:** *CHOL-X coordination numbers, where X is either CHOL, FFA $C_{24}$, the sphingosine chain of CER NS $C_{24}$ (SPH), or the fatty acid chain of CER NS $C_{24}$ FA.*

From the data shown in Figure 7, we can see that although there is a slight preference for CHOL to neighbor sphingoid chains, CHOL still has a significant number of CER acyl chain neighbors. It is also interesting to note that, contrary to the model hypothesized by Engberg et al., CHOL prefers to neighbor FFA molecules.[19] This may be because FFA molecules have higher mobility compared to CERs due to their single-tailed nature, which may more effectively accommodate packing defects induced by the bulky CHOL tails. This is further supported by the Voronoi analysis provided in the SI, which indicate that 5-neighbored FFAs mitigate hexagonal packing defects introduced by 7-neighbor CHOL molecules.

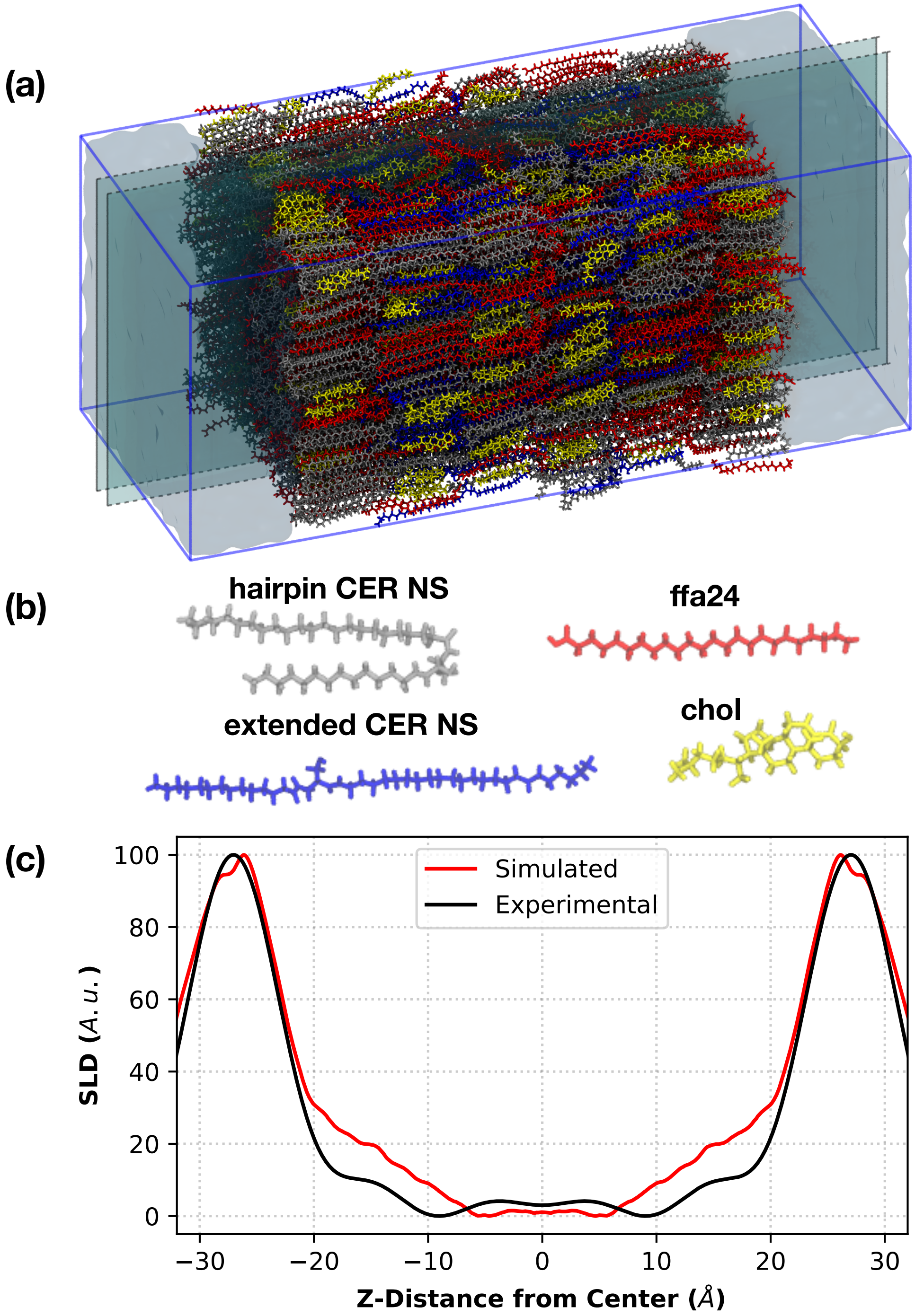


**Figure 8.** (a) *Simulation snapshot of a reverse-mapped atomistic 1:0.5:1 CER NS $C_{24}$/CHOL/FFA $C_{24}$ multilayer, with lipids colored according to (b). A 1 nm thick slice is highlighted in (a) and shown in detail in Figure 9a. (c) Protiated NSLD profiles for the same reverse-mapped 6-leaflet system from the atomistic simulation and experiment reproduced from Groen et al.*[15]

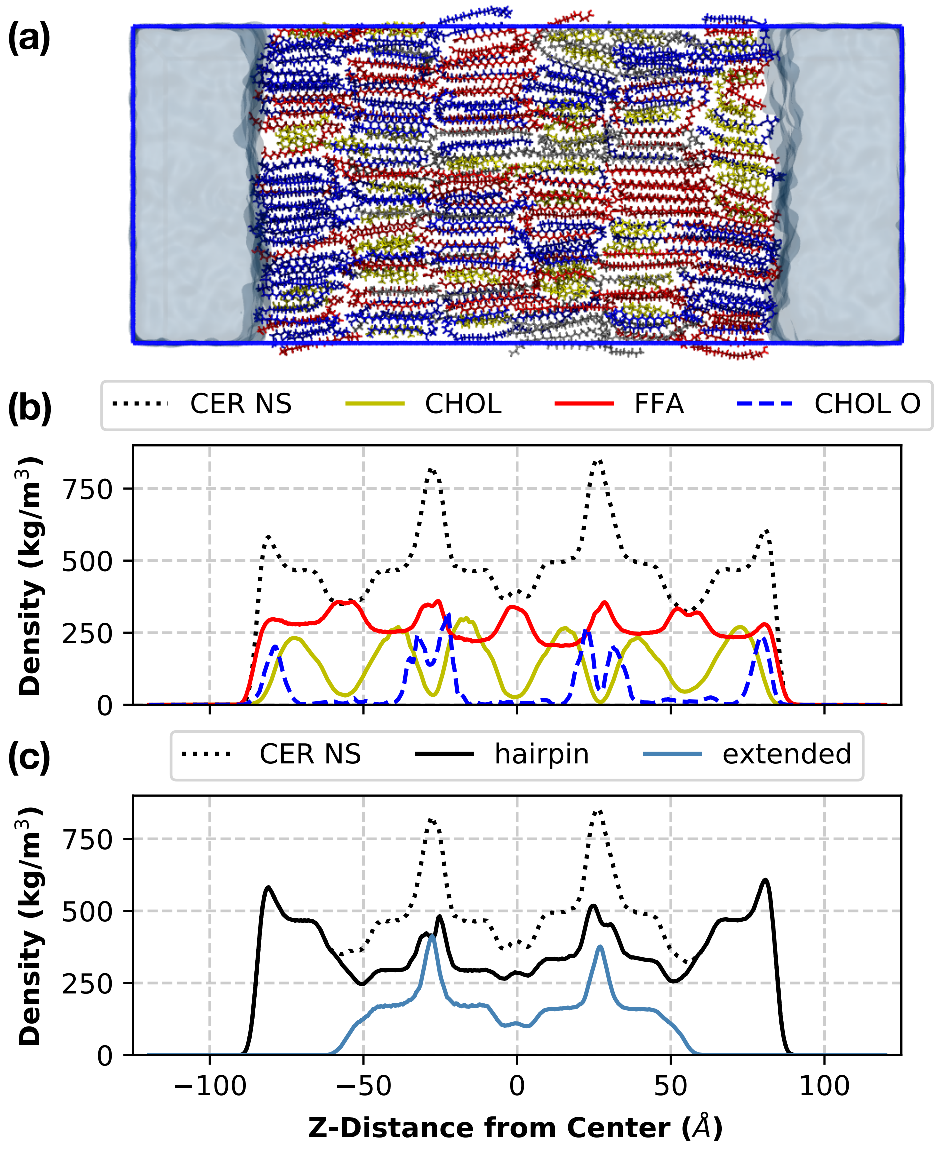


**Figure 9:** *(a) Slice from the center of the simulation cell for the 1:0.5:1 CER NS $C_{24}$/CHOL/FFA $C_{24}$ multilayer shown in Fig.8a, again colored following Figure 8b. (b) density profile of CER NS, CHOL, and FFA species. The density profile of the oxygen in the CHOL headgroup is also plotted, with the numerical values scaled by a factor of 10 in order to make them visible on the plot. (c) Density histograms for CER NS, plotting hairpin and extended CERs separately.*

**Table 3**: *Normalized Number of Lipid-Lipid Hydrogen Bonds by Pairs of Molecule Types*[a]

| Pairs | Inner Leaflets | Outer Leaflets | Normalization |
|---|---|---|---|
| CER-CER | 1.149 ± 0.034 | 0.500 ± 0.036 | # CERs |
| CER-CHOL | 0.267 ± 0.016 | 0.123 + 0.018 | # CERs |
| CER-FFA | 0.897 ± 0.024 | 0.299 + 0.024 | # CERs |
| CHOL-CHOL | 0.004 ± 0.004 | 0.003 ± 0.005 | # CHOLs |
| CHOL-FFA | 0.289 ± 0.029 | 0.105 ± 0.021 | # CHOLs |
| FFA-FFA | 0.442 ± 0.011 | 0.052 + 0.016 | # FFAs |
| FFA-CHOL | 0.149 ± 0.015 | 0.050 ± 0.010 | # FFAs |
| ALL-ALL | 1.178 ± 0.020 | 0.396 ± 0.020 | Total # of lipids |

[a] The values reported are the mean and standard deviation, in each case normalized by the type of lipid listed in the table in either the inner or outer leaflets; unnormalized counts of hydrogen bonds are provided in the SI.

**Table 4**: *Fraction of Lipid-Lipid Hydrogen Bonds by Pairs of Molecule Types comparing hairpin and extended CERs in the Inner Leaflets*[a]

| Pairs | Fraction |
|---|---|

| | |
|---|---|
| exCER-exCER | 0.144 ± 0.082 |
| exCER-hpCER | 0.443 ± 0.058 |
| hpCER-hpCER | 0.413 ± 0.053 |
| exCER-CHOL | 0.345 ± 0.139 |
| hpCER-CHOL | 0.655 ± 0.099 |
| exCER-FFA | 0.345 ± 0.067 |
| hpCER-FFA | 0.655 ± 0.043 |

**Supporting Information for**:

### Development of coarse-grained model for a minimal stratum corneum lipid mixture

Parashara Shamaprasad,[1] Timothy C. Moore,[1] Donna Xia,[1] Christopher R. Iacovella,[1] Annette L. Bunge,[2] and Clare M<sup>c</sup>Cabe[1,3]

[1]Department of Chemical and Biomolecular Engineering, Vanderbilt University, Nashville, TN, 37235-1604
[2]Department of Chemical and Biological Engineering, Colorado School of Mines, Golden, CO, 80401
[3]Department of Chemistry, Vanderbilt University, Nashville, TN, 37235-1604

**Table S1:** Target states used in the MS IBI optimization of CHOL self- and cross-interactions

| Description | State conditions | Temperature, K | Label |
|---|---|---|---|
| **Pure CHOL** | | | |
| Bulk fluid | 1.0 atm (NPT) | 550 | CHOL1 |
| Bulk fluid | 0.85 g/mL (NVT) | 400 | CHOL2 |
| Bulk fluid | 0.74 g/mL (NVT) | 500 | CHOL3 |
| Bulk fluid | 0.65 g/mL (NVT) | 700 | CHOL4 |
| **CHOL-water** | | | |
| 1.0 mol % CHOL | 1.0 atm (NPT) | 305 | W-CHOL1 |
| 2.0 mol % CHOL | 1.0 atm (NPT) | 305 | W-CHOL2 |
| **CER NS-CHOL** | | | |
| Bulk fluid | 1.0 atm (NPT) | 500 | CRCHL-NPT |
| Bulk fluid | 0.74 g/mL (NVT) | 500 | CRCHL-NVT |
| Bilayer | 1 atm (NPT) | 305 | CRCHL-BIL-NPT |
| Bilayer | 38.8 $Å^2$ (NVT) | 305 | CRCHL-BIL-NVT |
| Dehydrated multilayer | 1 atm (NPT) | 305 | CRCHL-MULT-NPT |
| Dehydrated multilayer | 38.8 $Å^2$ (NVT) | 305 | CRCHL-MULT-NVT |
| **CHOL-FFA C24:0** | | | |
| Bulk fluid | 1 atm (NPT) | 400 | CHLFFA-NPT |
| Bulk fluid | 0.79 g/mL (NVT) | 400 | CHLFFA-NVT |
| **CHOL-FFA C16:0** | | | |
| Monolayer | 33.1 $Å^2$ (NVT) | 305 | CHLFFA-MONO |
| Bilayer | 1 atm (NPT) | 305 | CHLFFA-BIL-NPT |
| Bilayer | 29.9 $Å^2$ (NVT) | 305 | CHLFFA-BIL-NVT |
| Dehydrated multilayer | 1 atm (NPT) | 305 | CHLFFA-MULT-NPT |
| Dehydrated multilayer | 29.9 $Å^2$ (NVT) | 305 | CHLFFA-MULT-NVT |

**Table S2:** States used to optimize each pair potential and the weight $\alpha_{max}$ given to each state. Pairs are denoted by bold text, with the states used to optimize a given (set of) interaction(s) listed below the interaction name. The state condition names correspond to the label given in Table S1.

| States used | $\alpha_{max}$ |
|---|---|
| **CHOL–CHOL** | |
| CHOL1 + CHOL2 + CHOL3 + CHOL4 | 0.7 |
| **CHOL–WATER** | |
| W-CHOL1 + W-CHOL2 | 0.4 |
| **CERHEADS–CHEAD** | |
| CRCHL-(NPT + NVT) | 0.05 |
| CRCHL-BIL-(NPT+NVT) | 0.5 |
| CRCHL-MULT-(NPT+NVT) | 0.5 |
| **CERHEADS–CBODY** | |

| | |
|---|---|
| CRCHL-(NPT + NVT) | 0.05 |
| **FHEAD–CHEAD** | |
| CHLFFA-(NPT+NVT) | 0.05 |
| CHLFFA-BIL-(NPT+NVT) | 0.5 |
| CHLFFA-MULT-(NPT+NVT) | 0.5 |
| **FHEAD–CBODY** | |
| CHLFFA-(NPT+NVT) | 0.05 |
| **TAILs–CHEAD** | |
| CHLFFA-(NPT+NVT) | 0.05 |
| **TAILs–CBODY** | |
| CHLFFA-(NPT+NVT) | 0.05 |
| CHLFFA-MONO | 0.5 |
| CHLFFA-BIL-(NPT+NVT) | 0.5 |
| CHLFFA-MULT-(NPT+NVT) | 0.5 |

### Coarse-Grained Simulation Methods

All CG simulations are performed using the GPU-accelerated HOOMD-Blue simulation engine with a 10 fs timestep.[52] Systems using the constant-temperature (NVT) ensemble use a Nosé-Hoover thermostat with a coupling constant of 1 ps,[54,70] and constant-pressure (NPT) CG simulations use the Martyna-Tobias-Klein barostat-thermostat.[54,71,72] Three distinct sets of CG simulations are performed for force field optimization, force field validation, and self-assembly.

**Force Field Optimization and Validation**

The first set is run during the MS-IBI optimizations to generate the RDF for each iteration. These use the final configuration from the corresponding atomistic simulation for the initial configuration, except configurations are replicated twice in each direction for bulk systems and three times in each lamellar lateral direction for the lamellar systems. These simulations use the force field derived in Moore et al. 2018,[30] except for interactions involving CHOL beads, which are optimized during this stage. Topological constraints, i.e., bonds/angles, are determined following the procedure in Milano, et al.[73] Briefly, distributions of the bond distances and/or angles are collected from atomistic simulations mapped to the CG level, with harmonic bond/angle parameters fit to best reproduce these distributions; 11 bond and 23 angle types are present in the CG model, and each have unique stiffnesses and equilibrium separations, where the parameters used for each bond/angle type are provided in the Table S3 of the SI.

**Table S3:** Bonded interaction parameters for CHOL.

| **Name** | **$r_0$ [Å]** | **$K_r$ [kcal/(mol•Å$^2$ )]** |
|---|---|---|
| CHEAD-RING1 | 4.27 | 125.5 |
| CHEAD-RING2 | 2.58 | 245.9 |
| RING1-RING2 | 3.57 | 324.1 |
| RING2-RING3 | 4.2 | 132.2 |
| RING3-RING4 | 3.21 | 362.6 |
| RING1-RING4 | 4.48 | 152.1 |
| RING1-RING3 | 3.6 | 352.3 |

| | | |
|---|---|---|
| RING4-CTAIL | 3.69 | 50.3 |
| CTAIL-CTERM | 4.22 | 13.2 |
| RING2-CHME | 2.37 | 65.4 |
| RING3-CHME | 2.66 | 116.2 |
| **Angle Name** | **$\theta_0$ [º]** | **$K_0$ [kcal/ (mol•rad$^2$)]** |
| RING1-CHEAD-RING2 | 56.3 | 1132.6 |
| RING2-RING1-RING4 | 115.1 | 663.1 |
| RING3-RING1-RING4 | 44.8 | 3617.7 |
| RING2-RING1-RING3 | 71 | 1323 |
| CHEAD-RING1-RING4 | 151.2 | 351.3 |
| CHEAD-RING1-RING3 | 144.3 | 180.2 |
| CHEAD-RING1-RING2 | 36.7 | 2941.4 |
| RING1-RING2-RING3 | 54 | 2678.8 |
| CHEAD-RING2-RING3 | 139.4 | 597.5 |
| CHEAD-RING2-RING1 | 85.4 | 1011.1 |
| CHEAD-RING2-CHME | 92.8 | 75.5 |
| RING3-RING2-CHME | 60.9 | 436.6 |
| RING1-RING2-CHME | 49.7 | 643.4 |
| RING2-RING3-CHME | 133.2 | 77.5 |
| RING1-RING3-RING4 | 81.1 | 1329.5 |
| RING1-RING3-RING2 | 53.4 | 2787.1 |
| RING4-RING3-CHME | 53.6 | 671.7 |
| RING2-RING3-RING4 | 133.4 | 446.5 |
| RING1-RING3-CHME | 95.5 | 354.5 |
| RING1-RING4-RING3 | 52.5 | 2675.5 |
| RING3-RING4-CTAIL | 94.4 | 276.1 |
| RING1-RING4-CTAIL | 144.3 | 180.2 |
| RING4-CTAIL-CTERM | 125 | 7.6 |

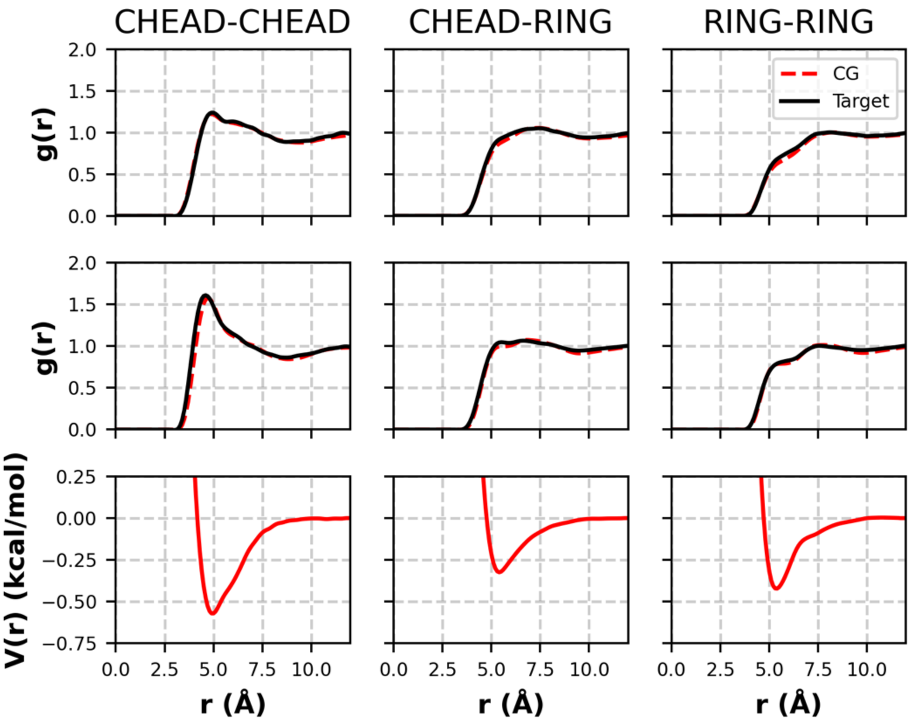


**Figure S1:** *RDFs and pair potential from the pure CHOL force field optimization. Top: target(black) and CG (ref) RDFs from the 400K NVT state; middle: target and CG RDFs at the 550 K NPT state; bottom: pair potential that yields the CG RDFs above.*

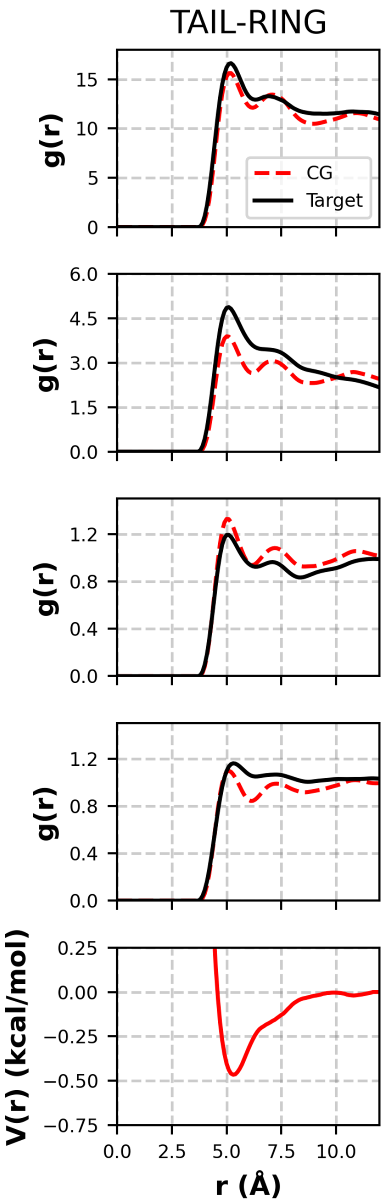


**Figure S2:** *TAIL–RING RDFs and pair potential. From top to bottom: RDFs from the monolayer state; RDFs at the NPT bilayer state; RDFs at the stacked bilayer state; RDFs at the bulk fluid NPT state; and the pair potential that yields the CG RDFs above.*

The CHEAD–FHEAD RDFs are decently captured at the lamellar states, as shown in Figure S3. Interestingly, the best fit is found at the stacked bilayer state, even compared to the bilayer state. This is likely a result of the relative locations of the first peaks: at the bilayer state, the first peak in the RDF is at a larger separation compared to the stacked bilayer and bulk states. To compensate, the pair potential shifts the first RDF peak to smaller separation values. Like in previous optimizations, the fits at the lamellar states are better than at the bulk states as a result of the lamellar states being given a larger weight in the optimization process. The RDFs between the mixed hydrophilic–hydrophobic beads, e.g., TAIL–CHEAD (shown in Figure S3), are captured with less accuracy than in previous Moore et al. (2016).[38] This may be a result of the shape of CHOL, which is more complex than the simpler chain-like CER and FFA molecules. Despite the discrepancies, the density of the bulk fluid NPT state was found to be 0.81 ± 0.03 g/mL, which is in good agreement with the atomistic value of 0.79 ± 0.01 g/mL. Additionally, in a lamellar configuration, there are likely to be very few of these types of interactions. Therefore, these deviations in the RDF are deemed acceptable considering the target objective of this force field.

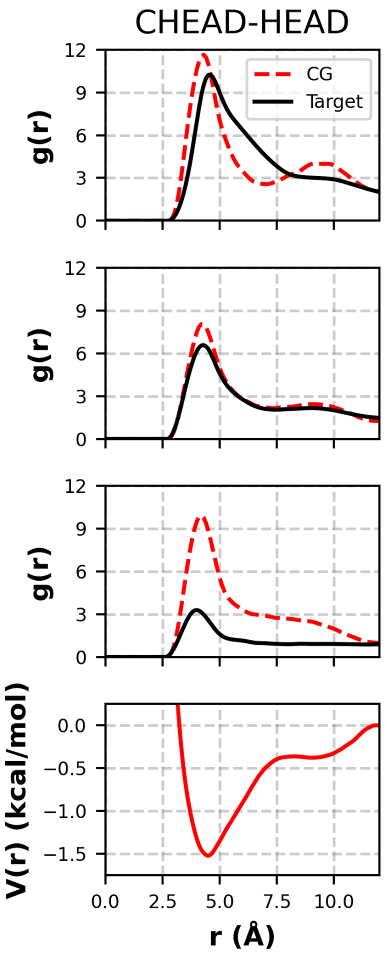


**Figure S3:** *CHEAD–FHEAD RDFs and pair potential. From top to bottom: RDFs from the bilayer state; RDFs from the stacked bilayer state; RDFs from the bulk fluid NPT state; and the pair potential that yields the CG RDFs above.*

**CER NS–CHOL**

Only the CERHEADs–CHEAD and CERHEADs–CBODY interactions are considered here, since the TAILs–CHOL interactions were optimized using target data from CHOL–FFA systems. In general, as shown in Figure S4, the CERHEADs–CHEAD interactions are accurately reproduced at the lamellar states.

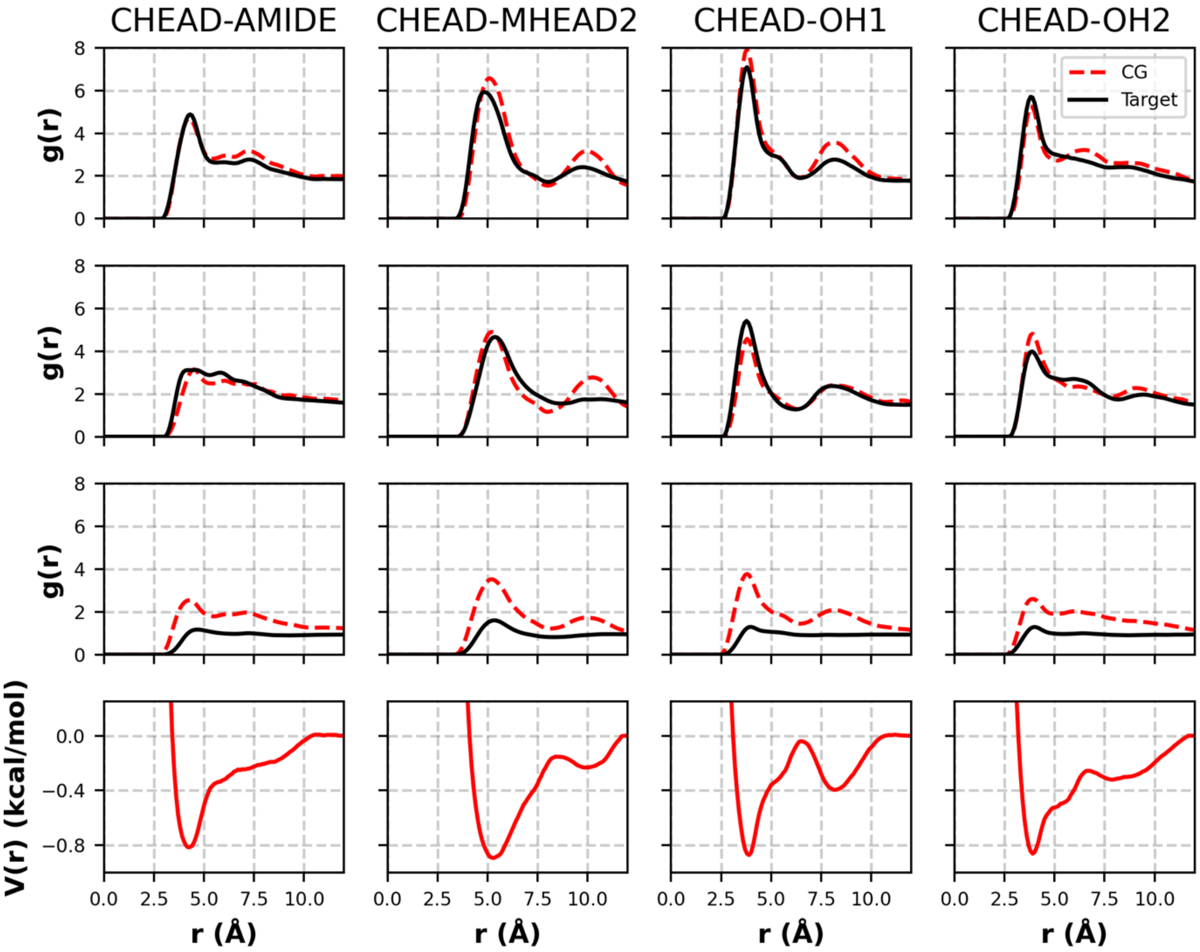


**Figure S4:** *CERHEADS–CHEAD RDFs and pair potential. From top to bottom: RDFs at the stacked bilayer state; RDFs at the bilayer state; RDFs at the bulk fluid NPT state; and the pair potentials that yield the RDFs above. Target RDFs are shown by solid black lines, and CG RDFs are shown as the dashed red lines. Each column corresponds to a given pair, listed at the top of that column.*

**Table S4:** Molar ratios of systems used for validation.

| CER:CHOL:FFA C24 | CER NS C16:CER NS C24 | Total CER NS C16:CER NS C24:CHOL:FFA C24 |
|---|---|---|
| 1:1:0 | 4:0 | 4:0:4:0 |
| 1:1:0 | 3:1 | 3:1:4:0 |
| 1:1:0 | 2:2 | 2:2:4:0 |
| 1:1:0 | 1:3 | 1:3:4:0 |
| 1:1:0 | 0:4 | 0:4:4:0 |
| 2:1:0 | 4:0 | 4:0:2:0 |
| 2:1:0 | 3:1 | 3:1:2:0 |
| 2:1:0 | 2:2 | 2:2:2:0 |
| 2:1:0 | 1:3 | 1:3:2:0 |
| 2:1:0 | 0:4 | 0:4:2:0 |
| 1:1:1 | 4:0 | 4:0:4:4 |
| 1:1:1 | 3:1 | 3:1:4:4 |
| 1:1:1 | 2:2 | 2:2:4:4 |
| 1:1:1 | 1:3 | 1:3:4:4 |
| 1:1:1 | 0:4 | 0:4:4:4 |

Figure S4 compares structural properties of the CG self-assembled systems to the atomistic simulation results obtained by Moore et al.[22] Prior to analysis, the atomistic trajectories are mapped to the CG resolution using the method described in the Analysis Methods section of the SI. The values in table S4 for CG systems are averaged from four replicate systems initialized from different randomized configurations. From these results, we can see close agreement between the CG and atomistic values. One should note that several CHOL molecules were "stuck" in the center of the atomistic bilayers simulated by Moore et al.[22] This resulted in a slightly lower area per tail, due to the decrease in bulky CHOL molecules at the water interface, and increased the height, as the CHOLs in the center of the bilayer act as spacers between the two leaflets.

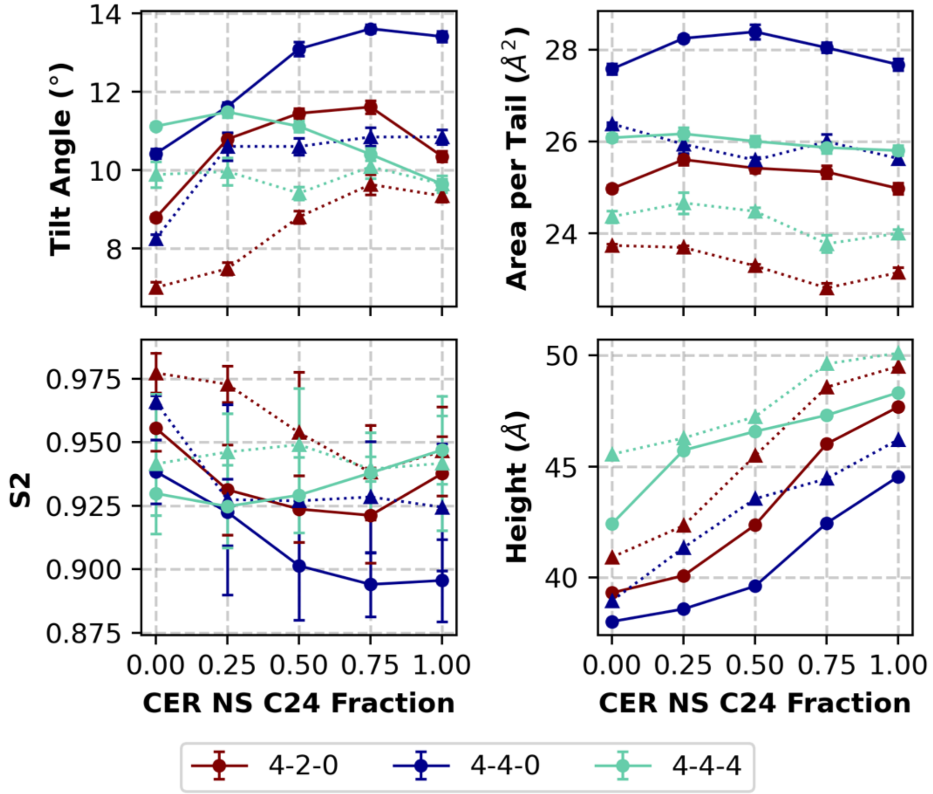


**Figure S4:** Structural parameters of CER NS:CHOL:FFA C24 systems with 4:2:0, 4:4:0 and 4:4:4 ratios, for which the ratio of CER NS C16:CER NS C24 is varied from pure CER NS C16 to pure CER NS C24 in five evenly spaced increments (i.e. 4:0, 3:1, 2:2, 1:3, and 0:4). Results from self-assembled CG bilayers are represented by circles and connected by a solid line, whereas results from preassembled atomistic bilayers (taken from Moore et al.) are represented by triangles and connected by a dashed line.

### Tail Angle Distributions for CER NS C24/CHOL/FFA C24 Multilayers with Varying CHOL content

A histogram of the angles between the sphingosine and acyl chains of interior CERs are plotted in figure S5.

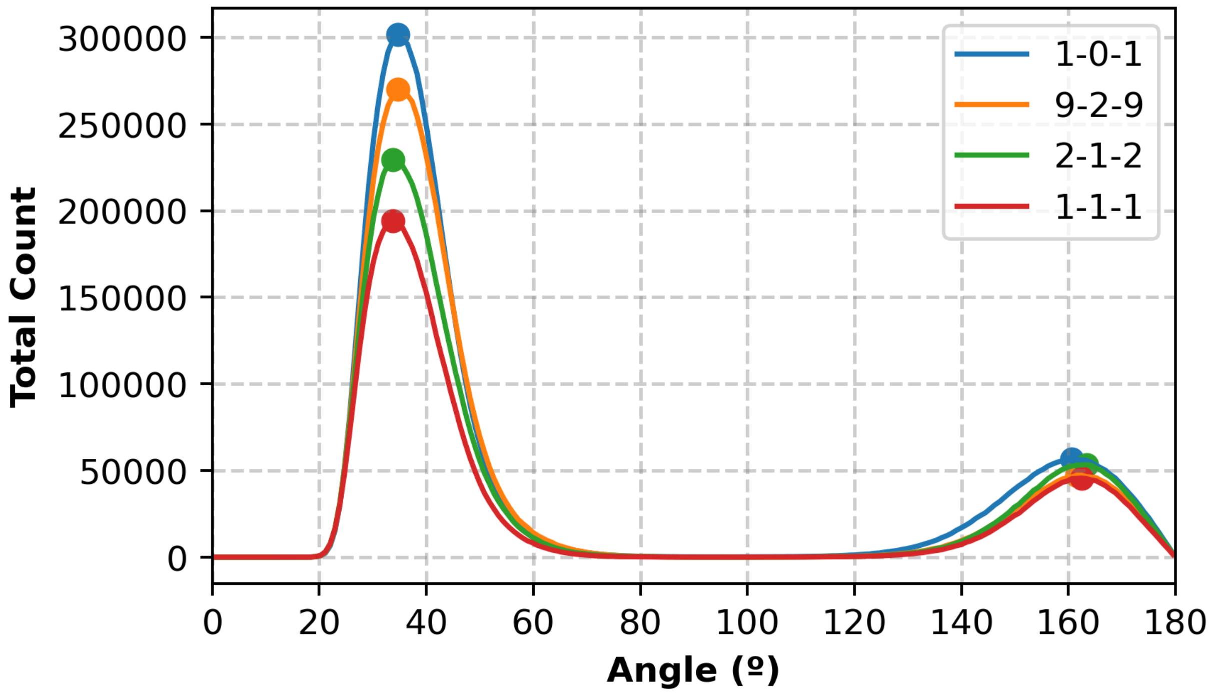


**Figure S6:** *Histograms of the angle between the acyl and sphingosine chains of CER NS molecules in the interior of CER NS C24/CHOL/FFA C24 6-leaflet multilayers in 1:0:1, 1:0.2:1, 1:0.5:1, and 1:1:1 ratios.*

#### Voronoi Analysis of CER NS C24/CHOL/FFA C24 Multilayers with Varying CHOL Content

Voronoi analysis of the interior leaflets of the CER NS C24/CHOL/FFA C24 multilayers was conducted using Freud as described in the "Analysis Methods" section of the SI. Voronoi diagrams of the last frame of first trial for each lipid ratio is shown in figures S6. Each polygon is colored by the number of sides in figure S6. Figure S7 shows the average number of sides by lipid type. Here we can see that CHOL molecules have greater than 7 sides due to their bulky tail, whereas FFA molecules have less than 7 sides due to their narrow tail. While the average number of sides stays constant for CHOL as the CHOL content increases, more defects are introduced in the packing of the other lipid tails since the average number of sides deviates from 6.

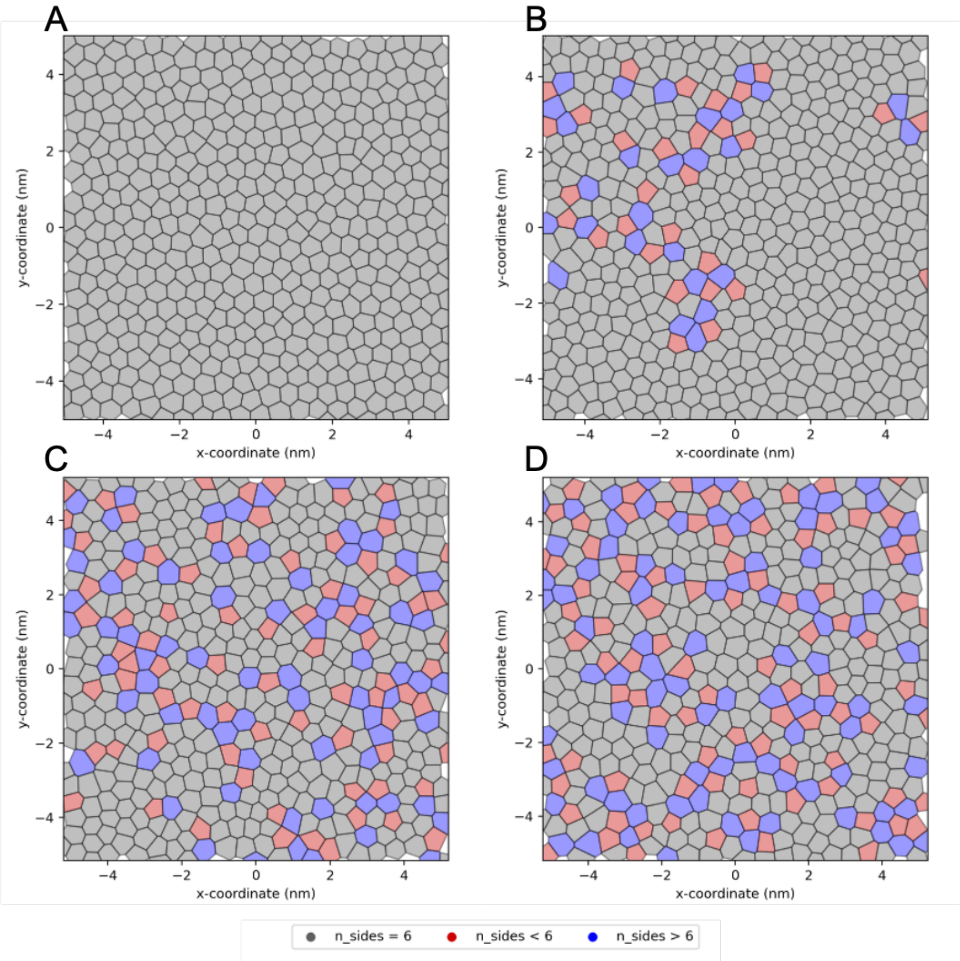


**Figure S6:** *Voronoi diagrams of an interior leaflet of a 6-leaflet CER NS C24/CHOL/FFA C24 system in a) 1:0:1, b) 1:0.2:1, c) 1:0.5:1, and d) 1:1:1 ratios. The polytopes are colored by the number of sides, where those with 6 sides are grey, less than 6 sides are red and greater than 6 sides are blue.*

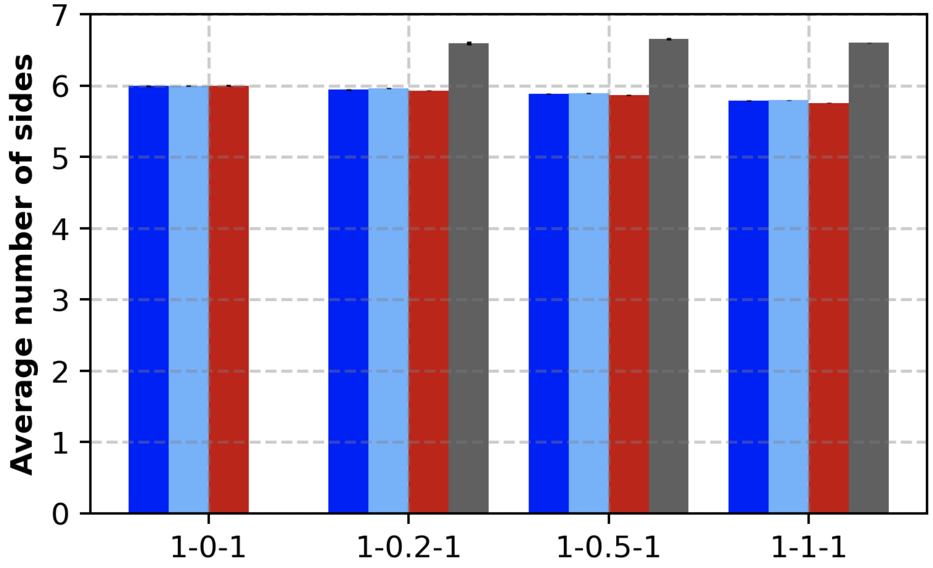


**Figure S7:** *Based on Voronoi diagrams, the average number of sides for each lipid type is plotted above, where dark blue represents the CER NS Acyl chain, light blue represents the CER NS sphingosine chain, red represents FFA, and grey represents CHOL.*

### Reverse Mapped Structures

Table S5 compares structural properties of the reverse-mapped atomistic structures to the CG self-assembled structures. Many of the small differences between the CG and reverse-mapped configurations can be explained by the significant presence of lipids stuck in the tail region of the bilayer. For example, the APT is smaller for the reverse-mapped system as the lipids stuck in the tail region are not in-plane with the other lipids, the tilt angle is higher as lipids are sitting parallel to the bilayer plane in the tail region, $S_2$ is smaller as the lipids stuck in the tail region are in alignment with the other tails, and the height is larger as the lipids in the tail region push the leaflets apart.

**Table S5:** *Mean and standard deviation of total number of hydrogen bonds in reverse-mapped atomistic structure. For the inner leaflets there are [538,264, 512] and for outer [262,136,288] [CER, CHOL, FFA24] lipids. For the interior, 187 CERs are I the extended (exCER) and 351 in the hairpin (hpCER) configurations.*

| Pairs | Inner Leaflets h-bonds | Outer Leaflets h-bonds |
|---|---|---|
| CER-CER | 618.3 ± 18.4 | 131.1 ± 9.3 |
| CER-CHOL | 143.5 ± 8.7 | 32.3 ± 4.8 |
| CER-FFA | 482.7 ± 13.1 | 78.3 ± 6.4 |
| CHOL-CHOL | 1.1 ± 1.0 | 0.4 ± 0.7 |
| CHOL-FFA | 76.4 ± 7.7 | 14.3 ± 2.9 |
| FFA-FFA | 226.3 ± 6.0 | 15.0 ± 4.7 |
| exCER-exCER | 88.9 ± 6.7 | NA |
| exCER-hpCER | 274.0 ±13.6 | NA |
| hpCER-hpCER | 255.5 ±11.2 | NA |
| exCER-CHOL | 49.4 ±6.2 | NA |
| exCER-FFA | 166.5 ±10.2 | NA |

| | | |
|---|---|---|
| hpCER-CHOL | 94.0 ±7.4 | NA |
| hpCER-FFA | 316.2 ±10.7 | NA |

**Table S6:** *Mean and standard deviation of total number of hydrogen bonds in reverse-mapped atomistic structure in the inner most two layers in order to capture only intra-layer lipid-lipid hydrogen bonding. This inner slice includes 278 CER, 139 CHOL, and 236 FFA molecules. Of the CERs, 104 are in extended conformations and 174 in hairpin.*

| **Pairs** | **h-bonds** |
|---|---|
| CER-CER | 187.2 ± 10.1 |
| CER-CHOL | 44.8 ± 4.7 |
| CER-FFA | 103.7 ± 8.8 |
| CHOL-CHOL | 0.5 ± 0.6 |
| CHOL-FFA | 18.3 ± 3.1 |
| FFA-FFA | 26.8 ± 2.8 |
